# STARNet enables spatially resolved inference of gene regulatory networks from spatial multi-omics data

**DOI:** 10.1101/2025.08.21.671434

**Authors:** Lei Hu, Shichen Zhang, Xuting Zhang, Yihai Luo, Haoteng Gu, Peng Liu, Sheng Mao, Li Chen, Yuhao Xia, Minghao Yang, Sai Zhang, Yaosen Min, Han Li, Peizhuo Wang, Hongtao Yu, Jianyang Zeng

## Abstract

Biological tissues are composed of distinct microenvironments that spatially orchestrate gene expression and cell identity. However, the regulatory principles governing domain-specific cellular functions remain poorly understood due to the lack of effective methods for mapping gene regulatory networks (GRNs) *in situ*. To address this gap, we introduce STARNet, a representation learning approach that leverages heterogeneous hypergraph modeling of spatial transcriptomic and epigenomic data to resolve tissue-domain–specific regulatory interactions. By integrating graph neural networks with contrastive learning in a self-supervised framework, STARNet learns unified embeddings that preserve both multi-modal molecular features and anatomical spatial context, enabling accurate and domain-resolved GRN reconstruction within complex tissues. Benchmarking on both simulated and real datasets demonstrates that STARNet achieves state-of-the-art performance. We further demonstrate its broad applicability across diverse biological contexts, including neural development, genetic disease risk, and drug-induced developmental toxicity. In the mouse brain, it delineates region-specific regulatory networks and reconstructs spatiotemporal programs underlying neural stem cell differentiation. In human genetics, it provides a mechanistic link between genotypes and phenotypes by showing how genome-wide association study (GWAS) variants for complex diseases perturb hippocampus-specific GRNs. In developmental toxicology, STARNet reveals that drug-induced disruptions of GRNs in defined embryonic regions underlie tissue-specific vulnerability. Collectively, STARNet offers a powerful and versatile framework for resolving the spatial regulatory logic of complex tissues, providing multi-angle insights into tissue patterning, development, and disease mechanisms.

## 1 Introduction

The formation, patterning, and homeostasis of biological tissues rely on both intercellular communication and cell–microenvironment interactions. Different tissue microdomains can impose unique regulatory codes on resident cells [1, 2]. For example, adjacent subfields in the hippocampus engage distinct regulatory programs to drive specialized neuronal functions, underscoring the need for a spatially resolved regulatory atlas that depicts domain-specific regulation and functions [3–5]. Gene regulatory networks (GRNs) link transcription factors (TFs) to their target genes and are critical for establishing cell identity and tissue functions. However, the field has lacked a computational framework capable of simultaneously learning from both spatial relationships and the interplay between chromatin accessibility and gene expression, as current methods tend to analyze these components in isolation. Consequently, the ability to delineate a truly integrated, spatially-aware landscape of regulatory interactions has remained out of reach. [6–13].

The advent of spatial multi-omics technologies, which simultaneously profile both gene expression and chromatin accessibility in the same cell *in situ*, provides an unprecedented opportunity to reconstruct spatially specific GRNs within intact tissues [14, 15]. However, due to the intrinsic noise, high dimensionality, and data heterogeneity, existing computational tools remain inadequate for analyzing such data [5].

Here, we introduce STARNet (**S**pa**T**i**A**l RNA-ATAC-seq gene **R**egulatory **Net**work), a framework designed to decipher spatially resolved GRNs by integrating spatial transcriptomic and chromatin accessibility data. Benchmarking using both simulated and real spatial datasets confirms that STARNet achieves state-of-the-art performance for inferring spatially resolved GRNs. We then demonstrate the broad utility of STARNet in several diverse downstream applications. First, we discover spatially distinct GRN modules that orchestrate region-specific cellular compositions and functions across complex tissues. Second, STARNet resolves how dynamic regulatory programs govern neural stem cell differentiation and uncovers key regulators driving lineage specification during brain development. Third, by revealing how trait-associated genome-wide association studies (GWAS) variants perturb spatial GRNs, STARNet links genotype to spatial regulatory architecture and provides insights into how genetic risk factors manifest at the tissue domain level. Finally, we illustrate its potential in therapeutic discovery by decoding spatially heterogeneous drug responses. As a unified framework, STARNet enables the elucidation of spatial regulatory principles in heterogeneous tissues. Our approach furnishes a comprehensive, multi-perspective understanding of tissue formation, developmental biology, and the molecular basis of disease.

## 2 Results

### 2.1 Overview of the STARNet framework

STARNet is a computational framework that integrates spatial transcriptomic (RNA-seq) and chromatin accessibility (ATAC-seq) data to infer spatially resolved GRNs (Fig. 1). Traditional GRN inference methods based on single modalities often yield spurious, co-expression–only links, especially under the high sparsity and noise characteristic of spatial multi-omics datasets. STARNet addresses these limitations by jointly embedding spatial spots along with their corresponding gene expression and chromatin accessibility profiles into a unified latent space. This multimodal embedding strategy simultaneously captures co-expression and chromatin accessibility-informed regulatory interactions in their native spatial context, enabling fine-grained delineation of cellular microenvironments and uncovering modality-specific molecular features that drive spatial gene regulation.

**Fig. 1.**
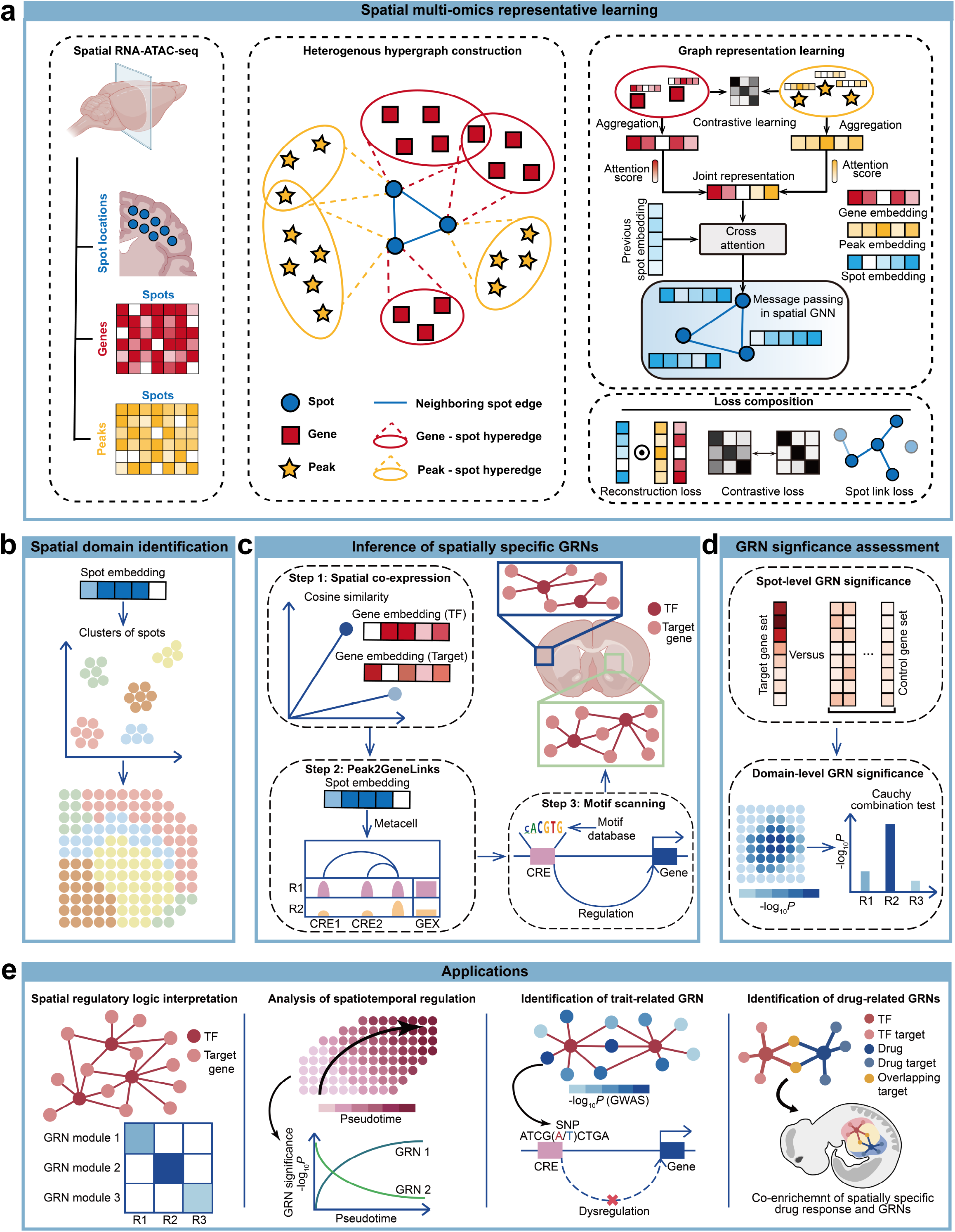
Overview of the STARNet framework. **a**, Schematic overview of the STARNet framework for representation learning from spatial multi-omics data. STARNet takes spatial RNA-ATAC-seq data as input, including spot coordinates, gene expression profiles, and chromatin accessibility peaks. It first constructs a heterogeneous hypergraph where spatial spots, genes, and peaks are represented as distinct node types. Modality-specific hyperedges link each spot to expressed genes (RNA modality) or accessible peaks (ATAC modality), while pairwise edges encode spatial adjacency between neighboring spots. A graph-based embedding strategy is then applied to generate modality-specific representations using a hybrid aggregation scheme, followed by a modality-level attention mechanism that fuses them into a unified joint embedding. This joint embedding is next combined with the previous spot embeddings via a cross-modality attention mechanism to produce updated spot representations. A spatial GNN further refines the embeddings by propagating local information through message passing over the spatial graph. The entire framework is trained end-to-end by optimizing a multi-component objective that includes reconstruction loss, cross-modal contrastive loss, and spot link prediction loss, guiding the model to learn embeddings that integrate transcriptomic and chromatin accessibility signals while preserving spatial context. **b**, Identification of spatial domains through Leiden clustering of STARNet-derived spot embeddings. **c**, Pipeline for constructing spatially specific GRNs. In Step 1, spatial co-expression is computed by correlating STARNet derived embeddings of TFs (from the CIS-BP database) with those of candidate target genes. In Step 2, spots are aggregated into metacells based on STARNet spot embeddings and candidate CREs are mapped via Peak2GeneLinks derived from chromatin accessibility versus gene expression correlations across these metacells. In Step 3, spatial co-expression and Peak2GeneLinks are integrated, and TF–target interactions are confirmed only when motif scanning detects the TF binding site in the corresponding CRE, resulting in spatially specific GRNs. **d**, Two-tiered assessment of GRN activity significance. First, a spot-level *P*-value is computed for each GRN using a permutation test on its target gene set. These spot-level *P*-values are then aggregated across spots within each spatial domain using the Cauchy combination test, yielding a domain-level significance score for each GRN. **e**, STARNet enables key downstream applications in spatial multi-omics data analyses, including the interpretation of spatially specific regulatory logic, decoding of spatiotemporal gene regulatory programs, identification of trait-associated GRNs, and analysis of spatial patterns of drug response. In this figure, abbreviations R1, R2, and R3 indicate different tissue regions, and GEX denotes gene expression. See Methods for details. The illustration of the mouse brain was created using BioRender.com.

Specifically, STARNet first constructs a unified heterogeneous hypergraph to represent the spatial multi-omics data (Fig. 1a). The graph consists of three distinct node types, including spatial spots, genes, and chromatin accessibility peaks, directly derived from the input matrices. Its topology is defined by two classes of connections. Spatial adjacency is encoded through standard pairwise edges that form a multi-scale k-nearest neighbor (k-NN) graph based on the physical coordinates of the spots (see Methods). To model different omics modalities, STARNet introduces modality-specific hyperedges, each linking a spot node to the full set of its corresponding gene or peak nodes. This structure offers a direct and biologically meaningful representation of the one-to-many relationships inherent in spatial multi-omics data.

Building on this heterogeneous hypergraph representation, STARNet learns node embeddings through a multi-stage process that progressively integrates transcriptomic, chromatin accessibility, and spatial information (Fig. 1a). First, modality-specific representations are generated using a hybrid aggregation scheme that distills biologically informative signals for each spot (see Methods). These representations are then fused via a two-level attention mechanism: a modality-level attention process estimates the relative importance of each omics modality, followed by a cross-attention step that integrates the joint representation with the previous spot embeddings. Finally, a spatial graph neural network (GNN) is applied to the spot nodes, refining their embeddings by propagating local information over the spatial adjacency graph, resulting in final node representations that capture both multi-modality features and spatial context.

To train the model, STARNet is optimized end-to-end using a composite loss function designed to impose synergistic constraints on the learned embeddings (Fig. 1a). This function is a weighted sum of three components: (1) a modality reconstruction loss that preserves fidelity to transcriptomic and chromatin accessibility profiles; (2) a cross-modality contrastive loss that aligns gene- and peak-derived representations for each spot into a coherent latent space; and (3) a spot link prediction loss that maintains neighborhood structure based on similarities in both physical and multi-omic feature space. This multi-task optimization ensures the learned node embeddings are data-consistent and biologically informative for downstream analyses.

Using the resulting embeddings, we then clustered the integrated spot representations to identify spatially coherent and biologically meaningful domains (Fig. 1b). We hypothesized that spots within each domain, sharing similar molecular profiles, correspond to distinct functional niches or anatomical regions governed by characteristic regulatory programs.

To uncover the regulatory logic underlying these spatial domains, STARNet infers spatially specific GRNs through a three-step workflow (Fig. 1c). First, it selects TFs whose expression patterns exhibit significant spatial autocorrelation and links them to putative target genes exhibiting high cosine similarity in the learned embedding space, serving as a proxy for spatial co-expression. Second, to incorporate epigenetic context, STARNet infers peak-to-gene linkages (Peak2GeneLinks) by correlating chromatin accessibility with gene expression across metacells (which are groups of spots with similar embeddings), thereby identifying candidate *cis*-regulatory elements (CREs). Third, these putative TF–target links are filtered based on sequence-level evidence: only interactions whose associated CREs contain a validated binding motif for the corresponding TF are retained. Together, these three steps ensures that the inferred GRNs represent a high-confidence set of spatially resolved regulatory interactions.

To quantify the spatial relevance of these GRNs, STARNet employs a two-stage statistical procedure (Fig. 1d). First, it tests whether the target genes of each GRN, as a collective set, exhibit significantly elevated aggregate expression within a given spot compared to a background gene set matched for overall expression levels, yielding a spot-level *P*-value. Second, these spot-level *P*-values are aggregated across spatial domains using the Cauchy combination test [16], producing a domain-level significance score for GRN activity (see Methods).

Together, these components enable critical downstream analyses in spatial multi-omics applications, including interpretation of spatially specific regulatory logic, decoding of spatiotemporal gene regulatory programs, identification of trait-associated GRNs, and characterization of spatial drug response patterns, all of which are explored in detail in the subsequent Results sections (Fig. 1e).

### 2.2 Comprehensive benchmarking confirms the accuracy and robustness of STARNet in spatial domain identification and GRN inference

We first evaluated STARNet’s performance in spatial domain identification, in comparison with eight state-of-the-art methods, including Scanpy [17], MultiVI [18], scGLUE [19], GraphST [20], STAGATE [21], Descart [22], COSMOS [23], and SpatialGlue [24]. We bench-marked these methods using simulated spatial multi-omics datasets, which were constructed by spatially distributing single-cell multi-omics data into predefined domains of distinct cell types, thus providing ground-truth for quantifying identification accuracy. Qualitative assessment of a simulated dataset demonstrated that STARNet accurately reconstructed all four spatial domains and exhibited near-perfect concordance with ground-truth annotations (Fig. 2a). In contrast, other methods exhibited critical limitations. For example, SpatialGlue produced diffuse domain boundaries. Moreover, STAGATE, scGLUE, and Scanpy failed to resolve Domain 4 under elevated noise conditions. Quantitative evaluation using multiple supervised metrics further confirmed the superior performance of STARNet in analyzing this primary simulated dataset (Fig. S1). To assess the robustness of our benchmarking, we further evaluated different approaches in analyzing four additional simulated datasets from distinct single-cell sources (see Methods) and observed consistent comparative advantage for STARNet (Fig. S2a-d). This performance advantage was also demonstrated on real human hippocampus spatial ATAC-RNA-seq data (Fig. S3a-b). STARNet uniquely delineated histologically validated anatomical regions, including the molecular layers and choroid plexus. In contrast, while other methods identified the granular cell layer, they failed to resolve additional structures, as confirmed by the Allen Brain Atlas [14]. Together, these results demonstrate the superior performance of STARNet in domain identification across both simulated and real spatial multi-omics datasets.

**Fig. 2.**
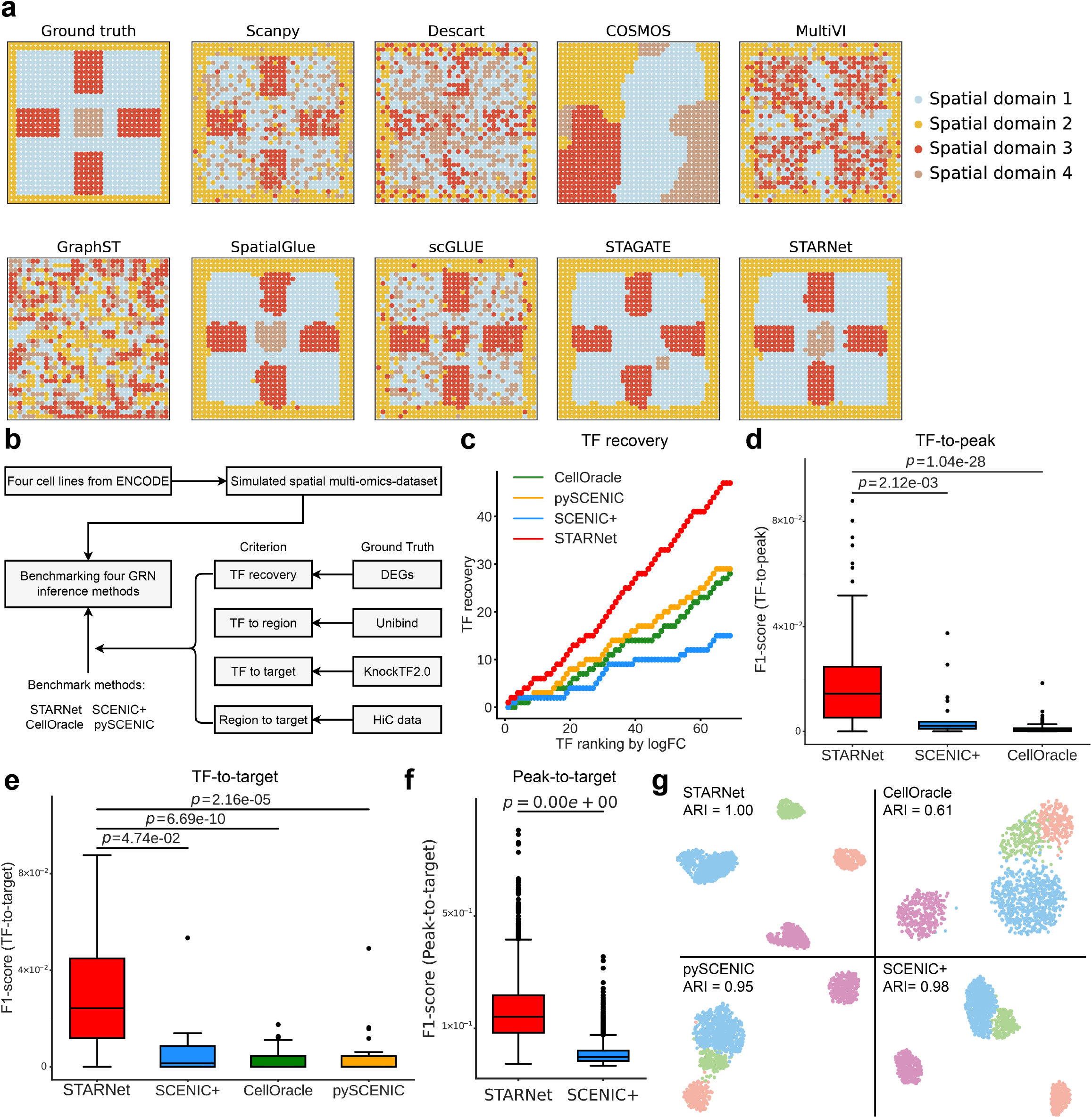
Comprehensive benchmarking of gene regulatory network inference methods on simulated spatial multi-omics data. **a**. Spatial distribution of a simulated spatial multi-omics dataset (ground-truth; first panel) and spatial domains inferred by STARNet and eight benchmark methods (other panels): Scanpy, DESCART, Cosmos, MultiVI, GraphST, SpatialGlue, scGLUE, and STAGATE. Each dot represents a spot, color-coded by its simulated domain label. **b**. Benchmarking workflow for GRN inference. We generated simulated spatial multi-omics datasets with ground-truth regulatory interactions from four deeply profiled cell lines and evaluated performance using four metrics, including TF recovery rate and association accuracies for TF-to-peak, TF-to-target, and peak-to-target. Reference benchmark data comprised: (1) spatially restricted TF expression patterns for GRN identification; (2) UniBind (a high-confidence TF-DNA interaction database) for TF-to-peak associations; (3) KnockTF2.0 (a curated database of gene expression profiles following TF knockdown or knockout) for TF-to-target associations; and (4) Hi-C data from ENCODE for peak-to-target associations (see Methods). We compared STARNet with the other three GRN inference methods: SCENIC+, CellOracle, and pySCENIC. Abbreviation: DEGs, differentially expressed genes. **c**. Cumulative TF recovery curves across methods. TFs were ranked by descending log_2_ fold-change from differential expression analysis between spatial domains. **d**. Boxplots of TF-to-peak F1 scores for each method evaluated against UniBind. *P*-values were calculated using two-sided independent t-tests (as in panels e and f). pySCENIC was excluded because it infers GRNs solely from scRNA-seq data and thus cannot perform ATAC-seq-based TF-to-peak or peak-to-gene analyses (likewise omitted from panel f). **e**. Boxplots of TF-target association F1 scores for each method, evaluated against KnockTF2.0. **f**. Boxplots of peak-to-target F1 scores for each method, evaluated against ENCODE Hi-C data. CellOracle was excluded because it generates too few peak-to-target predictions for evaluation (averaging only 1.5 peaks per target gene). **g**. UMAP projections of spot-level GRN enrichment scores, computed via AUCell based on GRNs inferred by STARNet, CellOracle, pySCENIC, and SCENIC+. Spots are colored by spatial domains, and sub-panels are annotated with ARI values. Abbreviations: ARI, adjusted rand index.

Next, to evaluate STARNet’s performance in inferring spatially specific GRNs, we conducted a comprehensive benchmark against existing GRN inference methods, originally designed for single-cell data, including SCENIC+ [10], pySCENIC [9], and CellOracle [11] (Fig. 2b). The evaluation was performed on a simulated spatial multi-omics dataset, which was rigorously generated using data from four well-characterized ENCODE (Encyclopedia of DNA Elements) [25, 26] cell lines (GM12878, MCF7, HepG2, and K562) to mimic a complex tissue architecture (see Methods; Fig. S4a). When applied to this simulated dataset, STARNet inferred a comprehensive GRN set comprising 277 TFs, each with an average of 226 predicted targets. This outperformed in both network scale and connectivity, against SCENIC+ (35 TFs; 60 targets per TF on average) and pySCENIC (226 TFs; 40 targets per TF on average). Although CellOracle identified more TFs (n = 546), its inferred GRNs were substantially sparser, with an average of only 15 targets per TF. Furthermore, spatial differential expression analysis confirmed that STARNet preferentially recovered TFs exhibiting spatially restricted expression patterns, consistent with biologically relevant regulatory activity (Fig. 2c). Together, these results establish STARNet as a powerful framework for inferring GRNs that are both comprehensive and spatially coherent.

We further evaluated the precision of STARNet’s inferred regulatory interactions through three types of validation benchmarks (see Methods). In the TF-to-peak benchmark using the UniBind database [27, 28], which consolidates uniformly processed ChIP-seq datasets, STARNet achieved a mean F1 score that was 3.3-fold higher than that of SCENIC+ (*P* = 2.12 *×* 10^*−*3^) and significantly outperformed CellOracle (*P* = 1.04 *×* 10^*−*28^; Fig. 2d). This strong performance persisted in the TF-to-target benchmark, which was conducted against the KnockTF 2.0 database [29], a resource that compiles causal evidence from TF perturbation experiments. In this evaluation, STARNet again ranked as the top-performing method, significantly outperforming all competitors (*P <* 0.05 for all comparisons; Fig. 2e). Finally, in the peak-to-target benchmark against ENCODE Hi-C data, STARNet’s accuracy was approximately 4.5-fold higher than that of SCENIC+ (*P <* 2.2 *×* 10^*−*16^), indicating its strong ability to resolve *cis*-regulatory interactions (Fig. 2f). Collectively, these benchmarks demonstrate that STARNet outperforms existing GRN inference methods in both accuracy and biological relevance.

Next, to assess whether the GRNs inferred by each method reflect spatially coherent regulatory programs, we performed unsupervised clustering of GRN enrichment scores to assess correspondence with spatial domains (see Methods). STARNet-inferred GRNs accurately identified all four spatial domains (ARI = 1.0). While pySCENIC (ARI = 0.95) and SCENIC+ (ARI = 0.98) performed well, CellOracle exhibited limited spatial coherence, failing to clearly separate anatomical domains (ARI = 0.61; Fig. 2g). Together, these results highlight STARNet’s ability to infer GRNs with superior spatial coherence and domain-level fidelity.

To assess the robustness of increasing levels of cellular mixing, we generated three additional simulated datasets in which the dominant cell type within each spatial domain accounted for 94%, 88%, and 76% of the total cells, respectively. Across all scenarios, STARNet consistently outperformed competing methods and maintained high accuracy. In contrast, the performance of other tools, most notably SCENIC+, deteriorated as cellular heterogeneity increased (Fig. S4b-d). These results highlight STARNet’s ability to deconvolve regulatory signals from complex cellular mixtures, underscoring its robustness in spatial GRN inference under realistic biological conditions.

Finally, we evaluated STARNet on a real-world benchmark: a spatial RNA-ATAC-seq dataset from the P22 mouse brain. We curated a literature-based reference set of key TFs, including Sox10, Olig2, and Foxp1, which are known to drive region-specific regulatory programs in the developing brain [30, 31]. STARNet correctly identified all reference TFs as central regulators in its inferred GRNs (Fig. S5a). In contrast, CellOracle failed to identify any of the reference TFs, while pySCENIC and SCENIC+ only partially recovered them (e.g., recovering Sox10/Sox4/Sox2 but missing others; Fig. S5b). These results confirm STARNet’s ability to recover biologically meaningful and spatially coherent GRNs from complex real-world data.

### 2.3 STARNet reveals spatially distinct GRN modules that orchestrate region-specific functions

To demonstrate the capacity of STARNet in resolving spatially specific GRNs, we applied it to spatial RNA-ATAC-seq data [14] from the mouse brain, a biologically complex system where region-specific transcriptional regulation establishes distinct cellular identities, organizes neuroanatomical structures, and drives functional specialization. Through systematic integration of spatial transcriptomic and chromatin accessibility profiles, STARNet identified 156 GRNs comprising 70,980 Peak2GeneLinks, regulating 4,285 target genes across six anatomically defined brain regions (Fig. 3a-c, and Fig. S6a). We observed strong spatial concordance between TF expression and the significance score of their corresponding GRNs which tent to be spatially enriched in regions where the TFs are highly expressed (Fig. S6b-c). This supports the validity of our inferred spatial GRNs.

**Fig. 3.**
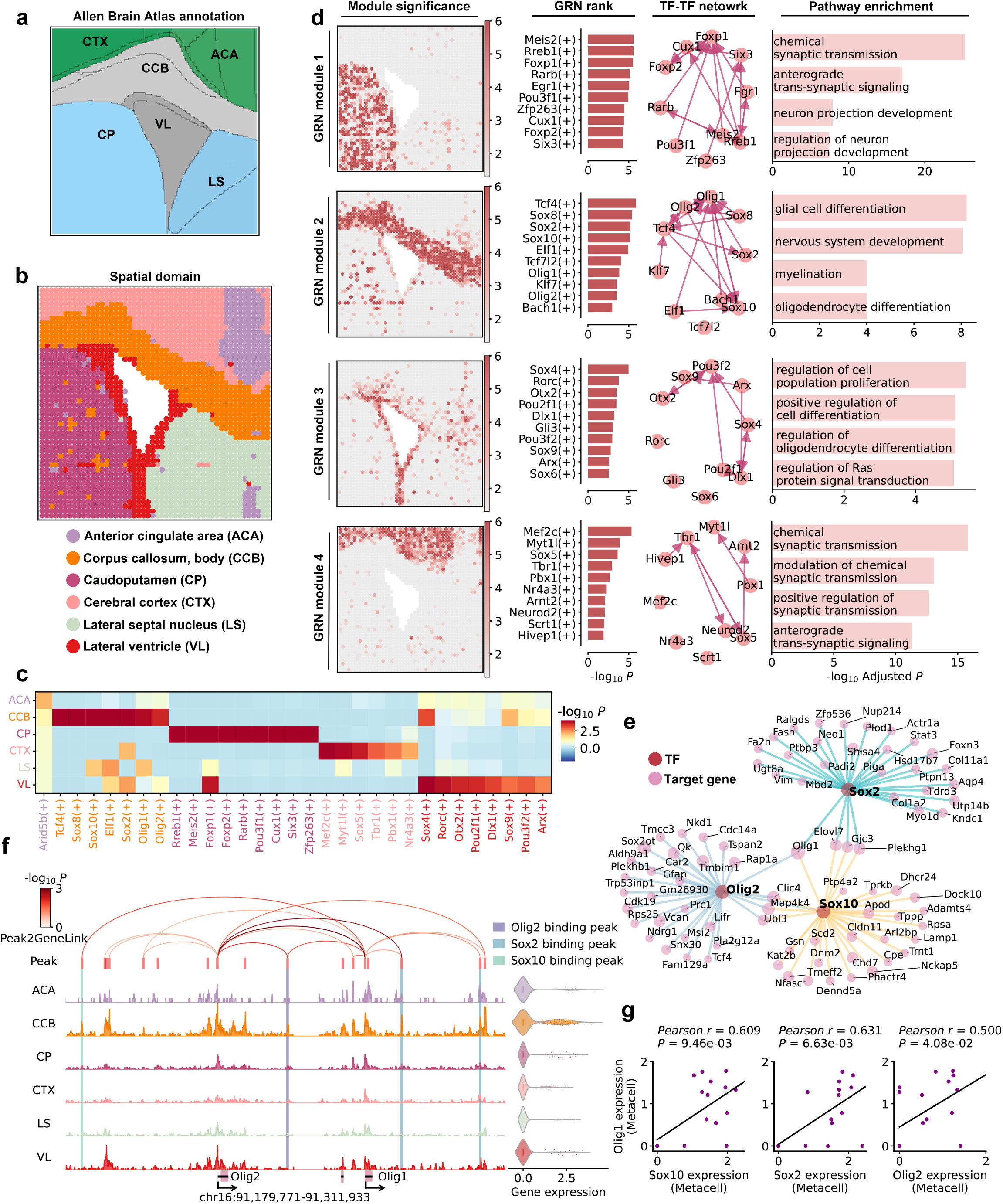
STARNet delineates domain specific gene regulatory modules in P21 mouse brain. **a**, Coronal section of the mouse brain annotated with labels from the Allen Mouse Brain Atlas. **b**, Spatial domains of a P21 mouse brain coronal section, identified by clustering spot embeddings derived by STARNet and annotated with the reference atlas. **c**, Heatmap of GRN-spatial domain associations derived from the Cauchy combination test. Heatmap intensity reflects the statistical significance of each association, with rows representing spatial domains and columns corresponding to GRNs. Column label colors indicate the anatomical regions where each GRN is enriched. The (+) suffix denotes GRN of the corresponding TF. **d**, Top four GRN modules identified from the P21 mouse brain section, which were derived by applying the Leiden clustering algorithm to spatially significant GRN patterns (see Methods). Each module is represented by four panels: (1) A spatial map showing the module significance scores, i.e., *P*-values obtained from the permutation tests involving all TFs and target genes within the module. The color scale minimum is set to *−* log_10_ 0.05; (2) A horizontal bar plot displaying the aggregated *P*-values over the corresponding domain for the top 10 GRNs in each module, calculated using the Cauchy combination test. (3) A TF-TF interaction network, with directional arrows indicating regulatory relationships; and (4) Gene Ontology (GO) term enrichment analysis for TFs and target genes associated with each module (*P*-values determined by the hypergeometric test). **e**, Visualization of the GRNs formed by TFs Sox10, Olig2 and Sox2. Nodes represent both TFs (red) and their top 30 target genes (pink) ranked by their regulatory scores (see Methods). Node size indicates the regulatory score of each target gene, and edge color distinguishes individual regulatory relationships corresponding to different TFs. **f**, Chromatin accessibility profiles of genomic regions spanning the Olig1 locus across different spatial domains. The Peak2GeneLinks are visualized as arcs, color-coded by the statistical significance of their corresponding correlations. Peak2GeneLinks were inferred based on correlations between chromatin accessibility peaks and gene expression across the metacells of all spatial spots (see Methods). The binding peaks of Sox10, Olig2, and Sox2, identified using the SnapATAC2 toolkit, are highlighted with a semi-transparent box. A violin plot on the right shows the normalized expression levels of Olig1 across different spatial domains. **g**, Scatter plots displaying the correlations between Olig1 expression and that of Sox10, Sox2 and Olig2. Each data point represents a metacell. *P*-values are calculated using the two-sided t-test.

Tissue functions are often orchestrated by the coordinated activity of multiple GRNs with spatial specificity. To systematically characterize their spatial organization, we grouped the GRNs inferred by STARNet into 11 modules based on the spatial patterns of their significance scores (Fig. 3c, and Fig. S7a). That is, GRNs exhibiting similar spatial distributions of significance scores were assigned to the same module (see Methods). Among these 11 modules, the top four exhibited significant spatial enrichment (Cauchy combination test, *P <* 0.05) and contained multiple spatially significant GRNs (Fig. 3d, and Fig. S7). Further analysis revealed that each of these four modules comprises distinct GRNs, unique TF-TF regulatory networks (i.e., regulatory interactions among TFs within each module), and is associated with distinct biological functions (Fig. 3d).

Specifically, GRN Module 1, enriched in the caudateputamen (CP) region, is associated with chemical synaptic transmission and neuronal projection development, and includes key TFs such as Foxp1, Rreb1, and Meis2, which are critical for the function of medium spiny neurons. GRN Module 2, localized in the corpus callosum, body (CCB) region, regulates glial and oligodendrocyte differentiation, with Sox2, Olig1, and Olig2 acting as core regulators. GRN Module 3, active in the lateral ventricle (VL) region, governs cell proliferation and differentiation through TFs such as Dlx1 and Otx2, both known to regulate neural progenitor cells. GRN Module 4, predominantly active in the cerebral cortex (CTX) region, contributes to synaptic plasticity and cortical neuron maturation via Mef2c, Myt1l, and Neurod2 [30, 31] (Fig. 3d).

Further analysis of the transcriptional regulatory networks within GRN modules revealed that multiple TFs cooperatively regulate specific subsets of target genes. Notably, Sox10, Sox2, and Olig2 act as key regulators of oligodendrocyte lineage commitment, thereby influencing myelin formation and neural signal transduction in the CCB region [32, 33]. STARNet showed that these TFs co-regulate Olig1 (Fig. 3e) by binding to multiple spatially specific enhancers near its promoter and jointly controlling its expression (Fig. 3f-g and Fig. S8). Previous studies have demonstrated that Olig1 is a critical regulator of oligodendrocyte maturation and myelination [33]. Thus, STARNet provided mechanistic evidence supporting this regulatory logic.

As an additional case, we examined Rgs9, a gene implicated in dopamine signaling, which is primarily expressed in the basal ganglia, including the CP region [34]. STARNet identified TFs, including Rreb1, Meis2, and Foxp1, which are essential for medium spiny neurons, as regulators of Rgs9 expression, which are important for the maintenance of dopamine signaling homeostasis (Fig. S9a-c). Mechanistically, these TFs co-bind to spatially specific CREs near the Rgs9 promoter to coordinately upregulate its expression (Fig. S9d).

In summary, STARNet systematically identifies key spatially specific regulators and their associated regulatory networks across distinct spatial domains. These regulators exhibit pre-dominant expression in regionally restricted cell types, highlighting their cell-type specificity. Furthermore, STARNet enables comprehensive characterization of spatial gene regulatory relationships across multiple GRNs, revealing how distinct networks cooperatively orchestrate cellular functions within spatially defined tissue regions.

### 2.4 STARNet enables dissection of spatiotemporal gene regulatory programs during neural stem cell-to-oligodendrocyte differentiation with STARNet

Neural stem cells (NSCs) in the subventricular zone (SVZ) give rise to oligodendrocytes from the early postnatal period into adulthood, although their output declines with age [35–38]. These newly formed oligodendrocytes migrate along white-matter tracts, particularly the corpus callosum, where they integrate and form functional myelin sheaths [39, 40]. While single-cell transcriptomic studies have already delineated the gene expression programs underlying this process [36, 41], the precise spatiotemporal orchestration of TFs, their binding sites, and downstream target genes that drive oligodendrocyte differentiation, migration, and maturation remains poorly understood. We hypothesize that STARNet can resolve spatiotemporal regulatory dynamics and uncover key gene regulatory networks that orchestrate oligodendrocyte lineage progression.

To test this hypothesis, we applied STARNet to a spatial multi-omics dataset of P22 mouse brain [14]. The inferred spatial clusters exhibited strong concordance with anatomical regions annotated in the Allen Brain Atlas (Fig. 4a-b). In total, STARNet identified 141 spatially specific GRNs and successfully recovered well-established regional regulators, including Sox10, Sox9, Sox2, and Olig1 for oligodendrocytes in the genu of the corpus callosum and anterior commissure (CCG/ACO); Sox4, Pax6, and Dlx1 for NSCs in the VL, which encompasses the SVZ; Mef2c and Neurod2 for neurons in the cerebral cortex layers (L1-L4 and L5-L6); and Meis2, Rreb1, and Foxp1 for medium spiny neurons in the CP [30, 31] (Fig. 4a-b and Fig. S10-S11). We further observed strong spatial concordance between regional expression patterns of key domain-specific regulators, such as Dlx1, Mef2c, Meis2, and Olig1, and spatial significance maps of their respective GRNs [30, 31] (Fig. S10b-c). GO enrichment analysis further validated that the target genes of these TFs were significantly associated with biological functions characteristic of their respective anatomical domains (Fig. S10d). These results establish STARNet’s capacity to decode spatial gene regulation as a foundation for exploring its dynamic changes along developmental trajectories.

**Fig. 4.**
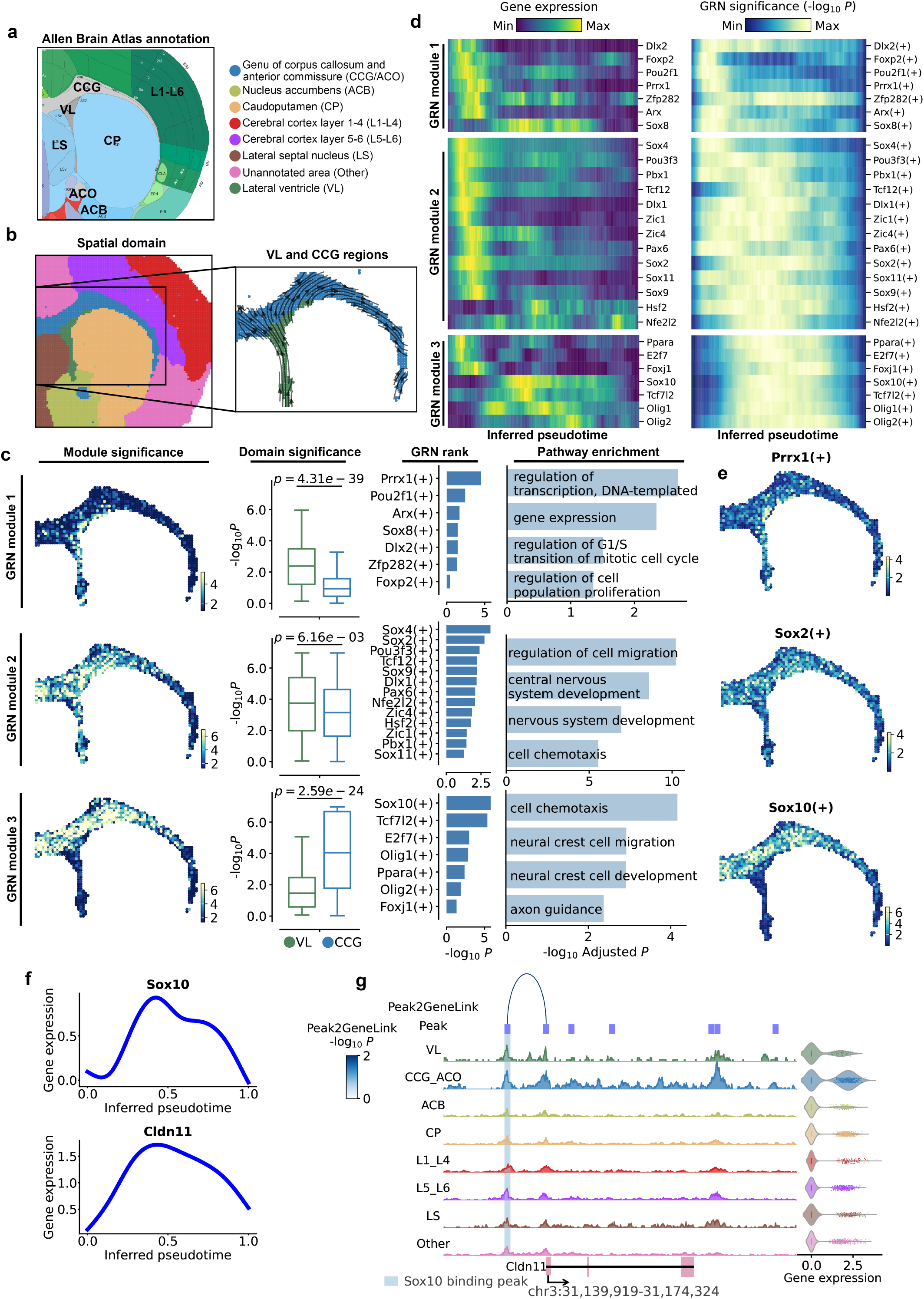
Identification of spatiotemporal regulatory patterns from STARNetinferred GRNs in P22 mouse brain. **a**, Annotated regional reference from the Allen Mouse Brain Atlas for a P22 mouse brain coronal section. **b**, Spatial domains of a coronal section of P22 mouse brain, clustered using the STARNet-derived spot embeddings and annotated with the reference atlas (on the left). Inset shows SpaTrack-derived velocity streamlines in the VL and CCG, with enlarged regions (on the right) illustrating dynamic cellular migration trajectories. SpaTrack was used to infer velocity streamlines, through calculating the transition probabilities and directions among neighboring cells (see Methods). **c**, Analysis of the three GRN modules enriched in the VL and CCG regions, which were derived using the Leiden clustering algorithm on the 27 spatially significant GRNs derived by STARNet (*P <* 0.05, Cauchy combination test). Each module is illustrated with four panels: (1) A spatial map showing the module significance scores, in which *P*-values were determined by the permutation tests involving all TFs and target genes within the module, and the color scale minimum was set to *−* log_10_ 0.05; (2) A boxplot displaying the regional significance distribution of GRN modules in VL and CCG regions; (3) A horizontal bar plot of the aggregated *P*-values for GRNs in each module, calculated using the Cauchy combination test; and (4) GO term enrichment analysis for TFs and target genes in each module, with *P*-values calculated by the hypergeometric test. Module 1 shows predominant enrichment in VL, module 2 spans both VL and CCG, while module 3 is chiefly enriched in CCG. **d**, Dynamics of gene expression (left panel) and GRN significance (right panel) along a pseudotime trajectory spanning from VL to CCG. Pseudotime values were inferred using SpaTrack (see Methods). **e**, Spatial significance maps of the GRNs involving Prrx1, Sox2, and Sox10 in modules 1, 2, and 3, respectively. Colors represent *−* log_10_ *P*. **f**, Pseudotemporal expression dynamics of Sox10 andits target Cldn11 with smoothing curves fitted by Generalized Additive Models (GAMs). **g**, Chromatin accessibility profiles of genomic regions spanning the Cldn11 locus across different spatial domains. The Peak2GeneLink is visualized as an arc, color-coded by the statistical significance of the correlation. The linkage was inferred based on the correlation between chromatin accessibility peaks and gene expression across metacells derived from all spatial spots (see Methods). The binding peak of Sox10, identified using the SnapATAC2 toolkit, is highlighted with a semi-transparent box. A violin plot on the right displays the normalized expression levels of Cldn11 across spatial domains. Abbreviations: ACB, nucleus accumbens; CCG/ACO, genu of corpus callosum and anterior commissure; CP, caudoputamen; LS, lateral septal nucleus; L1-L4, cerebral cortex layers 1-4; L5-L6, cerebral cortex layers 5-6; VL, lateral ventricle.

To further investigate how transcriptional regulation shapes spatial transitions in cellular states, we integrated STARNet-inferred GRNs with SpaTrack, a computational framework based on optimal transport theory that reconstructs developmental trajectories from spatial transcriptomic data [42]. This integration enabled a mechanistic linkage between region-specific regulatory programs and the spatial progression of cellular states. SpaTrack-derived pseudotemporal ordering revealed a global developmental hierarchy across anatomical domains (Fig. S12a-c). More specifically, the VL, encompassing the SVZ where NSCs reside, exhibited the earliest pseudotime and the highest transcriptional entropy. In contrast, the CCG/ACO, characterized by a high abundance of oligodendrocytes, displayed intermediate pseudotime and entropy, while terminally differentiated neurons in the cerebral cortex (L1-L4 and L5-L6) showed the latest pseudotime and the lowest entropy. These findings uncovered through STARNet analysis, reveal a spatially ordered differentiation continuum shaped by region-specific regulatory programs.

Next, to dissect the transition from NSCs in the VL to oligodendrocytes in the CCG, we reconstructed a spatiaotemporal trajectory representing this differentiation pathway (Fig. 4b). Along this axis, STARNet identified 27 statistically significant spatial GRNs (*P <* 0.05, Cauchy combination test). Clustering these GRNs based on spatial significance profiles revealed three regulatory modules (Fig. 4c-d). Module 1 exhibited strong activity in the VL (*P* = 4.31 *×* 10^*−*39^) and included canonical NSC-related TFs such as Prrx1 and Dlx2 [43, 44], with target genes enriched for cell proliferation and cell cycle-related pathways (Fig. 4d-e). Module 2 was active in both VL and CCG, with slightly higher activity in the VL (*P* = 6.16 *×* 10^*−*3^), and comprised key lineage-determining TFs including Dlx1 (GABAergic fate [45]), Sox2 (neurogenesis [46]), and Foxj1 (ependymal differentiation [47]), suggesting its role in NSC maintenance and multilineage priming. Module 3 showed specific activation in the CCG (*P* = 2.59 *×* 10^*−*24^), with target genes significantly enriched in axon guidance and gliogenesis-related pathways, and regulated by oligodendrocyte-associated TFs, such as Sox10, Olig2, and Tcf7l2 [48]. Notably, Tcf7l2 has been reported as a key regulator of oligodendrocyte differentiation, and its loss leads to defective maturation [49]. Additionally, within Module 2, a temporal lag was observed between TF expression and downstream GRN activation (Fig. 4d), potentially reflecting the time required for TF accumulation to initiate effective gene regulation. Taken together, these findings highlight the power of STARNet in resolving how transcriptional programs are temporally staged and spatially organized during NSC-to-oligodendrocyte lineage progression along the VL-CCG axis.

To further assess STARNet’s capacity for inferring spatiotemporal gene regulation, we examined Cldn11, a driver of myelin sheath formation and maintenance [50] as a representative gene. A spatiotemporal correlation was observed between the expression of Cldn11 and Sox10, a master transcriptional regulator of oligodendrocytes [41] (Fig. 4f and Fig. S12d-e). Focusing on the Cldn11 locus, STARNet identified CCG-specific chromatin accessibility patterns, including a high-confidence Peak2GeneLink between an accessible CRE and Cldn11, consistent with Sox10 binding (Fig. 4g). These chromatin features support enhancer-mediated regulation of Cldn11 in a spatially specific manner, thereby illustrating STARNet’s ability to decode *cis*-regulatory logic driving lineage-specific gene activation.

As an additional case study, we found that Sox4 likely acts as an upstream regulator of Dlx1, a TF essential for neural lineage specification (Fig. S13a). This regulatory relationship is supported by previous studies [51], and further corroborated by STARNet. Notably, Sox4 has been reported to suppress oligodendrocyte differentiation [52]. Both Sox4 and Dlx1 exhibited high expression in the VL but were downregulated in the CCG, suggesting suppression of neurogenic regulatory programs during oligodendrocyte lineage commitment (Fig. S13b-c). Chromatin accessibility at the Dlx1 locus further supported this regulatory interaction (Fig. S13d), suggesting that suppression of neurogenic programs is a key regulatory step during oligodendrocyte lineage commitment.

In conclusion, by integrating STARNet with spatial trajectory inference, we leverage its ability to resolve GRN activity across space and time that governs the progression of the oligodendrocyte lineage from NSCs within anatomically defined brain regions.

### 2.5 STARNet enables spatially resolved dissection of the genetic regulation of complex traits

GWAS has been instrumental in identifying genetic variants linked to diverse phenotypes and in elucidating the genetic basis of complex traits. However, how these variants remodel spatial domain-specific regulatory networks to influence complex traits remains largely unresolved. To address this, we developed an integrative pipeline that maps GWAS summary statistics onto spatial domain-specific GRNs inferred by STARNet (Fig. 5a). Briefly, we projected GWAS associations onto spatial regions using gsMap [53], applied the Cauchy combination test to identify trait-associated GRNs, intersected significant single nucleotide polymorphisms (SNPs) with TF-binding peaks to establish SNP-TF-target gene relationships, and assigned each gene the *P*-value of its most significant SNP (see Methods).

**Fig. 5.**
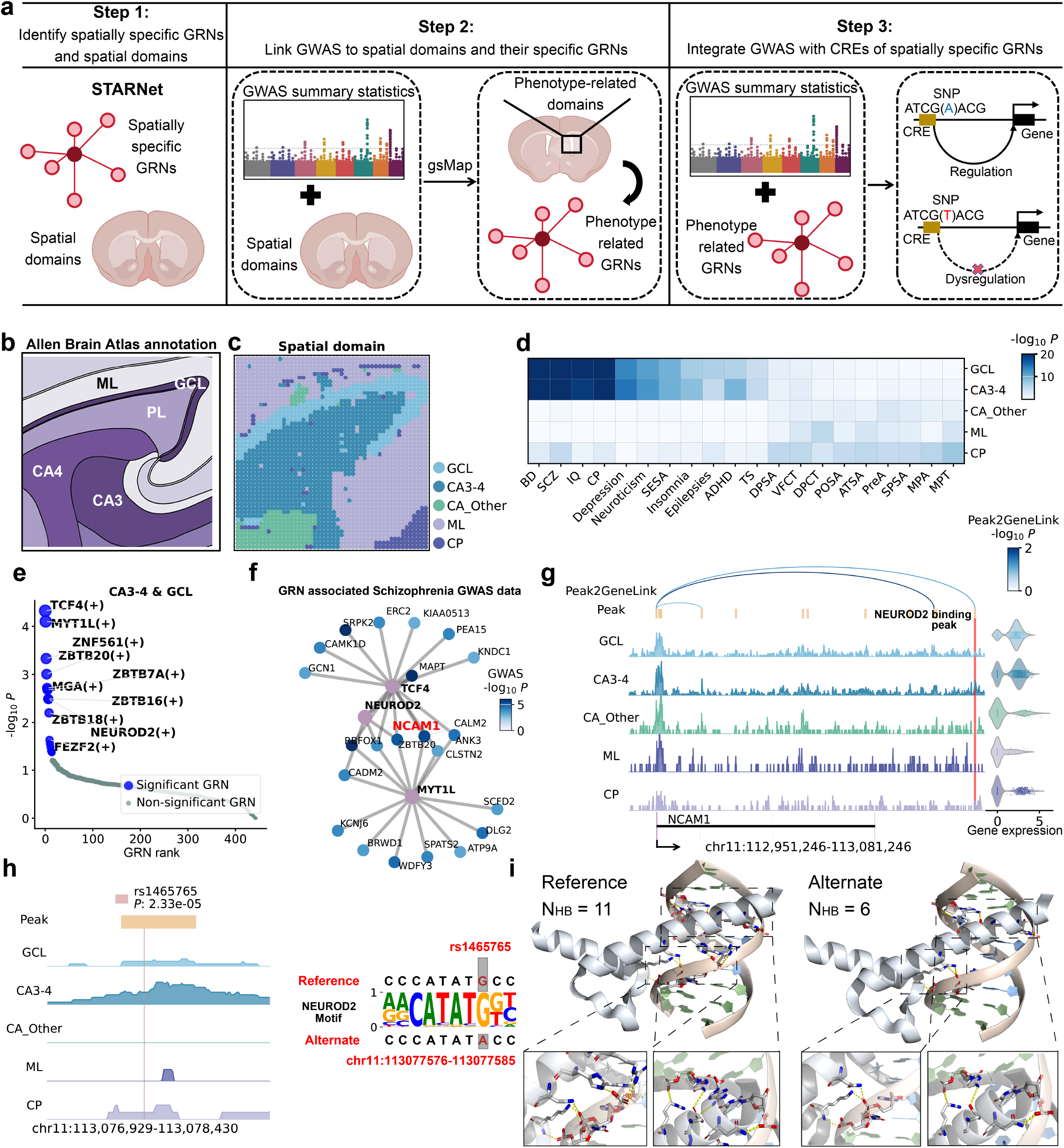
Mapping GWAS traits onto the STARNet-inferred spatially specific regulatory landscape of human hippocampus. **a**, Pipeline for linking GWAS associations to spatially specific GRNs inferred by STARNet. First, STARNet is employed to identify spatial domains and their domain-specific GRNs. Next, gsMap is applied to map GWAS associations onto these domains, highlighting trait-associated GRNs. Finally, candidate regulatory variants are identified by intersecting GWAS SNPs with CREs harboring TF-binding motifs (see Methods). **b**, Annotated regional reference from the Allen Human Brain Atlas for a human hippocampal section. **c**, Spatial domains of a human hippocampal section, determined by clustering the STARNet-generated spot embeddings and annotated with the Allen Human Brain Atlas. **d**, Heatmap showing associations between spatial domains and traits, derived from the Cauchy combination test and gsMap. Rows denote spatial domains and columns denote traits. Colors indicate association significance as *−* log_10_ *P*. Abbreviations: BD, bipolar disorder; SCZ, schizophrenia; IQ, intelligence quotient; CP, cognitive performance; SESA, sensitivity to environmental stress and adversity; ADHD, attention-deficit hyperactivity disorder; TS, tourette syndrome; DPSA, dorsolateral prefrontal surface area; VFCT, ventral frontal cortex thickness; DPCT, dorsolateral prefrontal cortex thickness; POSA, pars opercularis surface area; ATSA, anteromedial temporal surface area; PSA, precuneus surface area; SPSA, superior parietal surface area; MPA, motor premotor area; MPT, medial prefrontal thickness. **e**, Dot plot showing aggregated P-values (calculated via the Cauchy combination test) for all GRNs across individual spots in the GCL and CA3-4 regions. GRNs with *P <* 0.05 are designated significant, and the top ten significant GRNs are labeled. **f**, Visualization of GRNs regulated by selected TFs (TCF4, NEUROD2, and MYT1L, whose node colors are shown in purple). Only target genes harboring SNPs with *P <* 0.01 are shown. Node colors indicate the *P*-values of the associated SNPs. NCAM1 is highlighted in red for downstream analysis. **g**, Chromatin accessibility profiles of the genomic regions covering NCAM1 across different spatial domains. Peak2GeneLinks are visualized as arches, color-coded by the statistical significance of their corresponding correlations. Peak2GeneLinks were inferred based on correlations between chromatin accessibility peaks and gene expression across metacells from all spatial spots (see Methods). The NEUROD2 binding peak, identified using the SnapATAC2 toolkit and overlapping the significant SNP rs1465765, is marked by a red box. Violin plot (right) shows normalized NCAM1 expression across spatial domains. **h**, Chromatin accessibility profiles across spatial domains for the genomic region encompassing the predicted NEUROD2 binding peak (highlighted in yellow). The SNP rs1465765 is indicated by a semi-transparent vertical bar. In the right panel, the NEUROD2 motif logo shows both reference and alternative alleles, with the mutation site marked by a gray box. **i**, Molecular visualization of interactions between the DNA-binding domain of NEUROD2 and its target DNA, as predicted by AlphaFold3. Left: NEUROD2 bound to the reference motif sequence; right: NEUROD2 bound to the alternative motif sequence. Hydrogen bonds between specific amino acid residues and DNA bases are shown as yellow dashed lines. Eleven hydrogen bonds are observed in the reference complex (left), compared to six in the alternative (right). Abbreviations: CA3-4, cornu ammonis 3-4; ML, molecular layer; GCL, granule cell layer; CP, choroid plexus; CA Other, remaining cornu ammonis regions.

To illustrate our pipeline’s ability to resolve spatially organized regulatory landscapes underlying complex traits, we applied STARNet to a spatial multi-omics dataset of human hippocampus comprising 2,500 transcriptomic spots [14]. Clustering of STARNet-derived spot embeddings (see Methods) revealed spatially coherent domains that matched known anatomical subregions of the hippocampus (Fig. 5b-c), providing a robust basis for down-stream GWAS integration. We then used gsMap [53] to analyze GWAS summary statistics for 20 human complex traits [54–66], identifying significant associations between specific traits and spatially distinct hippocampal regions (Fig. 5d). For example, genetic signals associated with intelligence quotient (IQ), cognitive performance (CP), and schizophrenia (SCZ) were enriched in the granule cell layer (GCL) and cornu ammonis 3-4 (CA3-4) regions. These findings validated the spatial context and biological relevance of our approach in mapping complex traits onto hippocampal subregions.

Focusing on SCZ as a representative trait, we further explored its spatially specific regulatory landscape by identifying significant associations between SCZ and hippocampal subregions. Notably, CA3-4 and GCL exhibited the most significant spatial association signal of SCZ (Fig. S14a). We then inferred domain-specific GRNs using STARNet and prioritized those predominantly active in SCZ-enriched regions (Fig. 5e and Fig. S14b). This analysis highlighted TCF4, MYT1L, and NEUROD2 as key TFs enriched in SCZ-associated regions, with their activity patterns generally aligning with those domains (Fig. S14a, c-e). Importantly, these TFs have been previously established as important mediators of hippocampal neurogenesis and higher-order cognitive functions [67–69].

To elucidate the downstream regulatory landscape of these key TFs at the molecular level, we used STARNet to derive their corresponding GRNs, identifying 210 putative target genes. Among them, 22 genes harbored significant SNPs (*P <* 0.01) that overlapped with regulatory peaks (Fig. 5f). These regulatory peaks were defined as putative CREs bearing TF-binding motifs and linked to target genes via Peak2GeneLinks (see Methods). We further examined the *cis*-regulatory circuitry of a representative target gene, NCAM1, a known risk gene for intellectual disabilities [70]. We found that all these key TFs regulate NCAM1 and identified a regulatory peak near the NCAM1 promoter, spatially enriched in CA3-4 and GCL regions, harboring a significant SNP (*P* = 2.33 *×* 10^*−*5^) (Fig. 5f-g). This SNP lies within a predicted NEUROD2-binding motif, supporting a potential spatial regulatory mechanism by which NEUROD2 modulates NCAM1 expression (Fig. 5g-h).

To mechanistically assess whether rs1465765 affects NEUROD2 binding affinity, we further performed structural modeling of the corresponding DNA–protein complex using AlphaFold3 [71] (Fig. 5i). High-confidence binding models were obtained for both reference and alternate DNA sequences, with interfacial predicted TM (ipTM) scores 0.87 and 0.88, respectively.

Based on the predicted structures, we further evaluated the corresponding binding affinities using PDBePISA [72], a tool for detailed analyses of macromolecular interfaces. The results revealed a notable reduction in NEUROD2-DNA binding from the reference to the alternate allele. Specifically, the number of hydrogen bonds decreased from 11 to 6. Focusing on the core binding interface between NEUROD2 and the 3’-5’ DNA strand, we observed a Δ^*i*^*G* (free energy change upon docking) shift from -12.8 kcal/mol to -11.4 kcal/mol, and the interface area decreased from 428.0 Å^2^ to 395.9 Å^2^. These structural changes indicate reduced NEUROD2 binding affinity, which may lead to altered NCAM1 transcription. Collectively, these results provide a mechanistic link between rs1465765 and its spatially resolved regulatory effect on gene expression.

To evaluate the broader applicability of our framework to additional complex traits, we also analyzed GWAS summary statistics for cognitive performance. Significant spatial associations were observed in the CA3-4 (*P* = 2.88 *×* 10^*−*23^) and GCL (*P* = 3.16 *×* 10^*−*23^) regions (Fig. S15a-b). Within the regulatory network governed by TCF4, MYT1L, and NEUROD2, three TFs that are predominantly active in the GCL and CA3-4 domains, STARNet identified 148 target genes with regulatory peaks overlapping cognitive performance-associated variants, among which 23 were linked to SNPs reaching statistical significance (*P <* 0.01) (Fig. S15c).

Notably, two variants, including rs590745 (*P* = 1.74*×*10^*−*5^) and rs12616299 (*P* = 5.55*×*10^*−*6^), were predicted to modulate the transcription of SCN2A, a previously reported biomarker for cognitive impairment [73], through spatially constrained CREs (Fig. S15d-f). These findings point to a potential mechanism by which cognitive performance-associated SNPs may influence SCN2A expression by modulating spatially restricted regulatory activity.

In summary, our STARNet-based pipeline integrates genetic associations with spatially specific regulatory networks, offering a mechanistic framework for linking genetic variants to spatially resolved transcriptional programs underlying complex traits.

### 2.6 STARNet-based GRN inference enables mechanistic understanding of spatially resolved drug response patterns

Drug responses in complex tissues are highly dependent on spatial context, reflecting distinct local regulatory programs that modulate drug efficacy and resistance. Although various computational methods have been developed to study drug-gene interactions or predict drug responses using bulk or single-cell data [74–76], it remains challenging to elucidate how region-specific GRNs influence spatial drug responses (SDRs) within anatomically defined domains. To address this gap, we developed a pipeline that integrates spatial mapping of drug-target interactions with STARNet-inferred GRNs to systematically identify and prioritize co-enriched drug-GRN associations that may underlie domain-specific drug responses (Fig. 6a). Specifically, we prioritized drug-GRN pairs whose drug targets overlapped with TF-regulatory genes from STARNet-inferred GRNs and assessed potential associations by analyzing their proximity within the protein-protein interaction (PPI) network (see Methods).

**Fig. 6.**
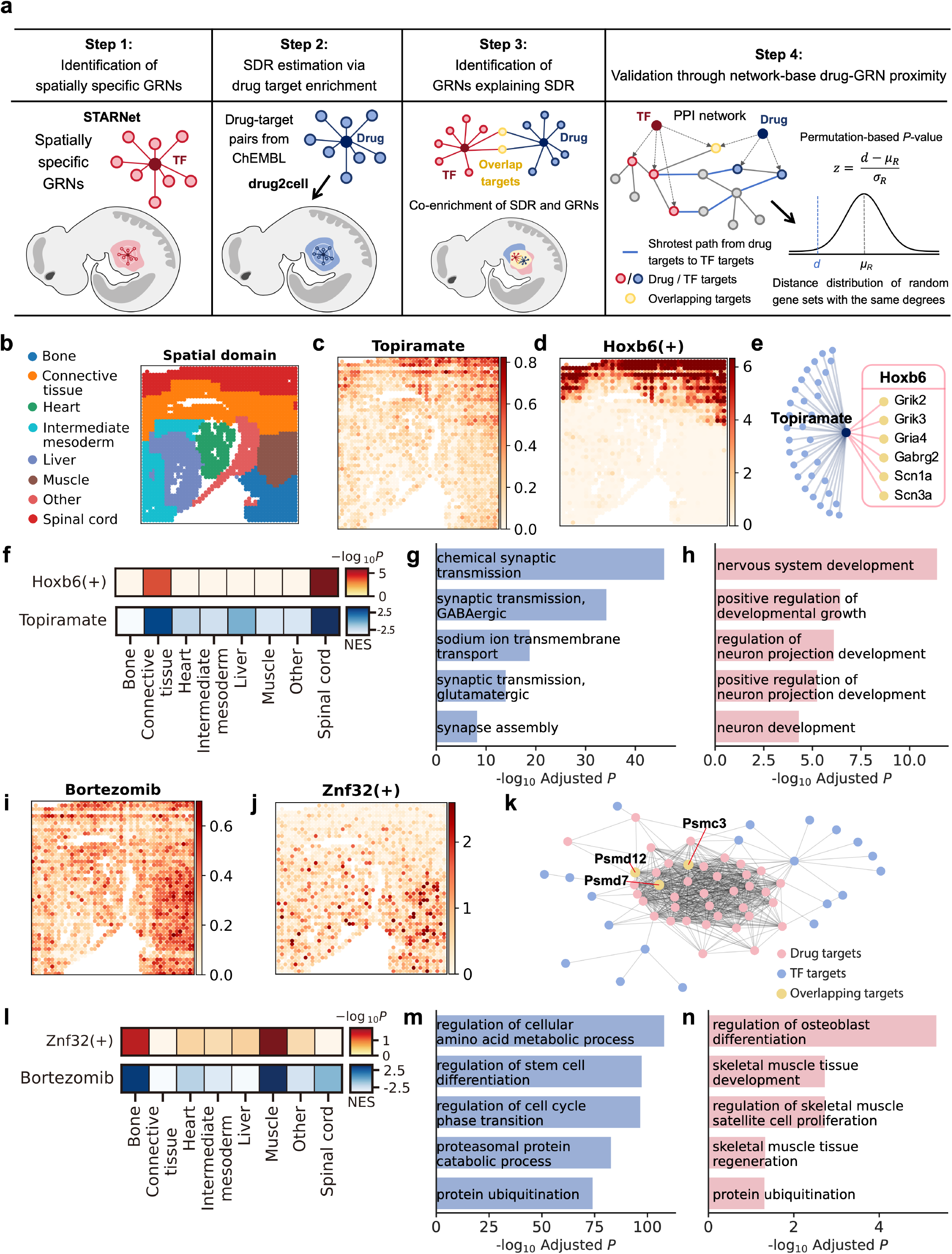
Spatial mapping of drug targets onto STARNet-inferred GRNs reveals domain-specific drug response mechanisms in E13 mouse embryo. **a**, Overview of the STARNet-based analysis pipeline for integrating spatial GRN inference with drug response mapping: (1) Predicting spatially specific GRNs using STARNet; (2) Mapping drug targets from ChEMBL to spatial domains and assess their enrichment using drug2cell; (3) Identifying drug targets that are also TF-regulated genes from STARNet-inferred GRNs within each spatial region; (4) Validating drug-GRN associations by quantifying the network proximity between drug targets and TF targets within a PPI network. **b**, Annotation of spatial domains of an E13 mouse embryo section, clustered using the spot embeddings generated by STARNet. **c**, Spatial pattern of the mean expression of topiramate target genes on the E13 mouse embryo section. **d**, Spatial pattern of significance scores for the Hoxb6(+) GRN on the E13 mouse embryo section, with colors indicating *−* log_10_ *P* scores. **e**, Network visualization of topiramate targets (from ChEMBL) and Hoxb6 targets (from GRNs inferred by STARNet), highlighting shared nodes (blue) and distinct nodes (yellow). **f**, Heatmap of normalized enrichment scores for topiramate targets and normalized significance scores for the Hoxb6(+) GRN across spatial domains of the mouse embryo. Higher values indicate stronger spatial enrichment. **g-h**, GO term enrichment analyses for topiramate and Hoxb6 targets, respectively, showing significantly enriched biological processes. **i**, Spatial pattern of the mean expression of bortezomib target genes on the E13 mouse embryo section. **j**, Spatial pattern of significance scores for the Znf32(+) GRN on the E13 mouse embryo section, with colors indicating *−* log_10_ *P* scores. **k**, Network visualization of bortezomib targets (from ChEMBL) and Znf32 targets (from GRNs inferred by STARNet) in the PPI network. Only connected nodes are displayed. **l**, Heatmap of normalized enrichment scores for bortezomib targets and normalized significance scores for the Znf32(+) GRN across spatial domains of the mouse embryo. **m-n**, GO term enrichment analyses for bortezomib and Znf32 targets, respectively, showing significantly enriched biological processes. Abbreviations: NES, normalized enrichment score.

We applied this pipeline to an E13 mouse embryo dataset [15] with annotated labels (Fig. S16). We first uncovered significant spatial overlap between the targets of the antiepileptic drug topiramate [77] and the Hoxb6-regulated genes from STARNet-derived GRNs in the developing spinal cord (Fig. 6b-d, f). Six genes, including three glutamate receptor subunits (Grik2, Grik3, Gria4), one GABAA receptor subunit (Gabrg2), and two sodium channel genes (Scn1a, Scn3a), were identified as both topiramate targets and Hoxb6-regulated genes (Fig. 6e). All are highly enriched in the spinal cord and encode critical ligand-gated ion channels and voltage-gated sodium channels required for neuronal excitability and synaptic signaling. PPI network analysis revealed close interconnectivity between topiramate targets and Hoxb6-regulated genes (Z = -7.9, *P <* 0.001), suggesting that topiramate may perturb the Hoxb6-mediated GRN in this region. GO enrichment analysis revealed that both topiramate targets and Hoxb6-regulated genes are commonly associated with nervous system-related processes, including synaptic transmission, ion channel activity, and nervous system development (Fig. 6g-h). Notably, prenatal exposure to topiramate has been linked to an increased risk of neurodevelopmental disorders, including intellectual disability, autism spectrum disorder, and attention deficit hyperactivity disorder [78–80]. Together, these results provide molecular and spatial evidence that topiramate may contribute to neurodevelopmental toxicity by disrupting Hoxb6-mediated regulatory networks during embryogenesis.

Similarly, analysis of the proteasome inhibitor bortezomib [81] showed strong spatial co-enrichment of its targets with Znf32-regulated genes from STARNet-inferred GRNs with the muscle and bone regions of the mouse embryo (Fig. 6i-j, and l). In particular, three overlapping target genes, including Psmc3, Psmd12, and Psmd7, are highly enriched in bone and muscle. PPI network analysis revealed extensive interactions and significant network proximity between bortezomib targets and Znf32-regulated genes (Z = -4.9, *P <* 0.01, Fig. 6k). GO enrichment analysis revealed that both gene sets were commonly enriched in proteasomal function, protein ubiquitination, and muscle and bone development (Fig. 6m-n). Clinically, bortezomib, an approved drug for multiple myeloma, has been reported to cause myasthenia, myalgia, and fatigue [82]. These findings provide a molecular explanation for bortezomib’s tissue-specific toxicity, potentially arising from disruption of Znf32-mediated regulatory networks.

Overall, our results underscore the effectiveness of integrating STARNet-inferred spatially specific GRNs with drug response mapping to uncover mechanistic links between drug action and region-specific gene regulation. By mapping drug targets onto spatially defined GRNs and assessing their network proximity, our integrative approach provides mechanistic insights into the origins of tissue domain-selective drug responses and adverse effects, with the potential to guide more targeted and safer therapeutic strategies.

## 3 Discussion

To the best of our knowledge, STARNet is the *first* computational framework specifically designed for inferring spatially resolved GRNs from spatial multi-omics data. Unlike existing methods tailored to either single-cell or spatial transcriptomic data alone, STARNet integrates transcriptomic and epigenomic modalities within their native spatial context. This integrative approach enables accurate inference of regulatory programs across tissue regions, thereby offering new opportunities for decoding spatial gene regulation in complex biological systems. STARNet is specifically designed to address the major computational challenges in inferring spatially resolved GRNs from spatial multi-omics data. A primary challenge lies in the systematic integration of transcriptomic and chromatin accessibility profiles within their native spatial context. STARNet overcomes this by encoding the multi-modality data into a heterogeneous hypergraph, treating spatial spots, genes, and peaks as distinct node types. Rather than relying on naive concatenation, it employs contrastive learning and a spatially informed loss function to produce unified node embeddings that preserve both modality-specific features and spatial relationships among spots. This design represents a critical advance over conventional GRN inference methods, which typically operate on single-modality data or lack spatial awareness. A second challenge stems from the high sparsity and noise inherent in spatial multi-omics datasets. To mitigate this, STARNet leverages its learned spot embeddings to define metacells, i.e., aggregates of transcriptionally and epigenetically similar spots, which serve as stable units for GRN inference, thereby improving signal robustness. Finally, quantifying the activity of GRNs within spatial domains has remained largely unexplored. To address this, STARNet incorporates a two-stage statistical framework: it first computes spot-level significance values via permutation testing and then aggregates them within spatial domains using the Cauchy combination test, yielding rigorous domain-level significance scores. Together, these innovations establish a powerful framework for accurate and spatially resolved GRN inference, overcoming key limitations of previous methods.

Building on this inference capability, STARNet supports multiple downstream applications that deepen insights into spatial gene regulation and its functional implications in complex tissues. First, it enables systematic dissection of region-specific regulatory modules, revealing how coordinated GRN activity shapes spatially localized cellular functions and cell-fate programs. Second, it integrates spatial trajectory inference to characterize spatiotemporal cell-fate specification within defined tissue regions. Third, by coupling with GWAS, STARNet pinpoints trait-associated genetic variants that may perturb spatial regulatory networks, uncovering molecular pathways by which genetic variations influence phenotypes. Finally, STARNet dissects region-specific drug response mechanisms by mapping drug targets onto spatially defined GRNs, highlighting pathways that underlie tissue-selective efficacy or toxicity. Together, these applications demonstrate STARNet’s utility in uncovering spatially resolved regulatory mechanisms across developmental, pathological, and pharmacological contexts.

Despite its broad applicability, STARNet is constrained by limitations intrinsic to current spatial multi-omics technologies. At the typical resolution of 20 µm, each spot often captures transcriptomic and epigenomic signals from multiple cells, particularly in densely packed tissues, thereby introducing intra-spot heterogeneity. Such cellular mixing can bias GRN inference in favor of the dominant regulatory programs of the most prevalent cell types. Moreover, STARNet is currently unable to model transcriptional repression, largely because it is challenging to reliably distinguish true repressive regulation from technical dropouts in sparsely expressed genes. Finally, currently STARNet only operates on two-dimensional tissue sections. Future work will extend the framework to align adjacent slices, enabling reconstruction of three-dimensional regulatory landscapes. Addressing these limitations will be critical for improving the resolution, interpretability, and dimensionality of spatial GRN analyses.

In conclusion, STARNet establishes a powerful and versatile framework for spatial multiomics analyses, offering mechanistic access to spatially resolved regulatory logic and paving the way for deeper understanding of tissue organization, developmental programs, and disease perturbations.

## 4 Methods

### 4.1 Data

#### 4.1.1 Spatial RNA-ATAC-seq dataset of P21 mouse brain

In this study, we first used a public spatial ATAC-RNA-seq dataset from a P21 mouse brain collected by Zhang et al. [14], which combines *in situ* Tn5 transposition with microfluidic barcoding to jointly profile chromatin accessibility and gene expression, yielding data from 2,373 spots covering 19,859 genes. We initiated our spatial ATAC-seq analysis using the raw fragment file provided by the authors.

To preprocess the transcriptomic data, genes expressed in fewer than 15 spots were filtered out. Subsequently, gene expression counts were normalized by library size and log-transformed using the Scanpy (v1.10.1) package [17]. To preprocess the epigenomic data, the fragment file was processed using SnapATAC2 (v2.8.0) [83] for peak calling, yielding 135,463 identified peaks. Finally, the top 40,000 highly variable peaks (HVPs), all quality-controlled genes, and all spots were selected for the construction of a heterogeneous hypergraph.

#### 4.1.2 Spatial RNA-ATAC-seq dataset of P22 mouse brain

We also used a spatial ATAC-RNA-seq dataset of a brain tissue section from a juvenile (P22) mouse, collected by Zhang et al. [14]. The dataset comprised 9,215 spatial spots and 22,914 genes. For the transcriptomic modality, genes expressed in fewer than 15 spots were filtered out. The resulting gene expression matrix was normalized by library size and log-transformed using the Scanpy package [17]. For the epigenomic modality, peak calling was performed using SnapATAC2 [83], resulting in identified 278,227 peaks. Subsequently, the top 40,000 variable peaks, along with all quality-controlled genes and all spatial spots, were selected to construct a heterogeneous hypergraph for integrative modeling of spatial multi-omics data.

To infer the spatial distribution of cell types in this dataset, a reference scRNA-seq dataset was mapped onto the spatial transcriptomic data. This reference dataset consisted of 12,692 brain-derived cells and 20,116 genes from the Tabula Muris Consortium [84]. Tangram (v1.0.4) [85] was applied with default parameters and the original cell type annotations from the reference dataset to compute the spatial probability distribution of each cell type across the spatial transcriptomic landscape.

#### 4.1.3 Spatial RNA-ATAC-seq dataset of human hippocampus

The spatial ATAC-RNA-seq dataset from a human hippocampal tissue section, collected by Zhang et al. [14], was also used in this study. The dataset consisted of 2,500 spatial spots and 29,293 genes. For the transcriptomic modality, genes expressed in fewer than 15 spots were filtered out. The resulting gene expression matrix was normalized by library size and log-transformed using the Scanpy package [17]. For the epigenomic modality, peak calling was performed using the SnapATAC2 package [83], resulting in identified 169,121 peaks. Subsequently, the top 40,000 variable peaks, all quality-controlled genes, and all spatial spots were retained to construct a heterogeneous hypergraph for integrative modeling of spatial multi-omics data.

#### 4.1.4 Spatial-Mux-seq dataset from an E13 mouse embryo

Guo et al. [15] developed a spatial multi-omics technique, termed spatial-Mux-seq, to profile an E13 mouse embryo section. Spatial-Mux-seq is a multimodal spatial profiling platform that enables simultaneous measurement of transcriptomes, chromatin accessibility, histone modifications, and targeted proteins. In this study, only the RNA and ATAC modalities were used for our downstream modeling and analyses. The dataset comprised 2,133 spatial spots. For the transcriptomic modality, genes expressed in fewer than 15 spots were filtered out. The resulting gene expression matrix was normalized by library size and log-transformed using the Scanpy package [17]. For the ATAC modality, peak calling was performed using the SnapATAC2 package [83], identifying 360,035 peaks. Subsequently, the top 40,000 variable peaks (from the epigenomic modality), all quality-controlled genes (from the transcriptomic modality), and all spatial spots were selected to construct a heterogeneous hypergraph for integrative modeling.

To annotate the spatial domains of the spatial-Mux-seq data from an E13 mouse embryo, we mapped a reference scRNA-seq dataset from E13.5 mouse embryos (Cao et al. [86]) onto the spatial multi-omics data, thus enabling spatial domain annotation based on cell-type composition. The scRNA-seq dataset initially contained 241,800 cells and 26,183 genes. To manage computational memory constraints, this dataset was reduced to a random subset of 100,000 cells. For the scRNA-seq data, we first performed quality control by excluding cells with fewer than 250 expressed genes, 500 UMIs, or over 20% mitochondrial content. The filtered data were then log-transformed, and then normalized by library size, and the top 800 highly variable genes (HVGs) were selected. Following this, we scaled the data, performed principal component analysis (PCA), and clustered the cells. Cell types were subsequently annotated based on known marker genes using Scanpy [17]. Concurrently, for the spatial transcriptomic data, the top 3,000 HVGs were identified. Finally, we used Tangram [85] with default parameters to project the annotated scRNA-seq cell types onto the spatial transcriptomic data, generating a spatial probability map of cell types.

### 4.2 Representation learning of spatial multi-omics data in STARNet

The STARNet framework is designed to comprehensively model a spatial multi-omics dataset by encoding it into a unified heterogeneous hypergraph that captures both modality-specific signals and spatial relationships. This graph-based representation effectively models the complex interplay between transcriptomic and epigenomic profiles and their spatial localizations within tissues. Through end-to-end optimization of a multi-component objective function, STARNet learns comprehensive embeddings for each node that integrates its biological state with spatial context.

#### 4.2.1 Construction of the STARNet heterogeneous hypergraph

Our graph construction is based on three primary data inputs derived from a preprocessed spatial RNA-ATAC-seq dataset. The first is a spot-by-gene matrix **X**^*SG*^ of size *N*_*s*_ × *N*_*g*_, which quantifies the expression levels of *N*_*g*_ genes across *N*_*s*_ spatial spots. The second is a binary spot-by-peak accessibility matrix **X**^*SP*^ of size *N*_*s*_ ×*N*_*p*_, indicating the presence or absence of chromatin accessibility for individual spots. The third input is the set of spatial coordinates associated with the *N*_*s*_ spots, defining their physical layout within the tissue.

The framework utilizes these inputs to construct a heterogeneous hypergraph, formally

defined as 𝒢 = (**V**, ℰ). A key innovation lies in the use of hyperedges to model the intrinsic one-to-many relationships between a single spatial spot and its associated transcriptomic and epigenomic features. By directly encoding these multi-modal associations as hyperedges, rather than relying on simple pairwise connections, the resulting graph structure more faithfully captures the complexity of spatial multi-omics data and aligns more closely with the underlying biological architecture.

##### Node sets

The node set **V** of the heterogeneous hypergraph comprises three distinct types reflecting the fundamental entities of spatial multi-omics data. These include: **(1) spot nodes** forming the set **V**_*S*_ = {*s*_*i*_|*i* = 1, …, *N*_*s*_}, representing individual spatial locations; (2) **gene nodes** forming the set **V**_*G*_ = {*g*_*j*_|*j* = 1, …, *N*_*g*_}, representing profiled genes; and (3) **peak nodes** forming the set **V**_*P*_ = {*p*_*l*_|*l* = 1, …, *N*_*p*_}, representing profiled chromatin accessibility peaks. The total node set is the union of these disjoint sets: **V** = **V**_*S*_ ∪ **V**_*G*_ ∪ **V**_*P*_. **Edge sets**. The edge set ℰ consists of two subsets that model different relationship types: **(1) spatial adjacency edges** and **(2) modality-specific hyperedges**. The spatial edges ℰ_*SS*_ *⊆* **V**_*S*_ × **V**_*S*_ are constructed by applying a k-NN algorithm to the spatial coordinates of the spots. To capture spatial dependencies across multiple tissue scales, graphs with varying values of *k* (default: 3, 4, 8) are constructed.

The second subset comprises modality-specific hyperedges, which capture one-to-many links between each spot and its associated transcriptomic or epigenomic features. The gene expression hyperedge set, denoted by ℰ_*SG*_, is constructed by connecting each spot node *s*_*i*_ to its expressed genes. More specifically, for each spot *s*_*i*_, a hyperedge 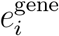 connects this spot node to all its expressed gene nodes. Similarly, the peak accessibility hyperedge set ℰ_*SP*_ links each spot *s*_*i*_ to the subset of peak nodes that are accessible at that spot.

#### 4.2.2 Multi-stage graph representation learning of STARNet

STARNet learns rich node representations through a multi-step representation learning process, beginning with the initialization of learnable embeddings for spots, genes, and peaks. Then modality-specific spot representations are computed by aggregating node features within each modality-specific hyperedge using a hybrid aggregation strategy that captures both comprehensive information and the most salient features. To integrate these modality-specific embeddings, a two-tier attention mechanism is employed: a modality-level attention module first estimates the relative importance of the gene and peak modalities for each spot, and then a subsequent cross-attention module fuses this information to refine the previous spot embeddings. This yields a unified, modality-aware representation for each spot. Finally, these embeddings are further refined through multi-scale Graph Convolutional Networks (GCNs) applied over the spatial adjacency graph, allowing local information to propagate across neighboring spots. The resulting node representations encode both multi-omic biological features and their spatial context, supporting downstream spatial GRN inference and analyses.

##### Node embedding initialization

Each node in the heterogeneous hypergraph is initialized with a learnable feature vector of dimensionality *d* (default *d* = 128). These initial embeddings are organized into three distinct matrices corresponding to the three node types: 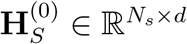 for spatial spots, 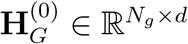 for genes, and 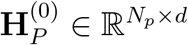 for peaks. All embeddings are initialized using values drawn from a standard normal distribution, and their parameters are jointly optimized in an end-to-end manner during model training.

##### Learning modality-specific spot representations

To learn modality-specific representations for each spot, the framework aggregates features from the corresponding modality hyperedges. This aggregation aims to extract salient features that characterize the biological signature of each spot in a context-specific manner. The process begins with the raw spot-modality matrix **X**^*Sm*^, where *m ∈* {gene, peak}. To mitigate biases from features that are globally prevalent across the tissue, the matrix is column-wise normalized to produce 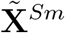, that is:

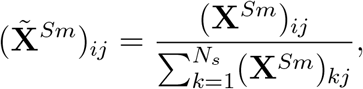

where the subscript *ij* denotes the association between spot *i* and feature *j*, while the denominator normalizes feature *j*’s abundance across all spots. This normalization downweights features that are ubiquitously expressed across the tissue and emphasizes those that are preferentially associated with a specific spot’s hyperedges. Through this normalization strategy, the learned representation for each spot can more accurately capture its modality-specific molecular features.

Building upon the initial normalization, the model further sharpens the feature signals by prioritizing the most informative components within each modality hyperedge. This is achieved by computing a salience-weighted probability matrix **P**^(*m*)^, *m ∈* {gene, peak}, which guides the subsampling of features during aggregation. The matrix is computed by applying a power transformation to the normalized matrix 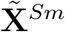, followed by row-wise normalization, that is,

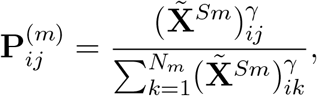

where *γ* (default: 3) is a tunable hyperparameter that controls the degree to which dominant features are emphasized. This transformation produces a spot-specific probability distribution over modality features, assigning nonlinearly higher weights to features with stronger modality-specific expression or accessibility, thereby refining the focus on biologically salient signals.

This probability matrix then guides a stochastic subsampling procedure that focuses on the most informative features in each modality-specific hyperedge. For each spot, a fixed number of salient features *k*_*m*_ are sampled according to the probability distributions in **P**^(*m*)^ (default: *k*_*gene*_ = 20, *k*_*peak*_ = 100). This sampling yields a sparse binary mask 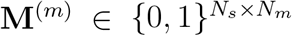, which selectively retains high-salience features while discarding low-contribution ones. This strategy reduces noise from ubiquitous or non-discriminative features and sharpens the representation of each spot’s modality-specific signals.

Building on this, STARNet constructs a hybrid aggregation matrix that integrates both global and focused perspectives. Specifically, it combines two independently row-normalization operators: one based on the full normalized modality matrix 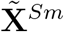, and the other derived from the masked, salience-weighted matrix 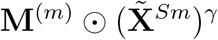. These are linearly combined to yield the final modality-specific spot representation 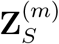, that is,

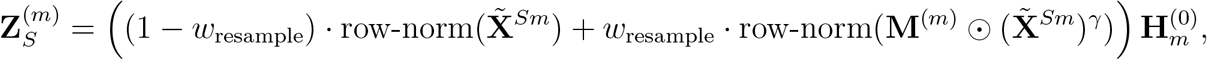

where *w*_resample_ (default: 0.8) controls the balance between the two perspectives, and ⊙ denotes elementwise multiplication. This process is performed independently for gene and peak modalities, resulting in two independent representations 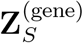 and 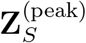. This hybrid mechanism allows the model to dynamically emphasize biologically critical features while retaining the broader feature context necessary for robustness.

##### Learning a unified spot representation

To create a unified and biologically coherent representation for each spot, STARNet fuses the modality-specific embeddings using a two-level attention mechanism.

The first level, a modality-level attention module, dynamically estimates the relative contribution of the gene and peak modalities to the overall spot representation. For each modality *m ∈* {gene, peak}, the model first computes a global context vector **c**^(*m*)^ by applying a learnable linear transformation followed by a tanh activation to the modality-specific embeddings 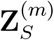, and then averaging the transformed vectors across all spots, that is,

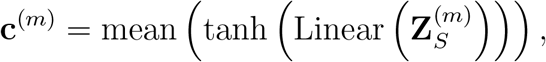

Then the importance score for each modality, denoted by *s*^(*m*)^, is obtained by measuring the similarity between its context vector and a learnable query vector **q**:

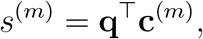

These scores are passed through a softmax function to yield normalized attention weights, that is,

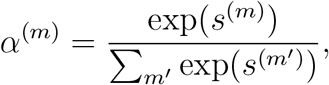

The fused representation 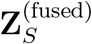 is then computed as the weighted sum of modality-specific embeddings, followed by a ReLU activation:

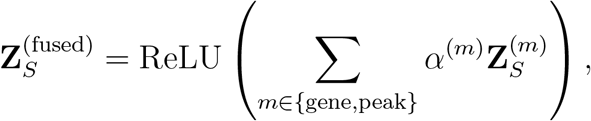

This process enables the model to adaptively prioritize the most informative modality for each spot, ensuring that the fused embedding reflects the dominant multi-modal signals in a spatially and biologically relevant manner.

The second level of the attention mechanism fuses the modality-integrated representation 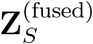 with the previous spot embeddings 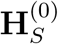 through a cross-attention mechanism. In this formulation, each spot’s fused embedding acts as the query (**Q**), while the previous spot embeddings serve as both the key (**K**) and the value (**V**). This allows each spot’s representation to be dynamically updated in proportion to its alignment with the integrated multi-modal embeddings, ensuring that the most relevant spatial and molecular context is incorporated. The updated, multi-modality-informed spot representation, denoted by 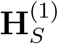, is computed using the standard scaled dot-product attention:

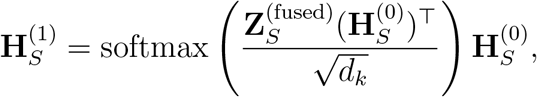

where *d*_*k*_ is the dimensionality of the key vectors. This operation effectively enriches the previous embeddings of each spot with cross-modal molecular information, yielding a biologically holistic representation.

##### Spatial contextualization via graph convolutional networks

Although the representation 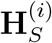 captures integrated multi-modal signals, it remains agnostic to the spatial context of each spot. To incorporate this crucial information, the framework applies multi-scale spatial propagation using graph convolutional networks (GCNs). This involves applying GCN layers across a set of neighborhood graphs 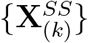, each corresponding to a different spatial resolution defined by varying the number of nearest neighbors (i.e., different values of *k* in the k-NN procedure). For each scale *k*, a spatially enriched embedding, denoted by 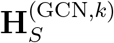, is computed using the standard GCN update rule:

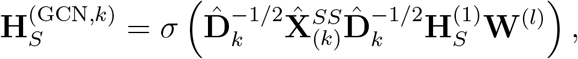

where 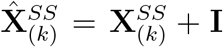 is the adjacency matrix for scale *k* with self-loops, 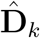 is its degree matrix, **W**^(*l*)^ is a learnable weight matrix, and *σ*(*·*) is a nonlinear activation function (default: ReLU). This multi-scale GCN scheme enables each spot embedding to be refined by integrating information across spatial scales, ranging from immediate neighbors to neighboring spots at larger spatial radius, thereby capturing features of spatial organization such as domain boundaries and expression gradients.

To synthesize the final embedding, the fused multi-modal representation 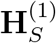 is concatenated with all spatially refined embeddings across scales, denoted by 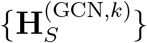. This concatenated tensor, which encodes both modality and spatial hierarchy information, is passed through a final linear transformation to produce the ultimate spot representation, denoted by 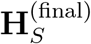:

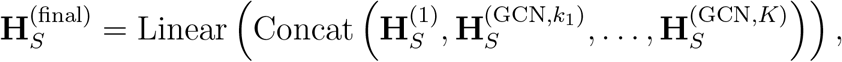

This final embedding unifies multi-modal biological information with spatial structure across scales, serving as a robust foundation for downstream spatial GRN analyses and related biological tasks.

#### 4.2.3 End-to-end training with a composite objective function

The STARNet model is trained end-to-end by optimizing a composite multi-objective loss function. This formulation simultaneously enforces several key properties of the learned embeddings: reconstruction fidelity to the input data, alignment across transcriptomic and epigenomic modalities, and preservation of spatial relationships. The overall loss function is defined as a weighted sum of these three components, ensuring that the resulting embeddings are both biologically meaningful and robust across data modalities and spatial context.

##### Modality reconstruction loss

The first component, the modality reconstruction loss (ℒ_recon_), enforces fidelity to the original input data. It ensures that the learned embeddings retain sufficient information to reconstruct the observed spot-gene and spot-peak matrices. Reconstruction is performed by computing the cosine similarity between the final spot embeddings and the corresponding gene or peak embeddings. The loss is then computed as the mean squared error (MSE) between these reconstructed ma trices and the original inputs, that is,

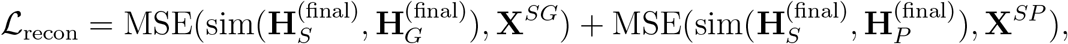

where sim(*·, ·*) denotes the cosine similarity function, 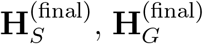, and 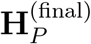 are the final embeddings for spots, genes, and peaks, respectively, and **X**^*SG*^ and **X**^*SP*^ are the original spot-gene and spot-peak matrices, respectively.

##### Cross-modal contrastive loss

The second component, the cross-modal contrastive loss (ℒ_con_), enforces the alignment of gene and peak modality representations within a shared embedding space. The goal is to ensure that the spot-level embeddings derived independently from gene expression and chromatin accessibility hyperedges are functionally consistent for the same spatial location. This is achieved by maximizing mutual information between modality-specific embeddings of the same spot, while simultaneously pushing apart embeddings from different spots. The objective is formalized using an InfoNCE loss:

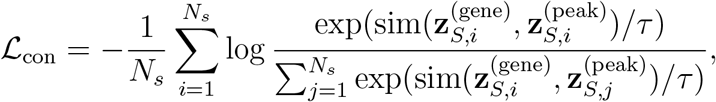

where 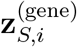 and 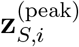 denote the gene- and peak-derived embeddings for the *i*-th spot (i.e., the *i*-th rows of 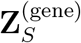 and 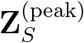, respectively), and *τ* is a temperature scaling factor (default: 0.02) that controls the sharpness of the similarity distribution.

This formulation drives the model to learn modality-consistent spot representations, which are critical for robust integration of multi-omics data.

##### Spot link loss

The third objective, the spot link prediction loss (ℒ_link_), enforces the learned spot embeddings to preserve meaningful neighborhood relationships. It operates on three graph structures: a spatial graph (**A**_spatial_) derived from spot coordinates, and two feature-similarity graphs (**A**_gene-sim_ and **A**_peak-sim_) constructed based on within-modality similarities.

To model these relationships, STARNet adopts a contrastive learning framework that distinguishes between true spot pairs and randomly sampled negatives. For each positive edge (*i, j*) defined in a given graph, the model minimizes an InfoNCE loss:

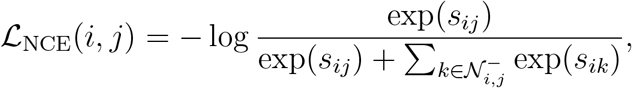

where *s*_*ij*_ denotes the similarity score between spot embeddings *i* and *j*, and 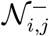 is a set of randomly sampled negative pairs (default: 32 negatives per positive).

To improve robustness to potential noise in the graph structure, this contrastive loss is combined with a bootstrapping regularizer, which encourages consistency between the current prediction and a “soft” target derived from the model’s own detached prediction:

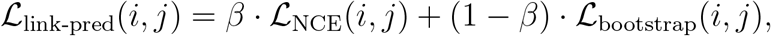

where ℒ_bootstrap_ is the cross-entropy between the detached soft targets and current predictions, and *β* is the balancing factor, which is set by default to 0.8 for the spatial graph and 0.9 for the feature-similarity graphs, reflecting their assumed noise levels. Then the total loss for a given graph, denoted by ℒ_link-pred_(**S**_sim_, **A**), is then calculated as the mean of this per-pair loss over all positive edges (E_**A**_) in that graph:

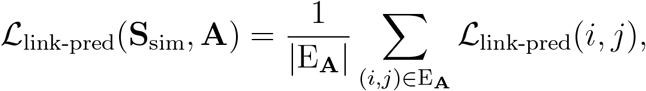

The construction of the graphs is as follows: **Spatial graph (A**_**spatial**_): constructed by connecting each spot to its *k* nearest spatial neighbors (default: *k* = 4) based on Euclidean distance. **Feature-similarity graphs (A**_**gene-sim**_, **A**_**peak-sim**_): constructed by first applying PCA (default: 32 components) to the spot-feature matrices, and then connecting each spot to its *k* nearest neighbors (default: *k* = 8) in the dimensionally reduced space.

The final link prediction loss is a weighted combination of the loss values across the three graphs, that is,

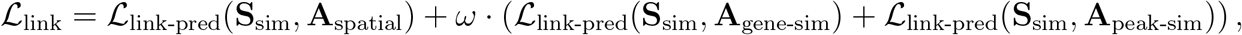

where 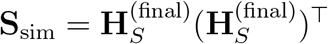 denotes the pairwise similarity between final spot embeddings, and *ω* is the weighting parameter (default: *ω* = 0.6).

##### Final loss

The model is trained end-to-end by optimizing a composite objective function, denoted by ℒ_total_, which linearly combines the above three loss components, including the reconstruction loss, the cross-modal contrastive loss, and the spot link prediction loss. This multi-task learning objective enforces the model to jointly preserve input fidelity, cross-modal alignment, and spatial/functional neighborhood structure. Thus, the total loss is defined as:

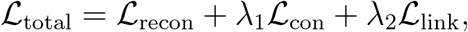

where *λ*_1_ and *λ*_2_ stand for scalar weighting hyperparameters that control the relative contribution of individual loss terms. By default, these are set to *λ*_1_ = 0.5 and *λ*_2_ = 0.025, allowing balanced optimization while slightly down-weighting the structural regularization component. These weights can be tuned to emphasize different aspects of representation learning depending on the application contexts.

#### 4.2.4 Implementation Details

By default, STARNet was trained end-to-end for 600 epochs using the AdamW [87] optimizer. The initial learning rate was set to 5 × 10^*−*3^, with a weight decay of 1 × 10^*−*4^ for regularization. The model was implemented using PyTorch (v2.3.1) and PyTorch Geometric (v2.5.3). All computational experiments were conducted on a workstation equipped with an NVIDIA GeForce RTX 4090 D GPU (24 GB VRAM).

### 4.3 Construction of spatially specific GRNs

#### 4.3.1 Identification of an initial set of spatially specific TFs

STARNet first uses the CIS-BP database [88] to obtain TF binding motifs. A fundamental assumption of STARNet is that TFs essential to a spatially specific GRN should themselves exhibit a non-random spatial expression pattern. To generate an initial set of candidate TFs, we assessed the spatial autocorrelation of all TF expression profiles using Moran’s I statistic, as implemented in the Squidpy (v1.5.0) package [89]. Based on this analysis, TFs with a Moran’s I score below 0.01 were filtered out, and only those exhibiting significant spatial patterns were retained for subsequent GRN construction.

#### 4.3.2 Construction of the initial GRN set

The STARNet model generates gene embeddings that encode spatial co-expression patterns, with higher embedding similarity directly corresponding to stronger spatial co-expression between genes. We leveraged this property to identify an initial set of potential regulatory interactions between TFs and their target genes. More specifically, for each TF, we selected the top 500 TF-target pairs exhibiting the highest cosine similarity scores to construct an initial set of GRN candidates.

To refine this initial GRN set and reduce spurious connections driven by stochastic gene expression, we applied a secondary filtering step using Pearson correlation calculated from aggregated expression profiles. More specifically, to counteract the inherent sparsity of spatial multi-omics data, we first generated metacells using the same pipeline as in SEACells [90], with our STARNet-generated spot embeddings as input. The gene expression counts from all spots within each metacell were then aggregated. Using these aggregated metacell expression profiles, we calculated the Pearson correlation coefficient for each TF-target pair in the initial GRN set. To ensure robust co-expression links, pairs with a correlation coefficient of 0.2 or lower were excluded from the final network.

#### 4.3.3 Inference of candidate CREs and Peak2GeneLinks

We hypothesized that TFs bind to candidate CREs to activate target gene expression in spatially specific GRNs. To identify significant Peak2GeneLinks, we followed the same procedure as in SEACells [90]. Specifically, Spearman correlation was computed between the normalized accessibility of each peak and the normalized expression of each gene. The search window included the gene body and extended 100 kb both upstream and downstream of its transcription start site. These calculations used normalized accessibility profiles from spatial ATAC-seq metacells and normalized expression levels from spatial RNA-seq metacells, which were derived from the spot embeddings generated by STARNet. To assess the significance of each peak-gene correlation, an empirical background was established by sampling 100 control peaks matched for GC content and accessibility. Given the inherent sparsity and noise of spatial multi-omics data, we adopted a relatively relaxed threshold (empirical *P <* 0.2) to avoid excessive false negatives while retaining potentially relevant regulatory links. Peak2GeneLinks meeting this criterion were considered significant, and their associated peaks were designated as candidate CREs for downstream analyses.

#### 4.3.4 Construction of final spatially specific GRNs

The construction of the final spatially specific GRNs was achieved by integrating evidence of co-expression with direct evidence of physical TF binding. To achieve this, we performed motif scanning within the set of previously identified candidate CREs. Potential binding sites for each TF were identified using the function “motif.with nucl prob().exists()” from the SnapATAC2 package [83]. A stringent statistical threshold (*P <* 10^*−*4^) was used to define a significant motif match. In STARNet, a TF-target gene pair was retained in the final GRNs only if its associated CREs harbored a statistically significant binding motif matching the TF.

Each retained regulated link is thus supported by independent lines of evidence: a significant TF-binding motif within a linked CRE and the spatial co-expression of the TF and target.

#### 4.3.5 Estimation of GRN significance at single-spot resolution

To quantify the activity of each GRN at single-spot resolution, we adapted the same procedure as in scDRS [91] to assess whether a spot exhibits significant co-expression among genes within the GRN’s target set. This approach yields spot-level *P*-values for GRN-associations by comparing each spot’s GRN score to a null distribution generated from control gene sets. More formally, given a GRN, for each spot *i* = 1, …, *n*_*spot*_, we first define its row GRN score, denoted by *R*_*i*_, and a control score, denoted by 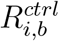:

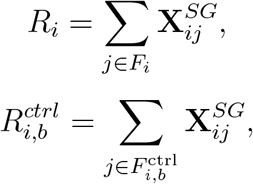

where **X**^*SG*^ stands for the input spatial transcriptomic data, *F*_*i*_ stands for the set of target genes regulated by this GRN at spot *i*, and 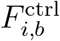 stands for the *b*-th control gene set, with a default of 1000 control sets. Each control set was matched to the GRN’s target gene set *F* in terms of both size and expression properties (mean and variance) of the corresponding genes.

To enable comparison across all spots, these raw scores *R*_*i*_ and 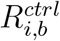 were normalized using Z-score normalization strategy. Then the final *P*-value was computed by comparing each spot’s normalized GRN score (*s*_*i*_) against a pooled empirical null distribution of all normalized control scores from all spots 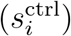, that is,

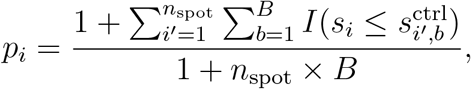

where *p*_*i*_ stands for the GRN’s *P*-value for spot *i, s*_*i*_ stands for the normalized GRN score for spot *i, B* represents the number of control gene sets (the number of control sets *B* is set to 1,000 by default), 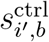 represents the normalized control score for spot *i*^*′*^ from the *b*-th control gene set, 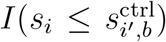 stands for an indicator function that equals to 1 if *s*_*i*_ is less than or equal to *s*_ctrl,*i′,b*_, and 0 otherwise, and *n*_*spot*_ represents the number of spots.

The above test evaluates whether the aggregate expression of the GRN’s target genes in a given spot is significantly higher than that of control gene sets with matched size and expression properties, thereby quantifying the GRN activity in that specific spot and enabling downstream statistical analyses.

#### 4.3.6 Estimation of GRN significance at domain resolution

To assess the overall enrichment of a GRN’s activity within a specific spatial domain, STARNet employs the same Cauchy combination test as in gsMap [53]. This test aggregates the *P*-values of individual GRNs from all spots within a given domain into a single domain-level significance score.

More formally, a combined test statistic, denoted as *T*_Cauchy_, for the domain is calculated as follows:

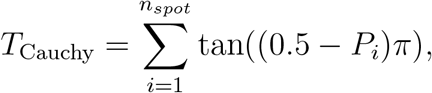

where *n*_*spot*_ stands for the number of spots within the spatial domain, *P*_*i*_ represents the *P*-value of the GRN at spot *i*, and tan(*·*) stands for the tangent function. Then the aggregated *P*-value for the spatial domain is approximated as:

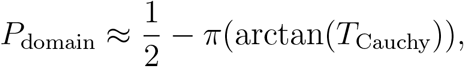

where arctan(*·*) stands for the arc tangent function. *P*_domain_ quantifies the overall significance of a GRN’s activity in that domain, serving as a statistical test for domain-specific GRN enrichment.

#### 4.3.7 Computation of regulatory scores for TF–target gene pairs

The regulatory score quantifies the contribution of a target gene to the activity of its regulating TF within the GRN. For each TF–target pair, we assess the change in the GRN significance score after computationally removing the target gene from the network. A larger decrease in significance indicates a stronger regulatory influence of that target gene. This score can be used to prioritize and visualize the most influential TF–target interactions in the network.

To calculate this score, we first define the following two aggregated *P*-values using the Cauchy combination test.

*P*_All Targets_: The aggregated *P*-value for the GRN, obtained using the Cauchy combination

test across all spots, where the GRN’s activity is statistically significant (*P <* 0.05). For each spot, *P*-values are computed from all target genes in the GRN using a permutation test. To accelerate computation, the number of control sets was fixed at 200.

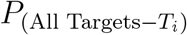: The aggregated *P*-value for the GRN when using all targets except gene *i* (*T*_*i*_) are used, computed in the same as *P*_All Targets_ for each spot.

The raw regulatory score *Y*_*i*_ for target gene *i* is defined as the difference of the negative logarithms between these two aggregated *P*-values, that is,0

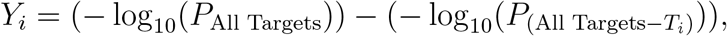

To enable comparability among target genes within a GRN, these raw regulatory scores are min-max normalized to the range [0,1], that is,

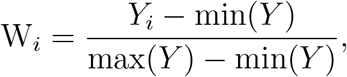

where min(*Y*) and max(*Y*) represent the minimum and maximum raw regulatory scores across all target genes in the GRN, respectively. This normalization facilitates consistent interpretation of regulatory scores within individual GRNs.

### 4.4 Integrative analyses and applications of the STARNet model

#### 4.4.1. Clustering of GRNs by spatial activity patterns

To identify functionally related GRNs that are co-regulated within specific spatial contexts, we developed an unsupervised clustering framework that groups the STARNet-inferred GRNs based on their spatial significance patterns. This approach systematically reveals GRN modules, i.e., groups of GRNs that may cooperatively maintain domain-specific functions.

Our clustering pipeline begins with a matrix where rows represent STARNet-inferred GRNs and columns represent spots, with each entry being the *−*log_10_*P* of GRN activity from our permutation test. We first applied PCA to this matrix for dimensionality reduction. Based on the resulting principal components, we then constructed a k-NN graph to capture the similarity structure among GRNs. Finally, the Leiden community detection algorithm was applied to this graph to identify distinct GRN modules [92].

To determine the significance of a GRN module at each spot, we performed a permutation test on a combined gene set comprising all TFs and target genes within that module. Then the associations between spatial domains and GRN modules were calculated using the Cauchy combination test [16].

#### 4.4.2 Spatiotemporal analysis

Spatial trajectories were inferred from the spatial multi-omics data using spaTrack (v1.0.2) [42] with default parameters. SpaTrack is a method for spatial trajectory inference that estimates cell transition costs by integrating both gene expression similarity and spatial proximity.

The spatial domain with the lowest Shannon entropy [93] was selected as the root. For each domain, Shannon entropy was calculated from the aggregated gene expression profiles as,

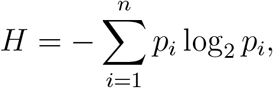

where *p*_*i*_ represents the probability of expression for the *i*-th gene, and *n* stands for the number of genes.

#### 4.4.3 GWAS analysis

To link human complex traits to the spatially specific GRNs inferred by STARNet, we implemented a multi-step analytical pipeline. First, we identified trait-associated spatial domains by integrating GWAS summary statistics with the transcriptomic modality of spatial multiomics data using gsMap (v1.72.3) [53]. Within these domains, key STARNet-inferred GRNs were selected by applying the Cauchy combination test to GRN activity *P*-values across all constituent spots.

Finally, significant GWAS SNPs were mapped to candidate target genes when overlapping a TF-binding peak linked to that gene within an identified key GRN. Each gene was then assigned the most significant *P*-value from its associated SNPs.

3D structural models of the NEUROD2–DNA complex were generated using AlphaFold3 (v3.0) [71] with default parameters via the AlphaFold3 web server (https://alphafoldserver.com/). The DNA-binding domain sequence of NEUROD2 was paired with either the reference (CCCATATGCC) or alternate (CCCATATACC) DNA motif sequence. For each motif, five structural models were produced, and the top-ranked model (by AlphaFold3 confidence score) was selected for downstream analysis.

Protein–DNA binding interface properties were calculated using PDBePISA [72] (https://www.ebi.ac.uk/pdbe/pisa/), including interface area, number of hydrogen bonds, and solvation free energy gain (Δ*iG*), to compare binding affinities between motifs. Larger interface areas, more hydrogen bonds, and more negative Δ*iG* values indicate stronger binding. Structural visualizations (Fig. 5i) were prepared using PyMOL [94].

#### 4.4.4. Analysis of spatially resolved drug responses

To assess potential tissue domain-specific drug responses, we first mapped the expression of drug target genes onto the spatial transcriptomic atlas using drug2cell [95]. Drug targets were curated from ChEMBL (v.30) [96], and the mean expression of each drug’s target genes was calculated for each spot, serving as a proxy for the potential response to the drug. To determine whether a drug preferentially affects a specific spatial domain, we performed Gene Set Enrichment Analysis (GSEA) for the drug target genes within each domain [97]. Significance was assessed using a permutation-based GSEA, with thresholds of normalized enrichment defined by a normalized enrichment score (NES) *>* 2, FDR *<* 0.25, and *P <* 0.05. Drug-GRN pairs were prioritized if both drug targets and GRNs were significantly co-enriched in the same spatial region and shared at least three target genes.

We further evaluated the network proximity between drug targets and GRN targets using a human PPI network compiled from 15 databases of experimentally validated interactions [98], comprising 243,603 interactions among 16,677 proteins. The largest connected component was extracted, and for each drug target, the average shortest-path length to its nearest GRN target was computed. Statistical significance was determined by generating 1,000 degree-matched random sets to form an empirical null distribution.

### 4.5 Benchmarking of spatial domain identification

#### 4.5.1 Simulated data generation for spatial domain identification benchmarking

To rigorously evaluate the model’s performance in the spatial domain identification task, we generated five distinct simulated spatial multi-omics datasets. Specifically, this was achieved by sampling cells from published single-cell multi-omics datasets into predefined spatial domains, thereby creating a ground-truth for benchmarking.

The overall simulation strategy was adapted from the SpatialGlue pipeline [24]. For each dataset, we defined four distinct spatial domains and populated them with a total of 1,296 spots. To ensure the robustness of our evaluation, each of the five simulated datasets was generated using different combinations of source data, cell types, and cell-type compositions. The count matrices of all simulated datasets were normalized by library size and then log-transformed, and the specific composition of each simulated dataset is as follows:

##### Simulated Dataset 1

This dataset was generated using single-cell SNARE-seq data from the adult mouse cerebral cortex [99]. We simulated four spatial domains by sampling 648 E2Rasgrf2 cells, 396 E3Rorb cells, 36 E5Galnt14 cells, and 216 E4Il1rapl2 cells into domains 1, 2, 3, and 4, respectively.

##### Simulated Dataset 2

For this dataset, we utilized a multiome dataset of CD34+ hematopoietic stem and progenitor cells from human bone marrow [90]. The spatial domains were populated with 648 hematopoietic stem cells (HSCs), 396 hematopoietic multipotent progenitors (HMPs), 36 common lymphoid progenitors (CLPs), and 216 monocytes in domains 1, 2, 3, and 4, respectively.

##### Simulated Dataset 3

We again used the adult mouse cerebral cortex SNARE-seq data [99]. The four domains were composed of 612 E6Tle4 cells, 288 E5Parm1 cells, 216 E5Sulf1 cells, and 180 OliM cells in domains 1, 2, 3, and 4, respectively.

##### Simulated Dataset 4

This dataset was also generated from the human bone marrow multiome data [90]. The domains were populated with 612 HSCs, 288 HMPs, 216 CLPs, and 180 monocytes.

##### Simulated Dataset 5

The final simulated dataset was also derived from the human bone marrow multiome data [90]. The four spatial domains comprised 612 HSCs, 288 HMPs, 216 megakaryocyte-erythroid progenitors (MEPs), and 180 erythrocytes.

By creating these varied and well-defined simulated environments, we comprehensively assessed and benchmarked the performance of different methods for spatial domain identification.

#### 4.5.2 Benchmarking methods for spatial domain identification

To evaluate the performance of STARNet in spatial domain identification, we compared it with eight competing methods, including single-cell omics integration tools (Scanpy [17], MultiVI [18], scGLUE [19]); spatial transcriptomic domain identification methods (GraphST [20], STAGATE [21]); a spatial ATAC-seq processing method Descart [22]; and spatial multiomics integration methods (COSMOS [23], SpatialGlue [24]).

All benchmarking methods were executed according to their provided pipelines. After generating spot embeddings for each model, we constructed a spot-spot k-NN graph (k = 15), followed by clustering using the Leiden algorithm [92]. We manually tuned the parameters to obtain four clusters, which were then used for subsequent benchmark calculations.

Below, we provide details of each method evaluated:

Scanpy [17] is a scalable Python toolkit for scRNA-seq analysis. We first selected the top 3,000 HVGs from the RNA modality. Spot embeddings were subsequently generated using principal component analysis via the sc.pp.pca function with default parameters.

MultiVI [18] is a deep generative model based on a variational autoencoder (VAE) architecture for integrating single-cell multi-omics data. To prepare the input for the MultiVI model, we selected the top 3,000 HVGs and 50,000 HVPs.

scGLUE [19] is a computational framework that integrates single-cell multi-omics data by using a regulatory interaction graph as prior knowledge. Following its standard pipeline, we first performed dimensionality reduction on each modality independently. PCA was applied to the RNA data, while Latent Semantic Indexing (LSI) was applied to the ATAC data. These reduced-dimension representations were then used as input for the scGLUE model.

GraphST [20] is a deep learning method that combines graph neural networks (GNNs) with self-supervised contrastive learning to analyze spatial transcriptomic data. The top 3,000 HVGs were selected as input for the GraphST model.

STAGATE [21] is a graph attention autoencoder framework that learns low-dimensional embeddings by integrating gene expression with spatial information. The model first constructs a spatial neighbor network from spot coordinates. We used the top 3,000 HVGs as input for the STAGATE model.

Descart [22] is an iterative model designed to denoise, enhance, and identify spatially variable peaks in spatial ATAC-seq data. In line with its protocol, we generated spot embeddings by performing PCA on the enhanced feature matrix derived from the spatial ATAC-seq data. COSMOS [23] is a GNN-based algorithm for multi-omics integration. It encodes each data modality using separate GCNs and generates a unified spot representation by using a weighted nearest neighbor algorithm to balance the modalities. The top 50,000 HVPs were used as the input for the ATAC modality.

SpatialGlue [24] is a GNN model featuring a dual-attention mechanism for spatial multiomics data integration. For preprocessing, we first filtered out genes expressed in fewer than 10 spots. Subsequently, PCA was applied to the RNA modality, and LSI was applied to the ATAC-seq modality. These modality-specific representations were then provided as input to the SpatialGlue model, which computes the final integrated embeddings.

#### 4.5.3 Evaluation metrics

To evaluate the performance of different benchmarking approaches in spatial domain identification, we used six supervised metrics: adjusted mutual information (AMI), normalized mutual information (NMI), adjusted rand index (ARI), mutual information (MI), V-measure, and homogeneity-implemented in the scikit-learn package [100]. Higher values indicate better performance for spatial domain identification.

The **Adjusted Rand Index (ARI)** compares the similarity between predicted spot labels from a clustering algorithm and their ground-truth labels, adjusting the Rand Index (RI) to account for agreements that may occur by chance.

Let *C* be the set of ground-truth labels and *K* be the set of predicted spot labels. We define *a* as the number of spot pairs assigned to the same cluster in both *K* and *C, b* as the number of spot pairs assigned to different clusters in *K* and *C*, and *n* as the total number of spots.

Then the Rand Index (*RI*) is calculated as:

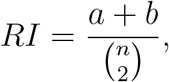

The *ARI* is then derived as:

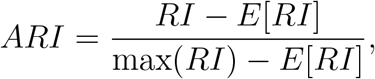

where *E*[*RI*] stands for the expected *RI* for a random clustering. *ARI* values range from -1 to 1, with 1 indicating perfect agreement and -1 perfect disagreement.

The **Mutual Information (MI), Adjusted Mutual Information (AMI)**, and **Normalized Mutual Information (NMI)** are information-theoretic measures used to quantify the consistency between predicted spot labels and truth labels.

Given two sets of spot labels, *U* = {*U*_1_, …, *U*_|*U*|_} (ground-truth) and *V* = {*V*_1_, …, *V*_|*V* |_}

(predicted), their joint probability distribution *P* (*i, j*) is the probability that a spot belongs to cluster *U*_*i*_ in *U* and *V*_*j*_ in *V*. Marginal probabilities are given by 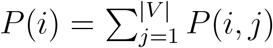 and 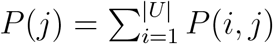.

Then the *MI* between *U* and *V* is defined as:

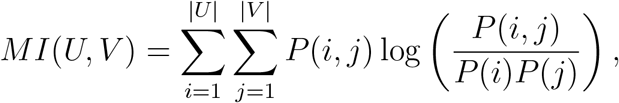

To account for chance, *AMI* is defined as:

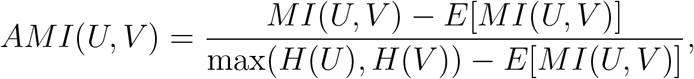

where *H*(*U*) and *H*(*V*) stand for the entropies of *U* and *V*, respectively, defined as *H*(*X*) = *−* ∑ *P* (*x*) log *P* (*x*). *AMI* values typically range from 0 (random agreement) to 1 (perfect agreement), with values near 0 indicating no correlation beyond chance.

The *NMI* normalizes *MI* to the range [0, 1]:

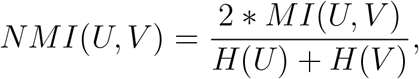

**Homogeneity** measures whether each predicted cluster contains only spots from a single ground-truth label, defined as:

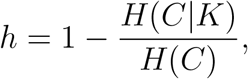

where *H*(*C*|*K*) is the conditional entropy of the ground-truth labels given predicted labels, and *H*(*C*) is the entropy of ground-truth labels. A Homogeneity score of 1 indicates that each predicted cluster contains only spots from a single ground-truth class.

The **V-measure** combines homogeneity (*h*) and completeness (*c*) as the harmonic mean:

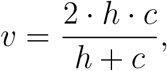

A V-measure score of 1 indicates a perfectly homogeneous and complete clustering.

#### 4.5.4 Benchmarking of spatial domain identification on a real dataset

We further evaluated STARNet’s performance in identifying spatial domains using the human hippocampus spatial RNA-ATAC-seq dataset [14]. To assess accuracy, annotations from the Allen Brain Atlas were used as anatomical references, enabling direct comparison between predicted domains and known anatomical subregions. STARNet was benchmarked against state-of-the-art methods to evaluate its ability to recover anatomically coherent spatial structures.

### 4.6 Benchmarking of spatially resolved GRN inference

To systematically evaluate state-of-the-art methods for GRN inference from spatial multiomics data, we developed a benchmarking framework incorporating both simulated and real-world datasets. Each method was run with its default parameters to infer GRNs, and the resulting networks were quantitatively compared against multiple ground-truth references.

#### 4.6.1 Generation of simulated spatial multi-omics data

To generate the simulated benchmark dataset, we first constructed a spatially coherent tissue architecture with four adjacent domains, following Section 4.5.1. Based on this layout, we simulated a total of 4,000 spatial spots, with 1,000 spots assigned to each domain. For each spot, we generated RNA-seq and ATAC-seq profiles comprising 5,000 RNA reads and 15,000 chromatin fragments, respectively.

To emulate biological tissue heterogeneity, each spot contained both dominant and secondary cell types. Each domain’s dominant cell type accounted for 76% of the spot’s cellular composition, while the remaining 24% was split evenly among the dominant cell types of adjacent domains, simulating realistic tissue mixing.

Simulated sequencing data were generated by sampling from bulk RNA-seq and ATAC-seq datasets of four human cell lines, including GM12878, HepG2, MCF-7, and K562, sourced from the ENCODE Project [25, 26]. For ATAC-seq, peaks were first identified using MACS2 [101], and fragments were sampled from these peaks according to the domain-specific cell type composition. We similarly sampled RNA-seq reads and quantified gene expression using featureCounts with GENCODE v47 gene annotation.

Finally, a composite tissue slice of 1,296 spots was assembled by random sampling from the 1,000 pre-generated spots per domain. The distribution across domains was as follows: Domain 1: 612 spots with GM12878 as the dominant cell type, and MCF-7, K562, and HepG2 as secondary types; Domain 1: 612 spots with GM12878 as the dominant cell type, and MCF-7, K562, and HepG2 as secondary types; Domain 3: 216 spots with MCF-7 as the dominant cell type, and GM12878 and K562 as secondary types; Domain 4: 180 spots with K562 as the dominant cell type, and GM12878 and MCF-7 as secondary types.

#### 4.6.2 Benchmarking methods for spatial GRN inference

**STARNet** was executed using default hyperparameters. The paired spatial RNA-seq and ATAC-seq data were preprocessed with Scanpy [17] and SnapATAC2 [83], respectively. Preprocessing steps included log-transformation, selection of the top 40,000 HVPs and removal of genes expressed in fewer than 15 spatial spots. The model was trained for 600 epochs with a learning rate of 5 × 10^*−*3^. To infer the GRN, metacells were constructed by aggregating transcriptomic and epigenomic count matrices across all spatial spots, using the spot embeddings inferred by STARNet as the basis for aggregation. Next, candidate TFs were identified based on spatial autocorrelation using Moran’s I score *≥* 0.01. A preliminary GRN was assembled by linking TFs to genes with Pearson correlation coefficients exceeding 0.2. These candidate links were subsequently refined by scanning for TF binding motifs within putative CREs, retaining only those with motif match *P <* 10^*−*4^. The scanning window encompassed the gene body as well as ±100 kilobases around the transcription start site (TSS).

**SCENIC+** [10] was run with default parameters according to its official documentation (https://scenicplus.readthedocs.io/en/latest/). Given that the dataset was simulated, the initial quality control step was omitted. Topic modeling was performed using pycisTopic with MALLET (500 iterations), exploring 11 models with topic numbers ranging from 2 to 50 in increments of 5. The model with 40 topics was selected based on performance metrics. The core SCENIC+ pipeline was then applied using both consensus peaks and SCREEN regions [102] as input, using the October 2016 Ensembl archive as the BioMart host. Regulons were considered high-confidence if they exhibited a Pearson correlation greater than 0.7 between gene-based and region-based AUC scores.

**pySCENIC** [9, 103] was executed via its command-line interface (https://pyscenic.readthedocs.io/en/latest/). The GRNBoost2 algorithm [104] was used to infer co-expression modules from the raw count matrix. These modules were then refined into regulons through *cis*-regulatory motif enrichment analysis, using hg38-based motif databases (hg38 10kbp up 10kbp down and hg38 500bp up 100bp down). Finally, regulon activity was quantified across spatial spots using AUCell.

**CellOracle** [11] was applied using default parameters (https://morris-lab.github.io/CellOracle.documentation/). The workflow involved separate preprocessing of scRNA-seq data using Scanpy and scATAC-seq data using Cicero [105], which identified 7,016 peak-gene connections with co-accessibility scores *≥* 0.8. Transcription factor binding sites (TFBS) were mapped using the default gimmemotifs collection [106]. To construct the GRN, links with statistical significance (*P <* 10^*−*3^) were first retained, from which the top 2,000 links per region were selected based on absolute coefficient weights.

#### 4.6.3 Evaluation of TF recovery performance

To evaluate each method’s ability to identify key regulatory TFs, we conducted a data-driven TF recovery analysis. A reference ranking of TFs was established based on their regulatory impact, assessed by the magnitude of expression differences across spatial domains in the simulated multi-omics dataset (Wilcoxon rank-sum test; *P <* 0.05, log_2_fold-change *>* 2.5). A TF was considered successfully “recovered” by a method if it was predicted to regulate more than 10 target genes. Method performance was quantified by plotting the cumulative number of recovered TFs against their positions in the reference ranking.

#### 4.6.3 Evaluation of predicted TF–peak interactions

The accuracy of predicted TF–peak interactions was evaluated using a ground-truth set of experimentally validated binding sites from the UniBind database [27, 28], which provides a comprehensive map of direct TF–DNA interactions uniformly derived from thousands of ChIP-seq datasets.

To enable a fair comparison, we first identified the TFs shared between each method’s predictions and those cataloged in UniBind. Next, a unified consensus peak set was generated from the simulated dataset using the SnapATAC2 pipeline [83]. Predicted TF–peak interactions from each method, as well as the UniBind reference interactions, were mapped onto this consensus peak set. For each TF, this process yielded a binary interaction vector, where each element indicated the presence (1) or absence (0) of a predicted or validated link to a consensus peak.

Performance was assessed for each TF by comparing its predicted and ground-truth vectors to determine TP, FP, and FN, from which precision and recall were calculated. These were then combined into the F1 score, i.e., the harmonic mean of the two:

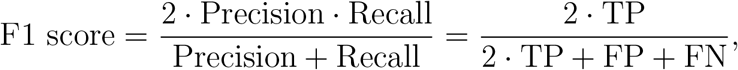

where TP, FP, and FN denote the numbers of true positives, false positives, and false negatives, respectively, with the UniBind interactions serving as the reference ground-truth. This evaluation produced a distribution of F1 scores across TFs for each method. To determine whether STARNet significantly outperformed competing approaches, we performed two-tailed independent two-sample t-tests comparing STARNet’s F1 score distribution to those of the other methods. PySCENIC was excluded from this benchmark, as it does not explicitly infer TF–peak interactions.

#### 4.6.5 Evaluation of predicted TF-target interactions

The accuracy of predicted TF–target interactions was evaluated using the KnockTF2.0 database (https://bio.liclab.net/KnockTFv2/), which contains a high-confidence, tissue- and cell-type-specific set of TF–target interactions derived from TF knockdown and knockout experiments [107].

To perform this benchmark, we first identified the set of TFs for each method that overlap with those included in KnockTF2.0. For each overlapping TF, we treated target prediction as a binary classification task, comparing the method’s predicted targets to experimentally validated targets from the database. Prediction performance was quantified using the F1 score, computed individually for each TF. This procedure yielded a distribution of F1 scores per method, reflecting its overall accuracy across multiple TFs. To assess whether STARNet significantly outperformed other methods, we performed two-tailed independent two-sample t-tests comparing STARNet’s F1 score distribution with those of other approaches.

#### 4.6.6 Evaluation of predicted peak-target links

To assess the quality of inferred peak–target links, we compiled a ground-truth set of 3D chromatin interactions using cell-type-matched Hi-C data from the ENCODE project. Following established protocols [10], Hi-C contact matrices were processed with Juicer Tools (v2.20.00), and interaction scores were extracted at 5 kb resolution and SCALE-normalized. High-confidence interactions, defined as contacts with a score greater than 10, were retained if they connected a genomic bin overlapping a consensus peak to another bin overlapping a transcription start site (TSS, ±1,000 bp).

To ensure consistency across methods, all predicted peaks were mapped to a unified consensus peak set. For each target gene, we then computed an F1 score by comparing the predicted set of linked peaks to the ground-truth interactions defined by the filtered Hi-C contacts. This resulted in a distribution of per-target F1 scores for each method. To evaluate whether STARNet significantly outperformed other approaches, two-tailed independent two-sample t-tests were conducted between the F1 score distribution of STARNet and those of each competing method. pySCENIC was excluded from this benchmark as it does not infer peak–target interactions, and CellOracle was omitted due to the sparsity of its predictions (averaging only 1.5 peaks per target), which precluded robust comparison.

#### 4.6.7 Spatial domain recovery based on inferred GRNs

The accuracy of predicted GRNs was evaluated based on their ability to distinguish known domains within a spatial transcriptomic dataset. For each method, GRN activity scores across all spots were computed using the AUCell algorithm [9, 103], which computes the AUC for the enrichment of each GRN’s target genes in a gene list ranked by expression for each spot. This yielded a spot-by-GRN activity score matrix for each method. To quantitatively assess the alignment between predicted GRN activities and ground-truth domain annotations, we applied PCA to each AUC matrix, followed by hierarchical clustering on the first two principal components. The resulting spot clusters were compared to the pre-annotated domain labels using the ARI, providing a metric of clustering accuracy. In parallel, qualitative assessment was performed by visualizing the AUC matrices with UMAP [108], enabling visual inspection of domain separation.

#### 4.6.8 Robustness evaluation of GRN inference under varying cellular heterogeneity

To systematically benchmark the robustness of GRN inference methods, we utilized the previously described spatial multi-omics dataset simulated from four ENCODE reference cell lines. To introduce varying degrees of cellular heterogeneity, we generated three versions of the dataset, each with a different level of domain purity. Specifically, the proportion of the major cell type within each of the four simulated domains was set to 76%, 88%, and 94%, respectively. By applying each GRN inference method to all three versions, each with five replicates using different random seeds to ensure statistical robustness, we quantitatively evaluated their performance stability under increasing levels of domain purity.

#### 4.6.9 Evaluation of spatially resolved GRN inference on real spatial multi-omics data

We further assessed the performance of the benchmarked methods on a spatially resolved GRN inference task using a P22 mouse brain spatial RNA-ATAC-seq dataset [14]. The evaluation consisted of two key analyses. First, we conducted a TF recovery analysis, as previously described in the benchmark on the simulated dataset, to evaluate whether each method could identify TFs with differential expression across spatial domains. Second, to assess the biological validity of the inferred GRNs, we performed a focused evaluation using a curated panel of TFs with well-established roles in mouse brain development, including Sox10, Olig2, Olig1, Sox4, Sox2, Mef2c, Dlx1, Pax6, Neurod2, and Foxp1 [30,31]. We then examined whether each method successfully recovered these literature-supported TFs, where methods with superior performance are expected to identify a larger subset of these key regulators.

### 4.7 The GO biological process enrichment analysis

GO biological process enrichment analysis was performed using the OmicVerse (v1.7.1) package [109] with default parameters. GO terms were restricted to the “biological processes” category. Enrichment results with false discovery rates (FDRs) less than 0.05, adjusted using the Benjamini-Hochberg procedure, were considered significant.

## 5 Data availability

All datasets used in this study are publicly available. Bulk RNA-seq, ATAC-seq, and Hi-C data for the human cell lines MCF-7, HepG2, GM12878, and K562 were obtained from the ENCODE Project (https://www.encodeproject.org/) under accession numbers: ENCFF136ANW, ENCFF772EFK, ENCFF420JTA (MCF-7); ENCFF660EXG, ENCFF239RGZ, ENCFF020DPP (HepG2); ENCFF626GVO, ENCFF415FEC, ENCFF053VBX (GM12878); ENCFF833WFD, ENCFF512VEZ, ENCFF080DPJ (K562). Spatial RNA-seq and ATAC-seq data for the P21 and P22 mouse brains and human hippocampus were obtained from the Gene Expression Omnibus under accession number GSE205055. Spatial multi-omics data of the mouse embryo were obtained under accession number GSE263333. Annotated single-cell RNA-seq reference data for the mouse brain and embryo were obtained from the Tabula Muris Senis (http://tabula-muris-senis.ds.czbiohub.org/) and the Mouse Organogenesis Cell Atlas (http://atlas.gs.washington.edu/mouse-rna), respectively. Genome annotation (GENCODE v47) was retrieved from GENCODE (https://ftp.ebi.ac.uk/pub/databases/gencode/Gencode_human/release_47/gencode.v47.annotation.gtf.gz), and the GRCh38 genome index for Cell Ranger ARC was obtained from 10x Genomics (https://cf.10xgenomics.com/supp/cell-arc/refdata-cellranger-arc-GRCh38-2020-A-2.0.0.tar.gz).

## 6 Code availability

The source code of STARNet is available on GitHub https://github.com/DBinary/STARNet, and the detailed tutorial is available at https://STARNet_tutorials.readthedocs.io.

## Acknowledgments

We thank Drs. Xi Xi and Xuegong Zhang for providing Chip-seq data, which used in the comprehensive benchmarking of GRN inference. We are grateful for Drs. Fangzhou Xiao, Zitong Jerry Wang, and Jian Yang for their insightful discussions.

## Author contribution statement

L.H., S.Z., X.Z., Y.L., H.Y., and J.Z. designed and developed the method. H.Y. and J.Z. supervised and conceived the project. L.H., S.Z., X.Z., Y.L, and M.Y. designed and implemented the STARNet algorithms, analyzed the data, and performed the performance evaluation task. S.Z., Y.M., H.L., and P.W. helped with the data analysis and discussed the results. H.G., P.L., and S.M. contributed to the method benchmarking. L.C. and Y.X. contributed to the GWAS analyses. L.H., S.Z., X.Z., Y.L., H.Y., and J.Z. wrote the manuscript with help from other authors. All authors have reviewed and approved the final manuscript.

