## supplementary figure for "STARNet enables spatially resolved inference of gene regulatory networks from spatial multi-omics data"

**1 Department of Artificial Intelligence, School of Engineering, Westlake University, Hangzhou, China.**

**2 School of Life Sciences, Westlake University, Hangzhou, China.**

**3 New Cornerstone Science Laboratory, School of Life Sciences, Westlake University, Hangzhou, China.**

**4 School of Computer Science, Beijing University of Technology, Beijing, China.**

**5 School of Advanced Engineering, University of Science and Technology Beijing, Beijing, China.**

**6 Department of Automation, Tsinghua University, Beijing, China.**

**7 AI Thrust, Information Hub, The Hong Kong University of Science and Technology (Guangzhou), Guangzhou, China.**

**8 Department of Biomedical Informatics and Data Science, Yale School of Medicine, New Haven, CT, USA.**

**9 Zhongguancun Institute of Artificial Intelligence, Beijing, China.**

**10 School of Mathematical Sciences and LPMC, Nankai University, Tianjin, China.**

**11 Sheffield Institute for Translational Neuroscience, University of Sheffield, Sheffield, UK.**

**12 School of Life Science and Technology, Xidian University, Xi'an, China.**

**13 Center for Interdisciplinary Studies, School of Science, Westlake University, Hangzhou, China.**

**† These authors contributed equally: Lei Hu, Shichen Zhang, Xuting Zhang, Yihai Luo.**

#### This document contains:

**- Supplementary Figures 1-16**

### Supplementary Figures

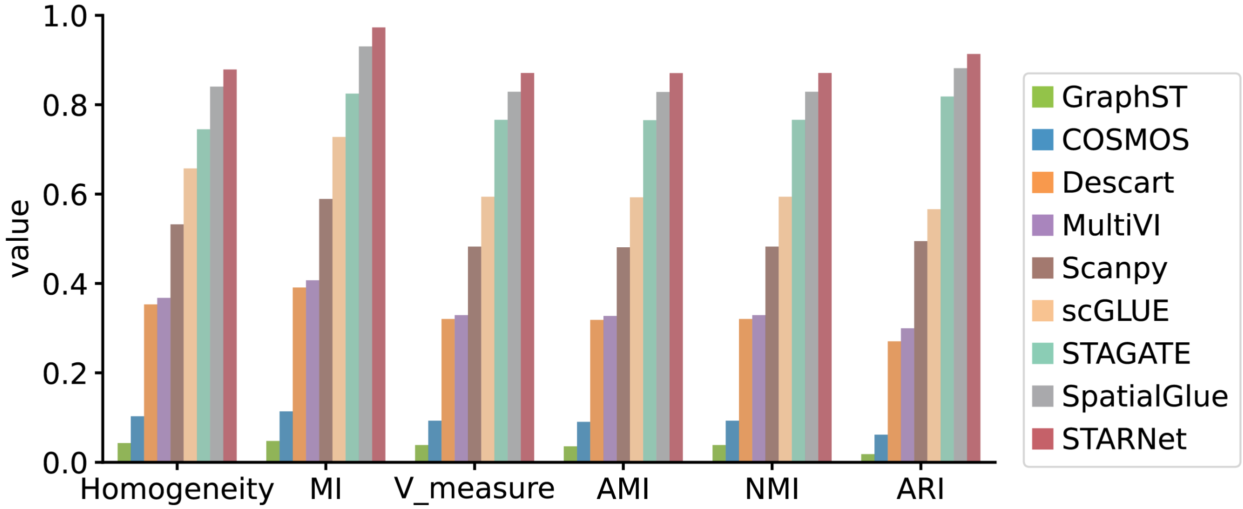


**Fig. S1.** **Bar plots showing the performance of various methods when applied to domain identification on the simulated spatial multi-omics dataset (Fig. 2a), evaluated using supervised integration metrics: homogeneity, mutual information (MI), V_measure, adjusted mutual information (AMI), normalized mutual information (NMI), and adjusted Rand index (ARI).**

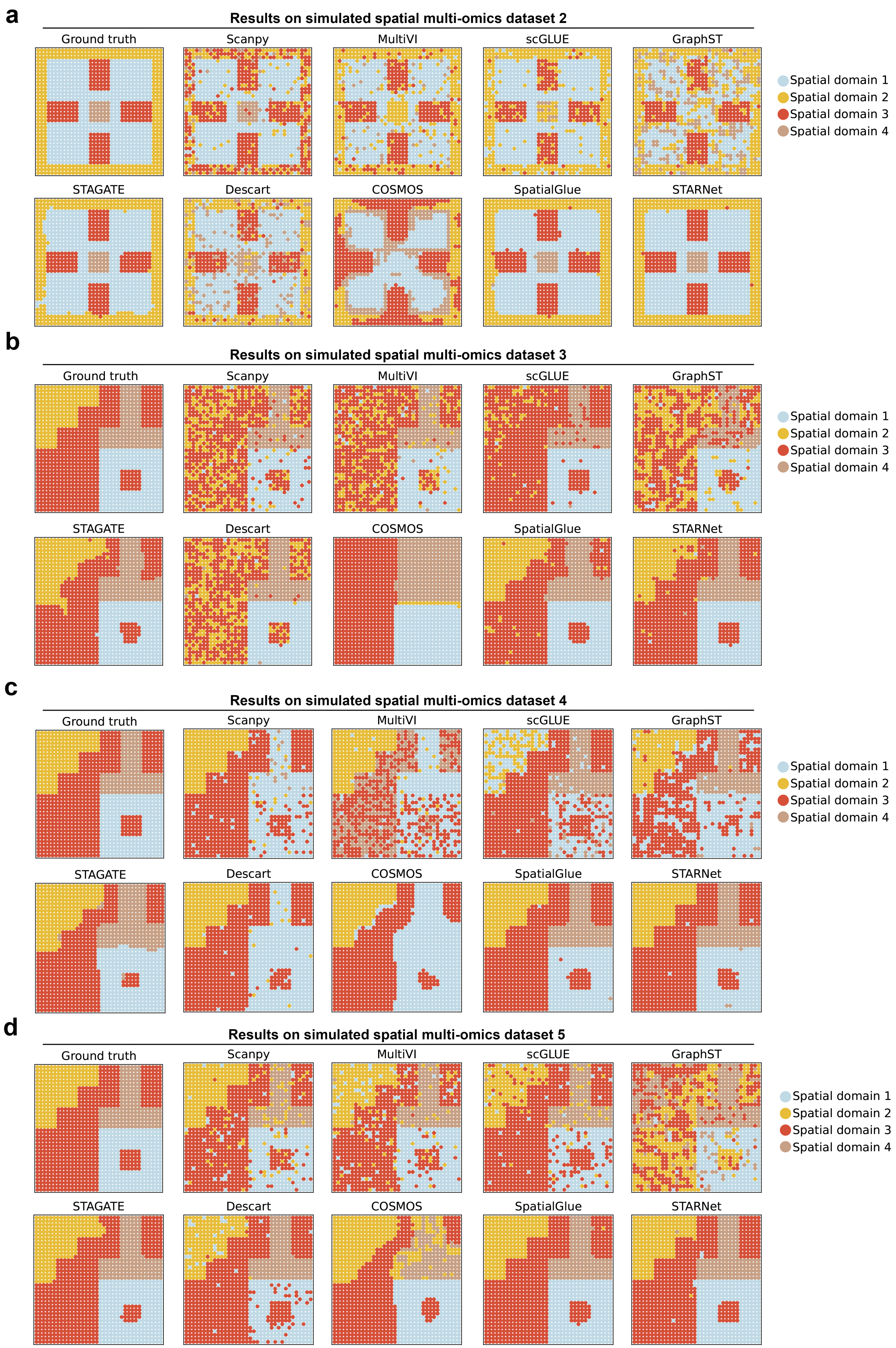
a
Results on simulated spatial multi-omics dataset 2
Ground truth
Scanpy
MultiVI
scGLUE
GraphST
STAGATE
Descart
COSMOS
SpatialGlue
STARNet
Spatial domain 1
Spatial domain 2
Spatial domain 3
Spatial domain 4
b
Results on simulated spatial multi-omics dataset 3
Ground truth
Scanpy
MultiVI
scGLUE
GraphST
STAGATE
Descart
COSMOS
SpatialGlue
STARNet
Spatial domain 1
Spatial domain 2
Spatial domain 3
Spatial domain 4
c
Results on simulated spatial multi-omics dataset 4
Ground truth
Scanpy
MultiVI
scGLUE
GraphST
STAGATE
Descart
COSMOS
SpatialGlue
STARNet
Spatial domain 1
Spatial domain 2
Spatial domain 3
Spatial domain 4
d
Results on simulated spatial multi-omics dataset 5
Ground truth
Scanpy
MultiVI
scGLUE
GraphST
STAGATE
Descart
COSMOS
SpatialGlue
STARNet
Spatial domain 1
Spatial domain 2
Spatial domain 3
Spatial domain 4

**Fig. S2**

**Fig. S2. Benchmarking STARNet alongside eight other methods for spatial domain identification on simulated spatial multi-omics datasets 2-5. Panels a-d,** Spatial mapping of spots in datasets 2-5, respectively, colored by ground-truth domain labels and by inferred domains from Scanpy, DESCART, Cosmos, MultiVI, GraphST, SpatialGlue, scGLUE, STAGATE and STARNet.

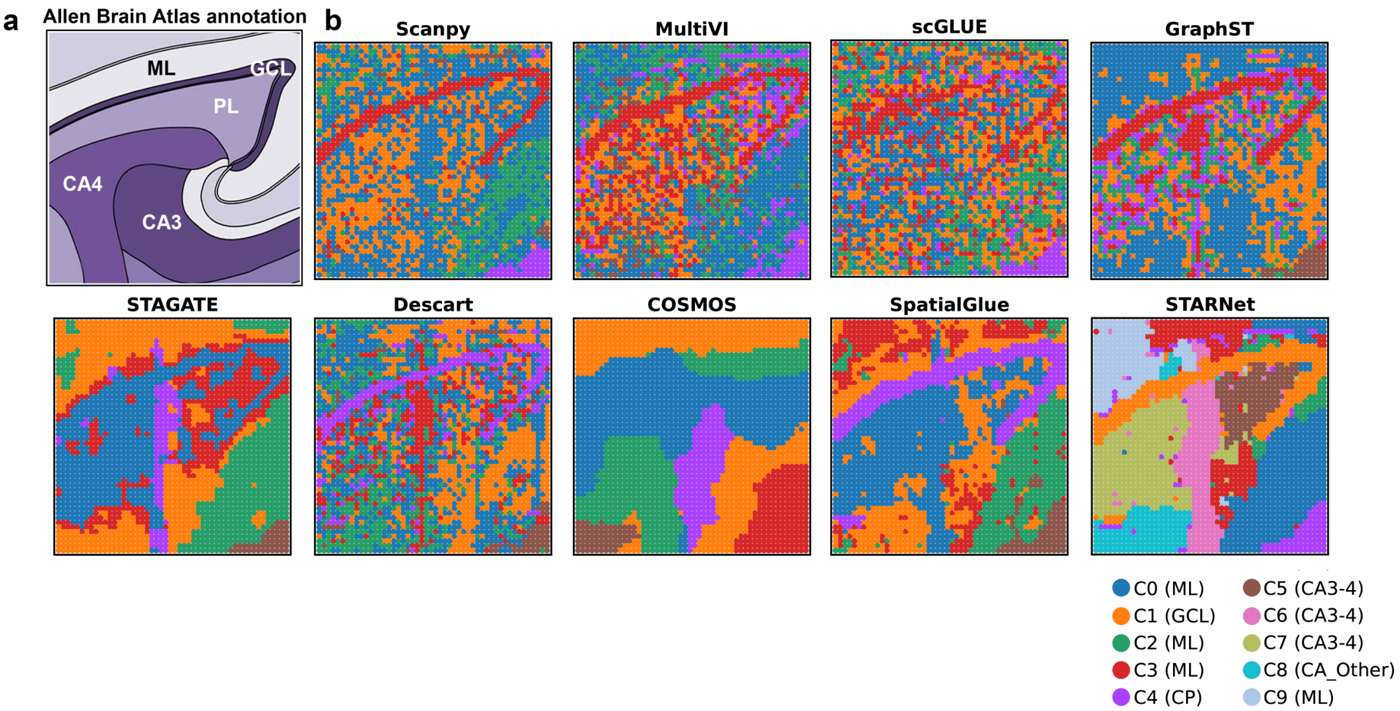
a
Allen Brain Atlas annotation
ML
GCL
PL
CA4
CA3
b
Scanpy
MultiVI
scGLUE
GraphST
STAGATE
Descart
COSMOS
SpatialGlue
STARNet
C0 (ML)
C1 (GCL)
C2 (ML)
C3 (ML)
C4 (CP)
C5 (CA3-4)
C6 (CA3-4)
C7 (CA3-4)
C8 (CA_Other)
C9 (ML)

**Fig. S3**

**Fig. S3. Comparison of spatial domain identification by STARNet and eight other methods (Scanpy, MultiVI, scGLUE, STAGATE, SpatialGlue, GraphST, COSMOS, and DESCART) on a human hippocampus spatial multi-omics dataset.** **a,** Annotated regional reference of the same hippocampus section from the Allen Human Brain Atlas. **b,** Spatial domains identified by each method for this tissue section. Abbreviations: CA3-4, cornu ammonis 3-4; ML, molecular layer; GCL, granule cell layer; CP, choroid plexus; CA Other, remaining cornu ammonis regions.

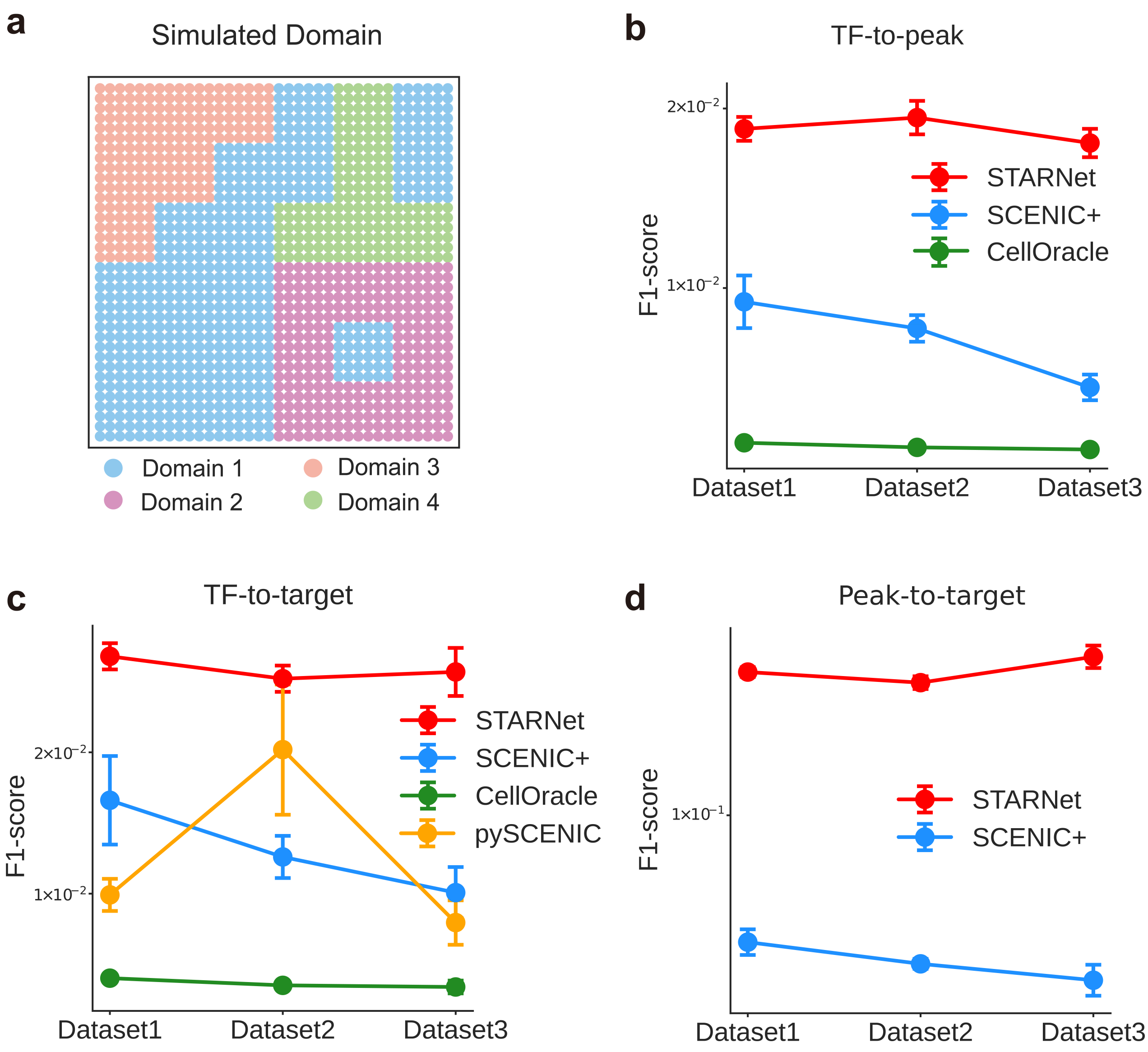
a
Simulated Domain
Domain 1
Domain 3
Domain 2
Domain 4
b
TF-to-peak
F1-score
2×10⁻²
1×10⁻²
STARNet
SCENIC+
CellOracle
Dataset1
Dataset2
Dataset3
c
TF-to-target
F1-score
2×10⁻²
1×10⁻²
STARNet
SCENIC+
CellOracle
pySCENIC
Dataset1
Dataset2
Dataset3
d
Peak-to-target
F1-score
1×10⁻¹
STARNet
SCENIC+
Dataset1
Dataset2
Dataset3

**Fig. S4**

**Fig. S4. Supplementary information for the performance comparison of GRN inference methods on simulated spatial multi-omics datasets. a,** Spatial distribution of the simulated spatial multi-omics datasets used to benchmark GRN inference methods. Each dot represents a spatial spot, color-coded by its simulated domain label. **b-d,** Line plots of mean F1 scores or (b) TF-to-peak, (c) TF-to-target, and (d) peak-to-target predictions on three additional simulated datasets. All datasets share the same spatial domain structure as in (a) but differ in the dominant cell-type proportion per spot (0.94, 0.88, and 0.76) to model increasing spatial complexity. Each point on the line represents the mean F1 score, with error bars indicating the standard deviations.

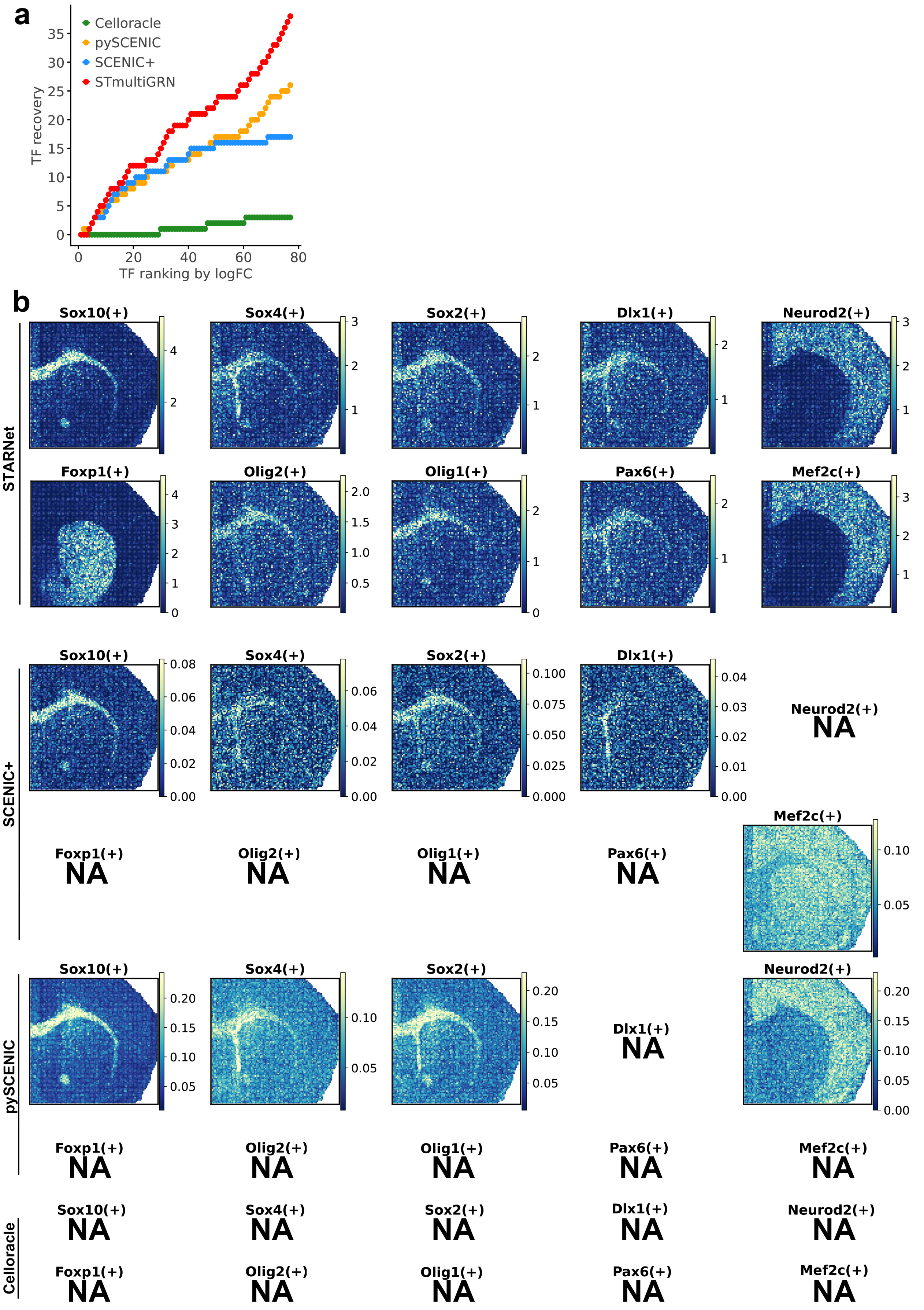
a
Celloracle
pySCENIC
SCENIC+
STmultiGRN
TF recovery
0
5
10
15
20
25
30
35
0
20
40
60
80
TF ranking by logFC
b
STARNet
Sox10(+)
Sox4(+)
Sox2(+)
Dlx1(+)
Neurod2(+)
Foxp1(+)
Olig2(+)
Olig1(+)
Pax6(+)
Mef2c(+)
SCENIC+
Sox10(+)
Sox4(+)
Sox2(+)
Dlx1(+)
Neurod2(+)
NA
Foxp1(+)
NA
Olig2(+)
NA
Olig1(+)
NA
Pax6(+)
NA
Mef2c(+)
pySCENIC
Sox10(+)
Sox4(+)
Sox2(+)
Dlx1(+)
NA
Neurod2(+)
Foxp1(+)
NA
Olig2(+)
NA
Olig1(+)
NA
Pax6(+)
NA
Mef2c(+)
NA
Celloracle
Sox10(+)
NA
Sox4(+)
NA
Sox2(+)
NA
Dlx1(+)
NA
Neurod2(+)
NA
Foxp1(+)
NA
Olig2(+)
NA
Olig1(+)
NA
Pax6(+)
NA
Mef2c(+)
NA

**Fig. S5**

**Fig. S5. Comparison of GRN inference results for key brain-development TFs by STARNet, pySCENIC, SCENIC+, and CellOracle on the P22 mouse brain data. a,** Cumulative recovery curves of TFs across different methods on the postnatal day 22 (P22) mouse brain data. TFs were ranked in descending order of $\log_2$ fold-change, based on differential expression analysis between spatial domains. **b,** Comparison of spatial significance or activity scores across four methods for GRNs of key brain-development TFs. The GRNs of TFs Sox10, Olig2, Olig1, Sox4, Sox2, Mef2c, Dlx1, Pax6, Neurod2 and Foxp1 were evaluated. For STARNet, spatial significance scores are displayed as $-\log_{10} P$ derived from spatial enrichment tests. For SCENIC+, pySCENIC, and CellOracle, GRN activity scores computed via AUCell are shown, reflecting the cumulative expression of each TF and its target genes in spots. "NA" indicates that the network was not inferred by the corresponding method (see Methods).

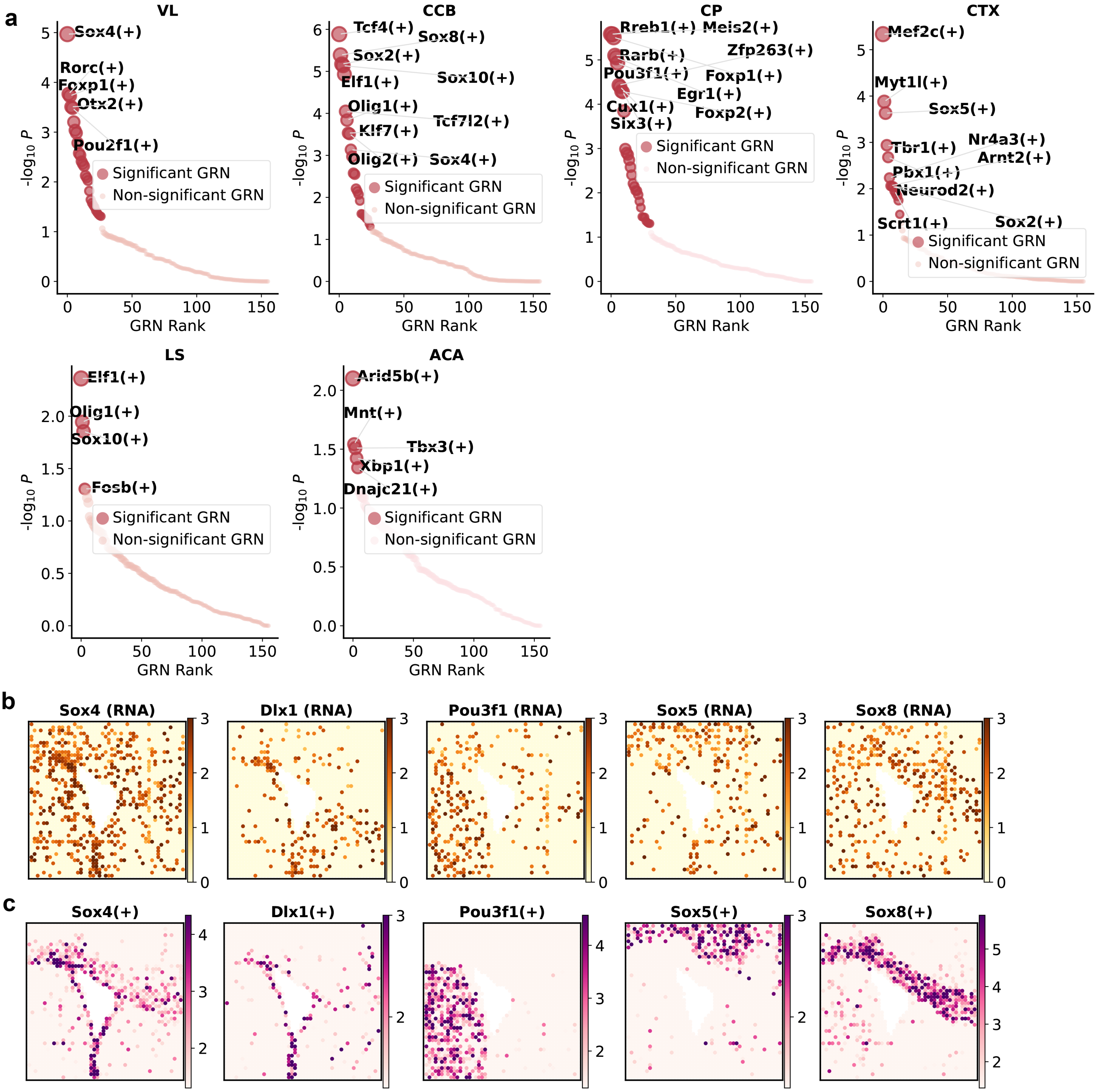

a
VL
Sox4(+)
Rorc(+)
Foxp1(+)
Otx2(+)
Pou2f1(+)
CCB
Tcf4(+)
Sox8(+)
Sox2(+)
Sox10(+)
Elf1(+)
Olig1(+)
Tcf7l2(+)
Klf7(+)
Olig2(+)
Sox4(+)
CP
Rreb1(+)
Meis2(+)
Zfp263(+)
Rarb(+)
Pou3f1(+)
Foxp1(+)
Egr1(+)
Cux1(+)
Foxp2(+)
Six3(+)
CTX
Mef2c(+)
Myt1l(+)
Sox5(+)
Nr4a3(+)
Tbr1(+)
Arnt2(+)
Pbx1(+)
Neurod2(+)
Scrt1(+)
Sox2(+)
LS
Elf1(+)
Olig1(+)
Sox10(+)
Fosb(+)
ACA
Arid5b(+)
Mnt(+)
Tbx3(+)
Xbp1(+)
Dnajc21(+)
Significant GRN
Non-significant GRN
-log10 P
GRN Rank
b
Sox4 (RNA)
Dlx1 (RNA)
Pou3f1 (RNA)
Sox5 (RNA)
Sox8 (RNA)
c
Sox4(+)
Dlx1(+)
Pou3f1(+)
Sox5(+)
Sox8(+)


**Fig. S6**

**Fig. S6. Spatial distribution patterns of GRNs inferred by STARNet across anatomical mouse brain domains.** **a,** Dot plots displaying the aggregated $P$-value of all GRNs across individual spots within each spatial domain. The $P$-value for each GRN within a domain is calculated through the Cauchy combination test, and GRNs achieving $P < 0.05$ are designated as significant. The top 10 significant GRNs are shown (If fewer than 10 significant GRNs are identified, all are displayed). **b,** Normalized expression levels of examples of spatially specific TFs (Dlx1, Sox4, Sox5, Pou3f1, and Sox8). Each dot represents a spot, color-coded by the normalized expression level of the corresponding gene. **c,** Significance scores of the spatially specific GRNs, involving TFs Dlx1, Sox4, Sox5, Pou3f1 and Sox8. Each dot represents a spot, colored by the $-\log_{10} P$. The color scale's minimum is set to $-\log_{10} 0.05$. Abbreviations: ACA, anterior cingulate area; CCB, corpus callosum, body; CP, caudoputamen; CTX, cerebral cortex; LS, lateral septal nucleus; VL, lateral ventricle.

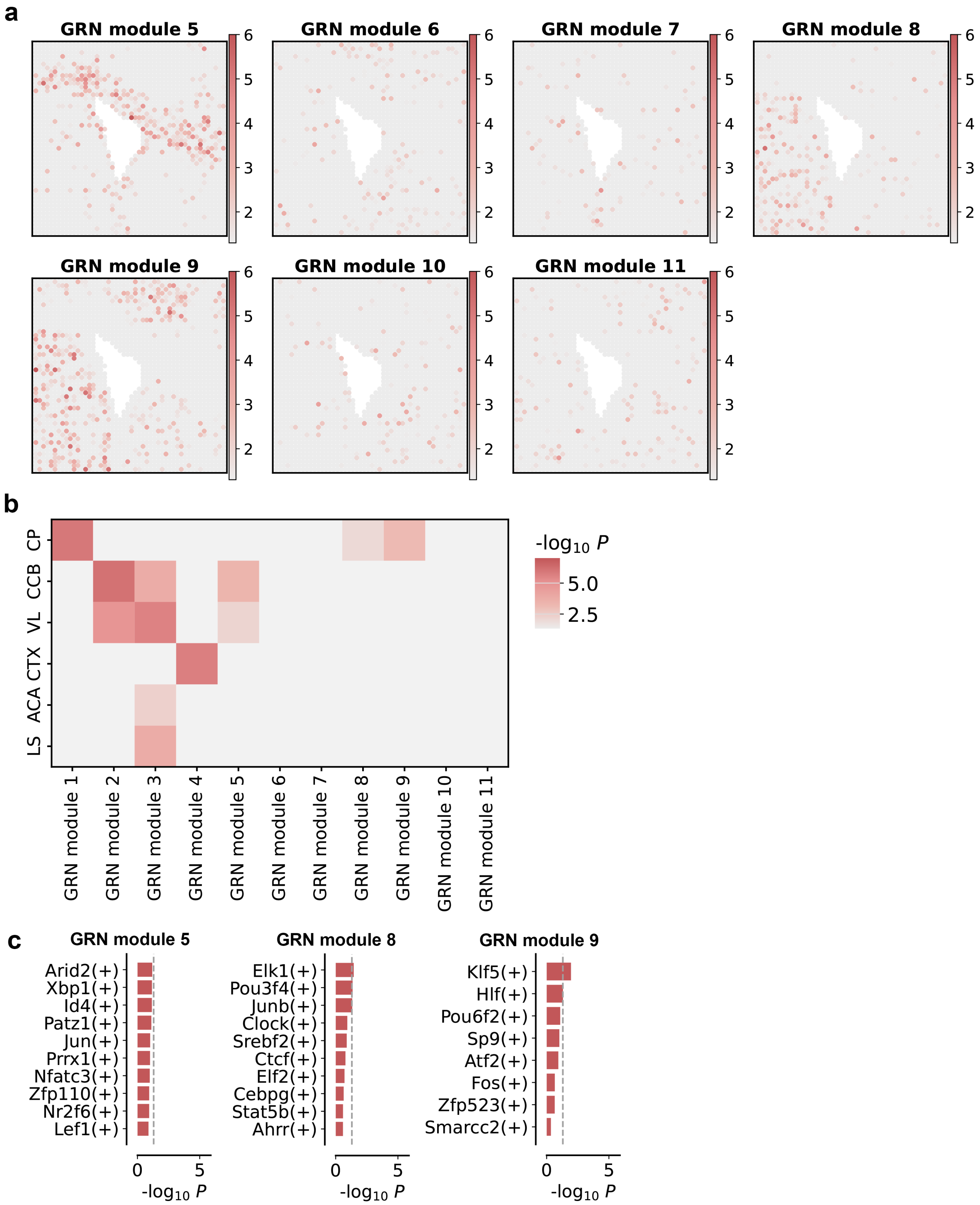
a
GRN module 5
GRN module 6
GRN module 7
GRN module 8
GRN module 9
GRN module 10
GRN module 11
b
CP
CCB
VL
CTX
ACA
LS
-log10 P
5.0
2.5
GRN module 1
GRN module 2
GRN module 3
GRN module 4
GRN module 5
GRN module 6
GRN module 7
GRN module 8
GRN module 9
GRN module 10
GRN module 11
c
GRN module 5
Arid2(+)
Xbp1(+)
Id4(+)
Patz1(+)
Jun(+)
Prrx1(+)
Nfatc3(+)
Zfp110(+)
Nr2f6(+)
Lef1(+)
GRN module 8
Elk1(+)
Pou3f4(+)
Junb(+)
Clock(+)
Srebf2(+)
Ctcf(+)
Elf2(+)
Cebpg(+)
Stat5b(+)
Ahrr(+)
GRN module 9
Klf5(+)
Hlf(+)
Pou6f2(+)
Sp9(+)
Atf2(+)
Fos(+)
Zfp523(+)
Smarcc2(+)
-log10 P

**Fig. S7**

**Fig. S7. Spatial distribution patterns of GRN modules and their associations with anatomical mouse brain domains. a,** Spatial mapping of significance scores of GRN module 5-11, derived by clustering GRN-level significance profiles across all spatial spots using the Leiden algorithm (see Methods). Clustering was performed based on inter-spot similarity in the low-dimensional representation of significance profiles obtained through PCA. Each point corresponds to a spatial spot, colored by the $-\log_{10} P$. The color scale's minimum is set to $-\log_{10} 0.05$, highlighting statistically significant spots ($P < 0.05$). **b,** Heatmap displaying the associations between GRN modules and spatial domains. The aggregated $P$-values for individual GRN modules within the corresponding domains were calculated using the Cauchy combination test (see Methods). Color intensity indicates the statistical significance of each association, represented as $-\log_{10} P$. **c,** Horizontal bar plots displaying the aggregated $P$-values for GRNs in modules 5, 8, and 9, calculated using the Cauchy combination test. GRN Module 5 (showing the top 10 GRNs ranked by $P$-value) is associated with the CCB region, while GRN Modules 8 (showing the top 10 GRNs) and 9 (displaying all 8 GRNs) are both associated with the CP region. A gray dashed line at $-\log_{10} 0.05$ indicates the significance threshold for individual GRNs. Abbreviations: ACA, anterior cingulate area; CCB, corpus callosum, body; CP, caudoputamen; CTX, cerebral cortex; LS, sateral septal nucleus; VL, lateral ventricle.

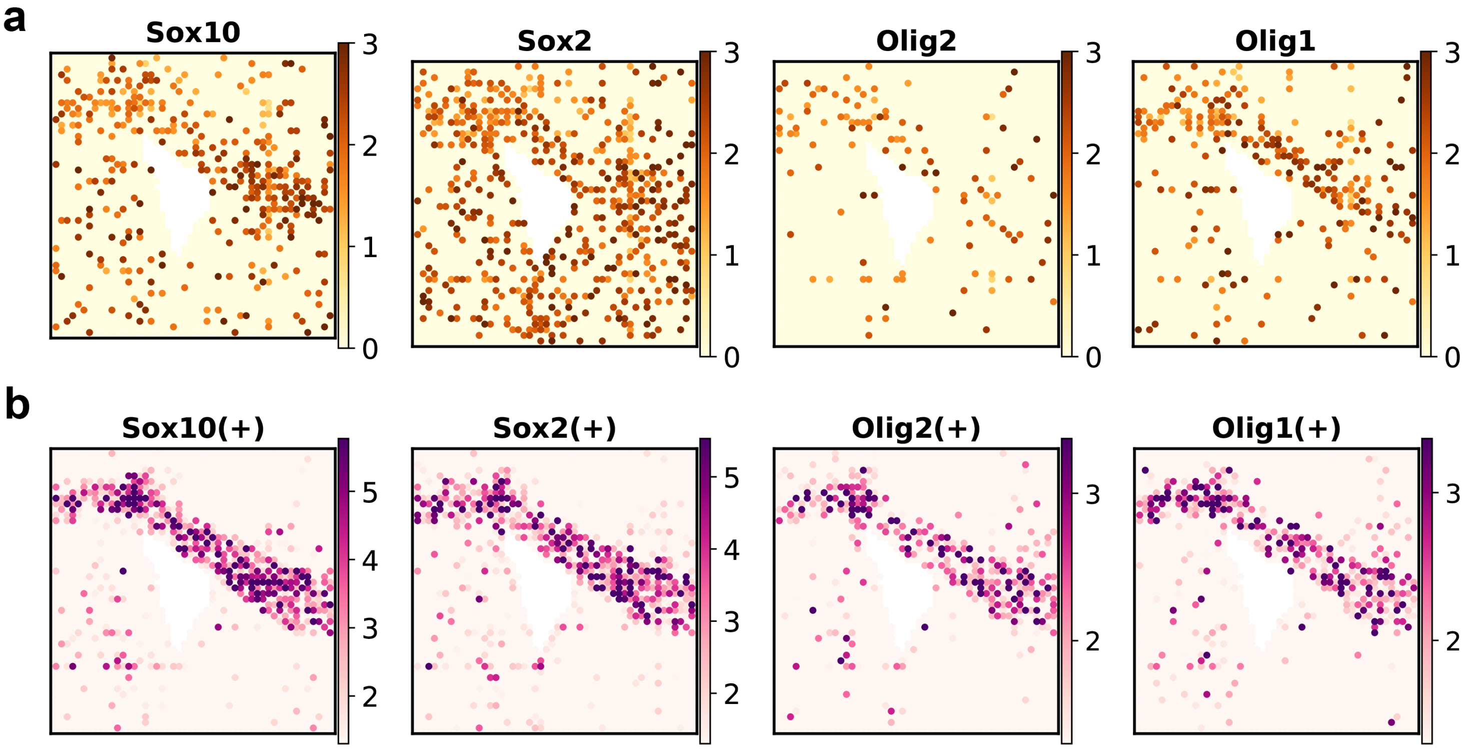


**Fig. S8**

**Fig. S8. Supplementary results for GRNs involving TFs by Sox10, Sox2, Olig2, and Olig1.** **a,** Normalized expression levels of TFs, Sox10, Sox2, Olig2, and Olig1, visualized with color coding based on normalized gene expression levels. **b,** Spatial significance score mapping of the GRNs, involving Sox10, Sox2, Olig2, and Olig1 with colors represented $-\log_{10} P$. The color scale minimum is set to $-\log_{10} 0.05$, highlighting the statistically significant regions.

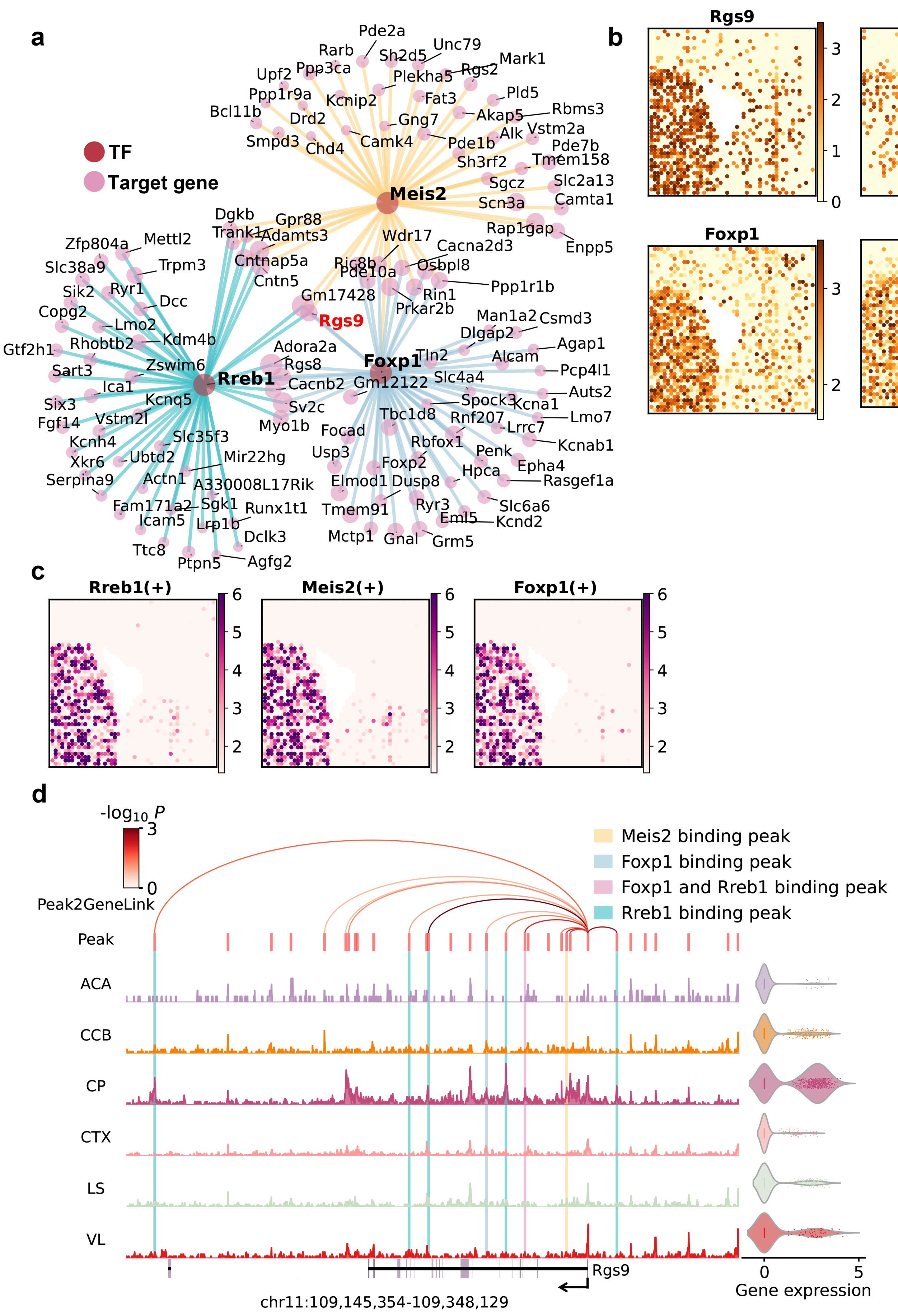
a
TF
Target gene
Meis2
Rreb1
Foxp1
Rgs9
Pde2a
Rarb
Unc79
Sh2d5
Mark1
Upf2
Ppp3ca
Rgs2
Plekha5
Ppp1r9a
Pld5
Kcnip2
Fat3
Bcl11b
Rbms3
Drd2
Akap5
Gng7
Vstm2a
Smpd3
Alk
Camk4
Pde1b
Pde7b
Chd4
Sh3rf2
Tmem158
Sgcz
Slc2a13
Scn3a
Camta1
Dgkb
Gpr88
Rap1gap
Trank1
Adamts3
Enpp5
Zfp804a
Mettl2
Wdr17
Cacna2d3
Cntnap5a
Trpm3
Slc38a9
Ric8b
Osbpl8
Cntn5
Pde10a
Ppp1r1b
Sik2
Ryr1
Dcc
Rin1
Gm17428
Copg2
Prkar2b
Man1a2
Csmd3
Lmo2
Dlgap2
Kdm4b
Agap1
Gtf2h1
Rhobtb2
Adora2a
Tln2
Alcam
Zswim6
Rgs8
Pcp4l1
Sart3
Slc4a4
Cacnb2
Gm12122
Auts2
Ica1
Spock3
Kcna1
Kcnq5
Six3
Sv2c
Lmo7
Vstm2l
Tbc1d8
Rnf207
Fgf14
Myo1b
Focad
Lrrc7
Slc35f3
Kcnh4
Rbfox1
Kcnab1
Mir22hg
Ubtd2
Usp3
Penk
Xkr6
Foxp2
Epha4
Actn1
Hpca
Serpina9
Elmod1
Dusp8
Rasgef1a
A330008L17Rik
Fam171a2
Sgk1
Runx1t1
Ryr3
Slc6a6
Tmem91
Icam5
Lrp1b
Eml5
Kcnd2
Dclk3
Mctp1
Gnal
Grm5
Ttc8
Ptpn5
Agfg2
b
Rgs9
Rreb1
Foxp1
Meis2
c
Rreb1(+)
Meis2(+)
Foxp1(+)
d
-log10 P
Peak2GeneLink
Meis2 binding peak
Foxp1 binding peak
Foxp1 and Rreb1 binding peak
Rreb1 binding peak
Peak
ACA
CCB
CP
CTX
LS
VL
Rgs9
chr11:109,145,354-109,348,129
Gene expression

**Fig. S9**

**Fig. S9. Analysis of the GRNs involving TFs Meis2, Rreb1, and Roxp1 in the Caudoputamen of P21 mouse brain. a,** Visualization of the GRNs involving TFs Rreb1, Meis2, and Foxp2. Nodes represent both the TFs (red) and their top 50 target genes (pink) ranked by their regulatory scores (see Methods). Node size indicates the regulatory score of each target gene, and edge color distinguishes individual regulatory relationships with different TFs. **b,** Normalized expression levels of the involved TFs Meis2, Rreb1, Foxp1, and Rgs9, visualized with color coding based on normalized gene expression levels. **c,** Spatial significance score mapping of the selected GRNs, involving TFs Meis2, Rreb1, and Foxp1, with colors representing $-\log_{10} P$. The color scale minimum is set to $-\log_{10} 0.05$, highlighting the statistically significant regions. **d,** Chromatin accessibility profiles of genomic regions spanning the Rgs9 locus across different spatial domains. The Peak2GeneLinks are visualized as arcs, color-coded by the statistical significance of their corresponding correlations. Peak2GeneLinks were inferred based on correlations between chromatin accessibility peaks and gene expression across the metacells of all spatial spots (see Methods). The predicted binding peaks of Meis2, Foxp1, and Rreb1, identified using the SnapATAC2 toolkit, are highlighted with semi-transparent boxes. A violin plot on the right shows the normalized expression levels of Rgs9 across different spatial domains.

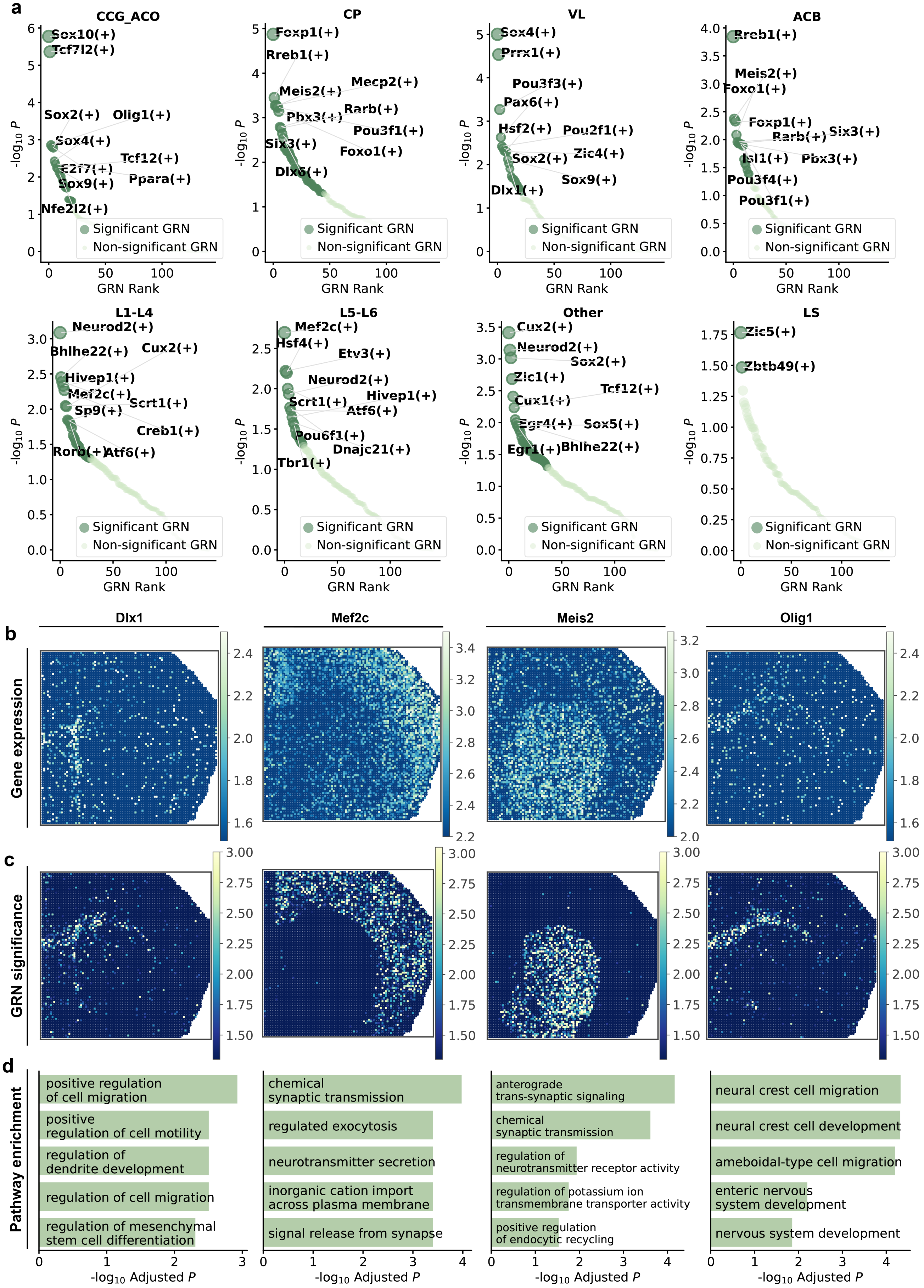
a
CCG_ACO
CP
VL
ACB
Sox10(+)
Tcf7l2(+)
Sox2(+)
Olig1(+)
Sox4(+)
Tcf12(+)
E2f7(+)
Ppara(+)
Sox9(+)
Nfe2l2(+)
Foxp1(+)
Rreb1(+)
Meis2(+)
Mecp2(+)
Pbx3(+)
Rarb(+)
Pou3f1(+)
Six3(+)
Foxo1(+)
Dlx6(+)
Prrx1(+)
Pou3f3(+)
Pax6(+)
Hsf2(+)
Pou2f1(+)
Zic4(+)
Dlx1(+)
Isl1(+)
Pou3f4(+)
L1-L4
L5-L6
Other
LS
Neurod2(+)
Bhlhe22(+)
Cux2(+)
Hivep1(+)
Mef2c(+)
Scrt1(+)
Sp9(+)
Creb1(+)
Rorb(+)
Atf6(+)
Hsf4(+)
Etv3(+)
Pou6f1(+)
Dnajc21(+)
Tbr1(+)
Zic1(+)
Cux1(+)
Egr4(+)
Sox5(+)
Egr1(+)
Zic5(+)
Zbtb49(+)
Significant GRN
Non-significant GRN
GRN Rank
-log10 P
b
Dlx1
Mef2c
Meis2
Olig1
Gene expression
c
GRN significance
d
Pathway enrichment
positive regulation of cell migration
positive regulation of cell motility
regulation of dendrite development
regulation of cell migration
regulation of mesenchymal stem cell differentiation
chemical synaptic transmission
regulated exocytosis
neurotransmitter secretion
inorganic cation import across plasma membrane
signal release from synapse
anterograde trans-synaptic signaling
regulation of neurotransmitter receptor activity
regulation of potassium ion transmembrane transporter activity
positive regulation of endocytic recycling
neural crest cell migration
neural crest cell development
ameboidal-type cell migration
enteric nervous system development
nervous system development
-log10 Adjusted P

**Fig. S10**

**Fig. S10. Spatial patterns of GRNs inferred by STARNet across anatomical regions of the P22 mouse brain. a,** Dot plots of aggregated $P$-values for all GRNs across individual spots within each spatial domain. The $P$-values of each GRN were calculated through the Cauchy combination test, and GRNs with $P < 0.05$ are considered significant. The top 10 significant GRNs are shown (If fewer than 10 meet the threshold, all are shown). **b,** Normalized expression levels of the selected TFs, including Dlx1, Mef2c, Meis2, and Olig1, visualized with color coding based on normalized gene expression levels. **c,** Spatial significance score mapping of the GRNs involving the same TFs, with colors representing the $-\log_{10} P$. The color scale minimum is set to $-\log_{10} 0.05$, highlighting the statistically significant regions. **d,** GO term enrichment analysis for target genes associated with each GRN with $P$-values determined by the hypergeometric test.

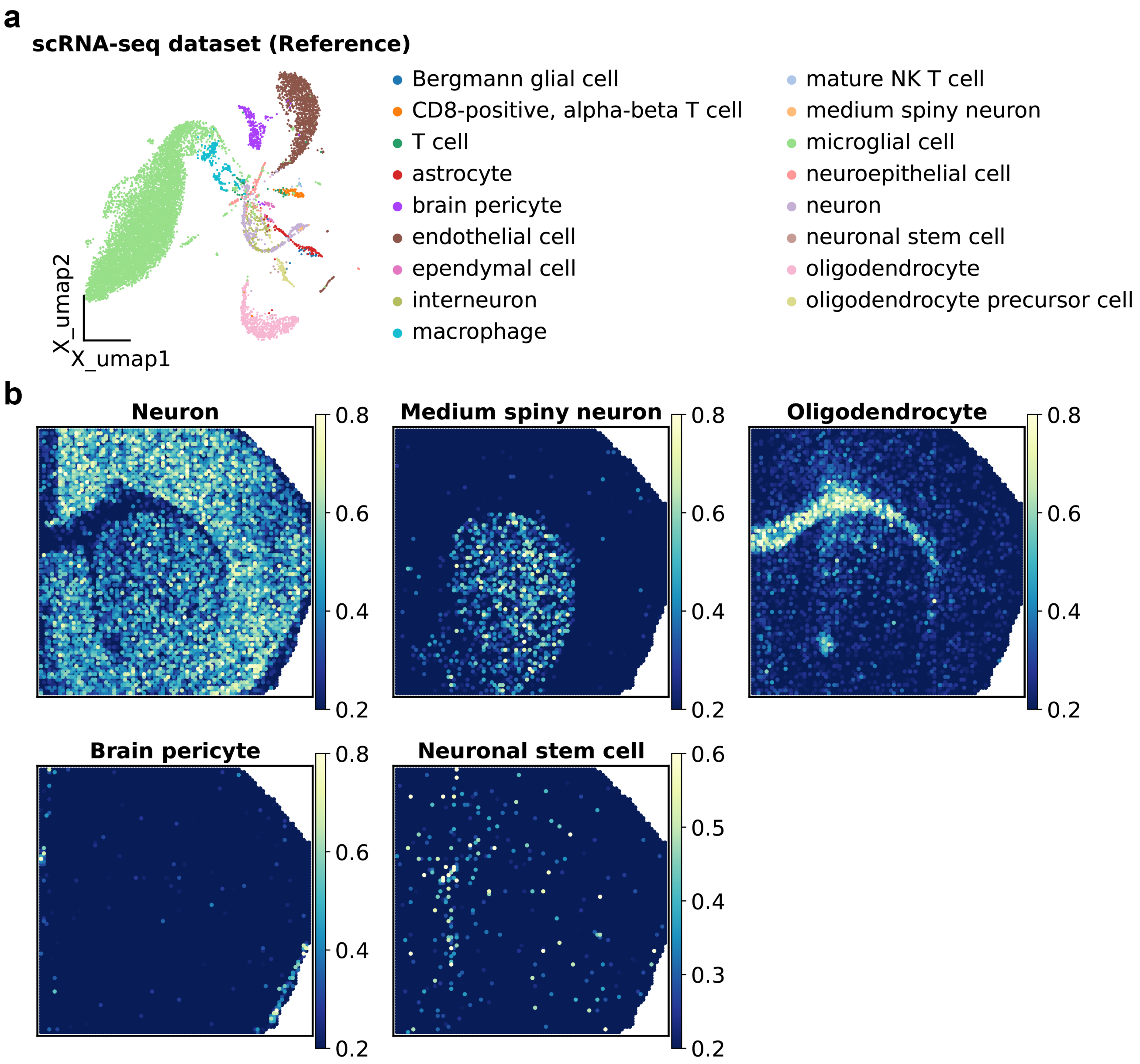
a
scRNA-seq dataset (Reference)
Bergmann glial cell
CD8-positive, alpha-beta T cell
T cell
astrocyte
brain pericyte
endothelial cell
ependymal cell
interneuron
macrophage
mature NK T cell
medium spiny neuron
microglial cell
neuroepithelial cell
neuron
neuronal stem cell
oligodendrocyte
oligodendrocyte precursor cell
X_umap2
X_umap1
b
Neuron
Medium spiny neuron
Oligodendrocyte
Brain pericyte
Neuronal stem cell
0.8
0.6
0.5
0.4
0.3
0.2

**Fig. S11**

**Fig. S11. Mapping scRNA-seq cell types onto the spatial multi-omics data of P22 mouse brain with Tangram. a,** UMAP visualization of the scRNA-seq reference dataset used in the deconvolution analysis of spatial multi-omics data, with colors indicating the cell-type labels from the source study. **b,** Probabilistic mapping of scRNA-seq cell types onto the spatial multi-omics data using the Tangram algorithm, visualized by color-coded cell-type probabilities (see Methods). Tangram is a deep learning-based method that integrates single-cell and spatial transcriptomic data to generate a probability matrix representing the likely spatial location of each cell.

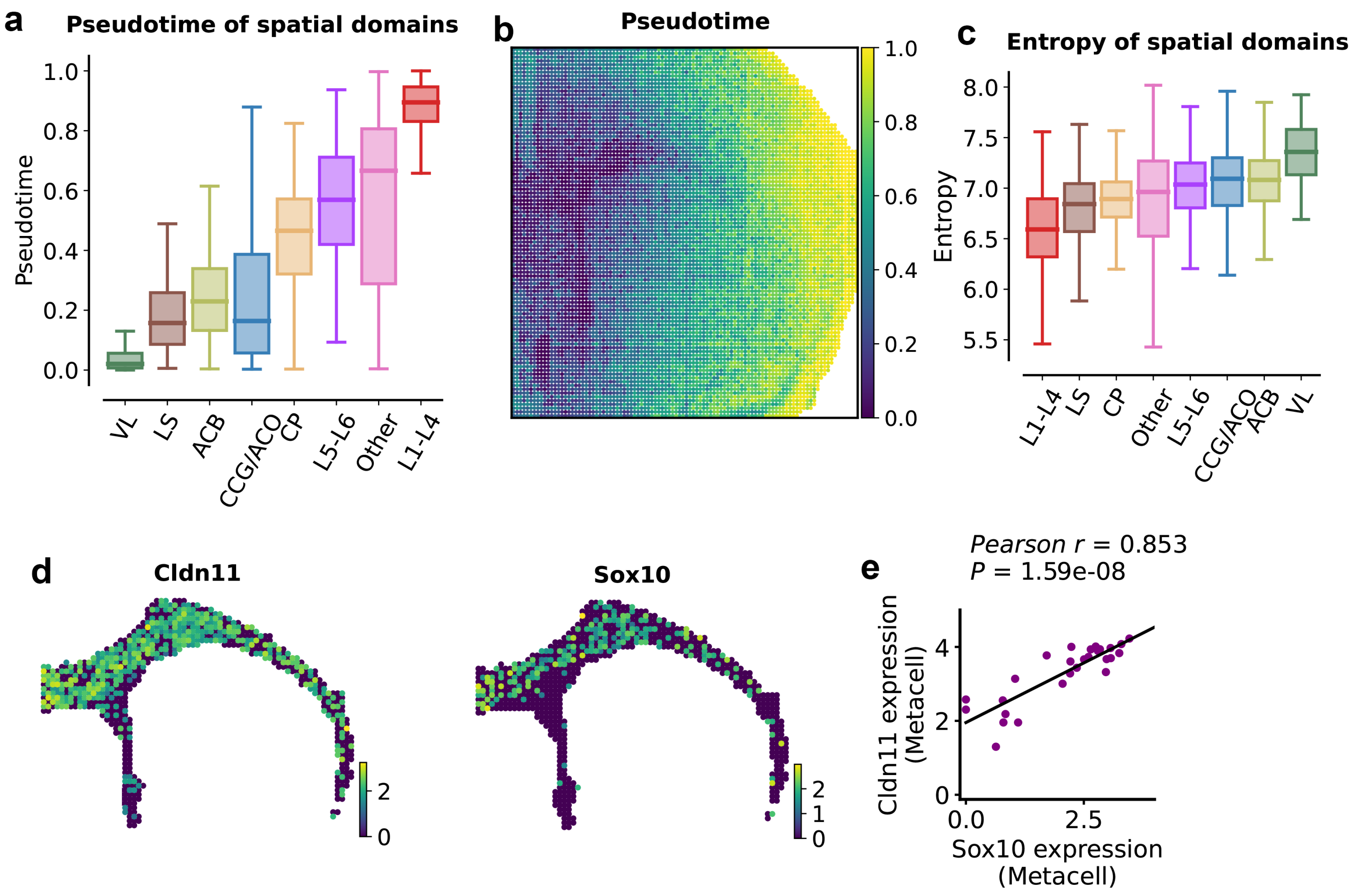
a
Pseudotime of spatial domains
Pseudotime
VL
LS
ACB
CCG/ACO
CP
L5-L6
Other
L1-L4
b
Pseudotime
c
Entropy of spatial domains
Entropy
d
Cldn11
Sox10
e
Pearson r = 0.853
P = 1.59e-08
Cldn11 expression (Metacell)
Sox10 expression (Metacell)

**Fig. S12**

**Fig. S12. Supplementary results for the spatiotemporal analysis of P22 mouse brain.** **a,** Boxplots of pseudotime values grouped by spatial domains, ordered by median pseudotime. Boxes show the median (center line) and interquartile range (box edges), with colors indicating spatial domains. Pseudotime was calculated per spot using spaTrack (see Methods), which applies optimal transport theory to reconstruct developmental trajectories from spatial transcriptomic data. **b,** Spatial distribution of pseudotime values, with color gradient from dark (low pseudotime) to high (high pseudotime). **c,** Boxplot of entropy values grouped by spatial domains ordered by median entropy. Shannon entropy was computed for each spot (see Methods) to quantify functional uncertainty and differentiation potential. Boxes represent the median (center line) and interquartile range (box edges), with colors indicating spatial domains. **d,** Normalized expression levels of Sox10 and Cldn11 in CCG and VL regions, visualized using a color scale. **e,** Scatter plot showing the correlation between Sox10 and Cldn11 expression in CCG and VL regions. Each point represents a metacell derived from STARNet-generated spot embeddings (see Methods). Pearson correlation coefficient and its associated two-sided $P$-value are reported.

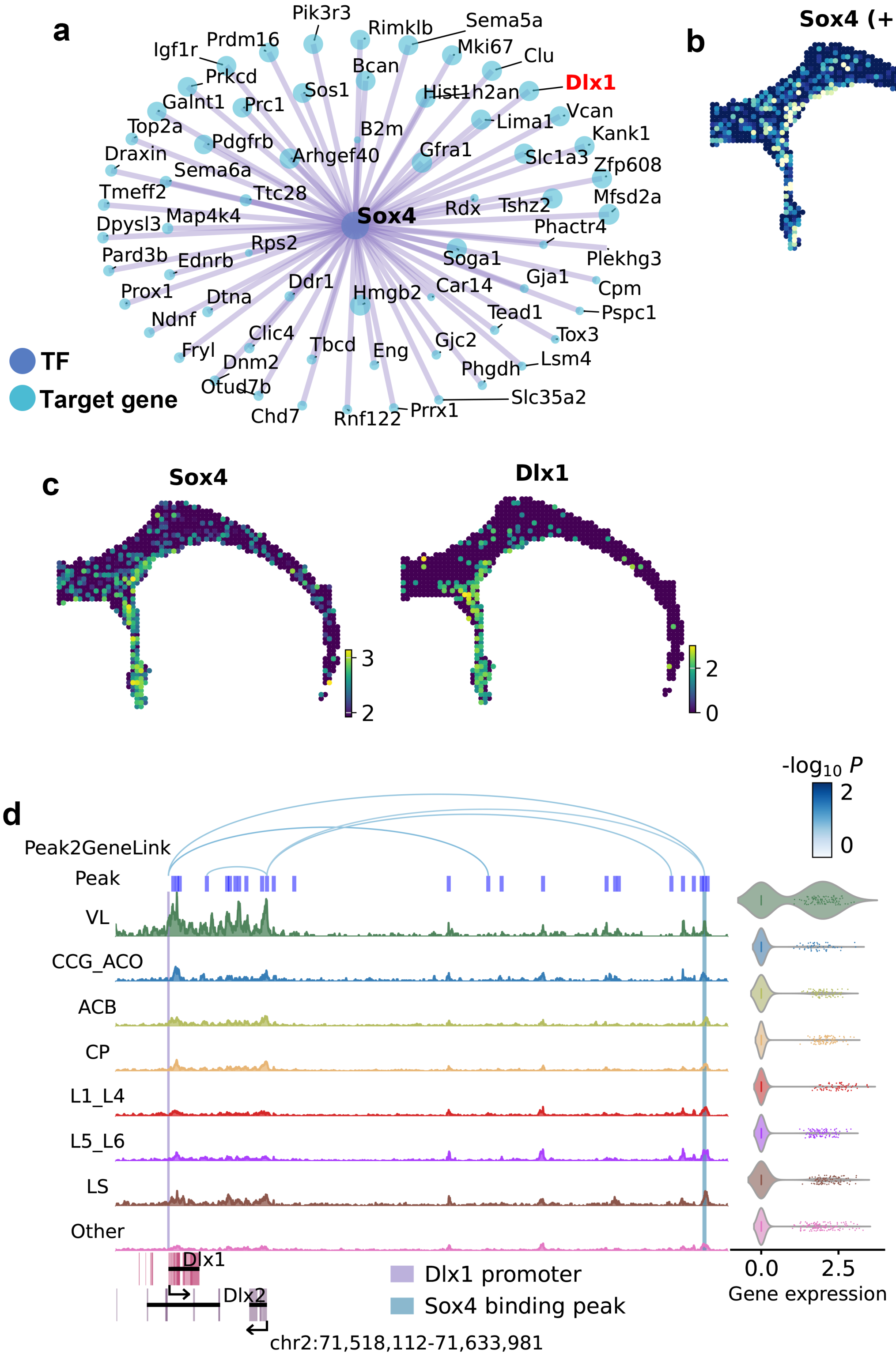
a
TF
Target gene
Sox4
Pik3r3
Rimklb
Sema5a
Igf1r
Prdm16
Mki67
Clu
Bcan
Dlx1
Prkcd
Sos1
Hist1h2an
Galnt1
Prc1
Vcan
Lima1
Top2a
B2m
Kank1
Pdgfrb
Gfra1
Draxin
Arhgef40
Slc1a3
Zfp608
Sema6a
Tmeff2
Ttc28
Mfsd2a
Tshz2
Rdx
Map4k4
Dpysl3
Phactr4
Rps2
Pard3b
Ednrb
Soga1
Plekhg3
Gja1
Ddr1
Car14
Cpm
Prox1
Dtna
Hmgb2
Tead1
Pspc1
Ndnf
Clic4
Tox3
Gjc2
Fryl
Tbcd
Eng
Lsm4
Dnm2
Phgdh
Otud7b
Slc35a2
Prrx1
Chd7
Rnf122
b
Sox4 (+)
4
2
c
Sox4
3
2
Dlx1
2
0
d
-log10 P
2
0
Peak2GeneLink
Peak
VL
CCG_ACO
ACB
CP
L1_L4
L5_L6
LS
Other
Dlx1
Dlx2
Dlx1 promoter
Sox4 binding peak
chr2:71,518,112-71,633,981
0.0
2.5
Gene expression

**Fig. S13**

**Fig. S13. Analysis of the Sox4-regulated GRN in NSCs from the P22 mouse brain. a,** Visualization of the GRN regulated by Sox4. Nodes represent both the TF (deep blue) and its target genes (light blue), with node size proportional to regulatory strength (see Methods). Dlx1 is highlighted in red for downstream analysis. **b,** Spatial mapping of significance scores for the GRN regulated by Sox4, with colors representing $-\log_{10} P$. The color scale lower bound is set to $-\log_{10} 0.05$, highlighting only regions with $P \leq 0.05$. **c,** Normalized expression of selected genes, including Sox4 and Dlx1, visualized using a color scale. **d,** Chromatin accessibility profiles of genomic regions spanning the Dlx1 locus across different spatial domains. Peak2GeneLinks are shown as arches, color-coded by correlation significance (see Methods). Predicted TF-binding peaks are highlighted with shaded boxes. A violin plot on the right demonstrates the normalized expression levels of Dlx1 across different spatial domains.

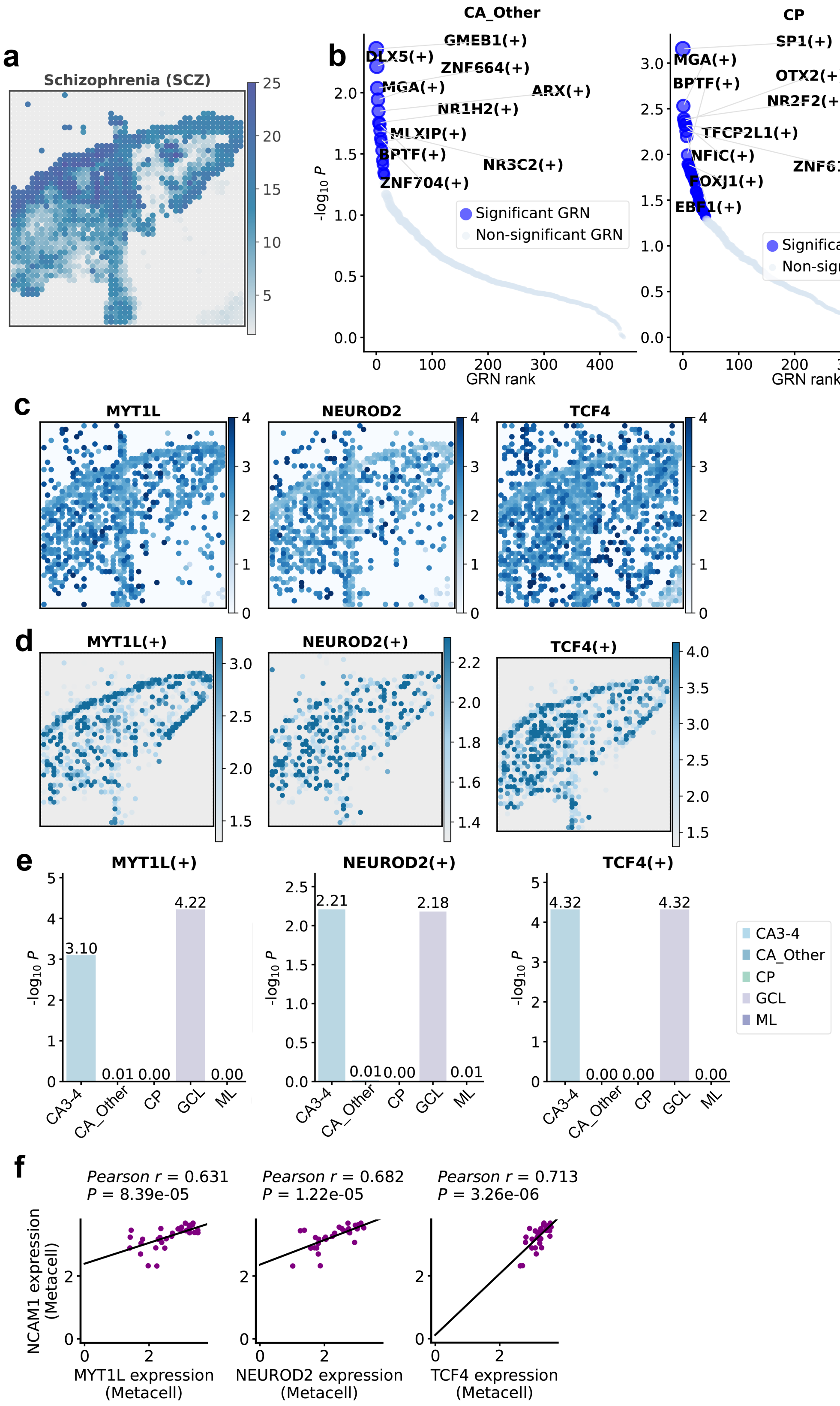
a
Schizophrenia (SCZ)
b
CA_Other
CP
ML
GMEB1(+)
DLX5(+)
ZNF664(+)
MGA(+)
ARX(+)
NR1H2(+)
MLXIP(+)
BPTF(+)
NR3C2(+)
ZNF704(+)
SP1(+)
OTX2(+)
NR2F2(+)
TFCP2L1(+)
NFIC(+)
ZNF619(+)
FOXJ1(+)
EBF1(+)
ZFY(+)
ZIC1(+)
CREBZF(+)
SATB1(+)
ZNF84(+)
ZNF615(+)
ZNF341(+)
SP3(+)
FOXG1(+)
LMX1A(+)
HIC2(+)
ZNF430(+)
Significant GRN
Non-significant GRN
$-\log_{10} P$
GRN rank
c
MYT1L
NEUROD2
TCF4
d
MYT1L(+)
NEUROD2(+)
TCF4(+)
e
CA3-4
CA_Other
CP
GCL
ML
f
Pearson r = 0.631
P = 8.39e-05
Pearson r = 0.682
P = 1.22e-05
Pearson r = 0.713
P = 3.26e-06
NCAM1 expression (Metacell)
MYT1L expression (Metacell)
NEUROD2 expression (Metacell)
TCF4 expression (Metacell)

**Fig. S14**

**Fig. S14. Spatial patterns of GRNs inferred by STARNet across different spatial domains of the human hippocampus.** **a,** Spatial map of schizophrenia association significance computed by gsMap on a human hippocampal section. Colors indicate $-\log_{10} P$. **b,** Dot plot showing aggregated $P$-values (calculated using the Cauchy combination test) for all GRNs across individual spots in CA_Other, ML, and CP regions. GRNs with $P < 0.05$ are designated significant, and the top ten significant GRNs are labeled. **c,** Spatial maps of normalized expression for selected genes (MYT1L, NEUROD2, and TCF4), color-coded by their expression levels. **d,** Spatial maps of significance scores for STARNet-inferred GRNs regulated by selected TFs (MYT1L, NEUROD2, and TCF4), color-coded by $-\log_{10} P$. The color-scale lower bound is set to $-\log_{10} 0.05$, highlighting regions with $P < 0.05$. **e,** Bar plots showing aggregated $P$-values (calculated using the Cauchy combination test) for the STARNet-inferred GRNs regulated by selected TFs (MYT1L, NEUROD2, and TCF4) across spatial domains. **f,** Scatter plots showing correlations between NCAM1 expression and that of MYT1L, NEUROD2, and TCF4, across metacells in the human hippocampal section. Metacells were defined by STARNet-derived spot embeddings (see Methods). $P$-values were calculated using the two-sided t-test. Abbreviations: CA3-4, cornu ammonis 3-4; ML, molecular layer; GCL, granule cell layer; CP, choroid plexus; CA_Other, remaining cornu ammonis regions.

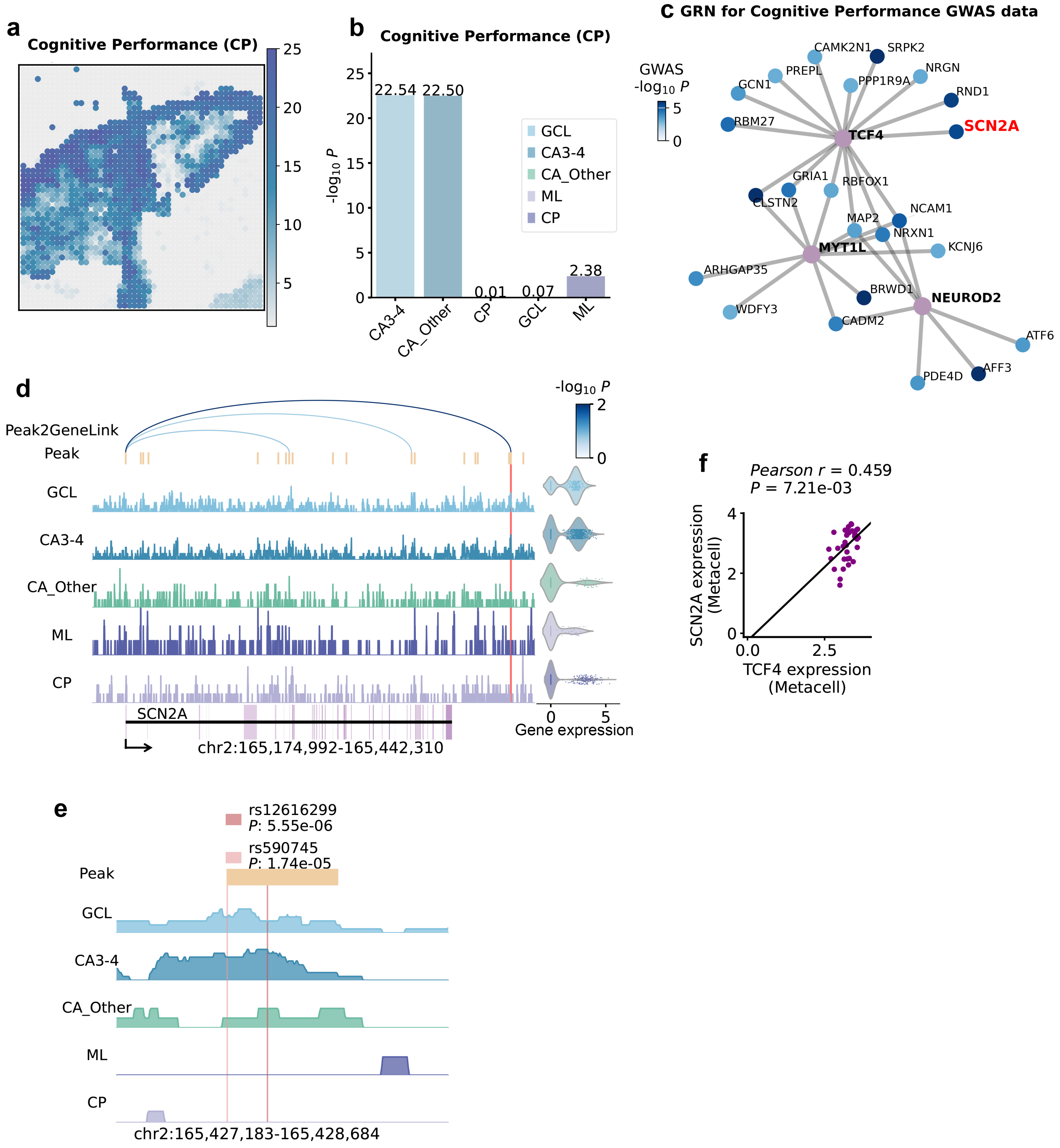
a
Cognitive Performance (CP)
25
20
15
10
5
b
Cognitive Performance (CP)
-log10 P
22.54 22.50
0.01 0.07 2.38
GCL
CA3-4
CA_Other
ML
CP
CA3-4
CA_Other
CP
GCL
ML
c GRN for Cognitive Performance GWAS data
GWAS
-log10 P
CAMK2N1
SRPK2
PREPL
NRGN
GCN1
PPP1R9A
RND1
RBM27
TCF4
SCN2A
GRIA1
RBFOX1
CLSTN2
NCAM1
MAP2
NRXN1
MYT1L
KCNJ6
ARHGAP35
BRWD1
NEUROD2
WDFY3
CADM2
ATF6
PDE4D
AFF3
d
Peak2GeneLink
Peak
GCL
CA3-4
CA_Other
ML
CP
SCN2A
chr2:165,174,992-165,442,310
-log10 P
Gene expression
f
Pearson r = 0.459
P = 7.21e-03
SCN2A expression
(Metacell)
TCF4 expression
(Metacell)
0.0
2.5
e
rs12616299
P: 5.55e-06
rs590745
P: 1.74e-05
Peak
GCL
CA3-4
CA_Other
ML
CP
chr2:165,427,183-165,428,684

**Fig. S15**

**Fig. S15. Mapping cognitive performance GWAS associations onto STARNet-derived spatially specific GRNs in the human hippocampus.** **a,** Spatial map of cognitive performance association significance across a human hippocampal section, computed with gsMap. Colors represent $-\log_{10} P$. **b,** Bar plot showing aggregated $P$-values (calculated using the Cauchy combination test) for associations between cognitive performance and different spatial domains. **c,** Visualization of the STARNet-inferred GRNs regulated by selected TFs (TCF4, NEUROD2, and MYT1L). Only target genes harboring SNPs with $P < 0.01$ in GWAS summary statistics are shown. Node colors indicate the $P$-values of their associated SNPs. The gene SCN2A is highlighted in red for downstream analysis. **d,** Chromatin accessibility profiles of the genomic regions encompassing SCN2A across spatial domains. Peak2GeneLinks were inferred by computing correlations between chromatin accessibility peaks and SCN2A expression across metacells from all spatial spots (see Methods), and are visualized as colored arches, with color indicating the correlation significance. The predicted TCF4 binding peak, overlapping significant SNPs rs12616299 and rs590745 and identified using the SnapATAC2 toolkit, is highlighted by a vertical red bar. The violin plot (right) shows normalized SCN2A expression across spatial domains. **e,** Chromatin accessibility profiles across spatial domains for the genomic region encompassing the predicted TCF4 binding peak (yellow), with SNPs rs12616299 and rs590745 highlighted by vertical semi-transparent bars. **f,** Scatter plot showing the correlation between SCN2A and TCF4 expression across the human hippocampal section. Each point represents a metacell defined based on the STARNet-derived spot embeddings (see Methods). $P$-values were calculated using a two-sided t-test.

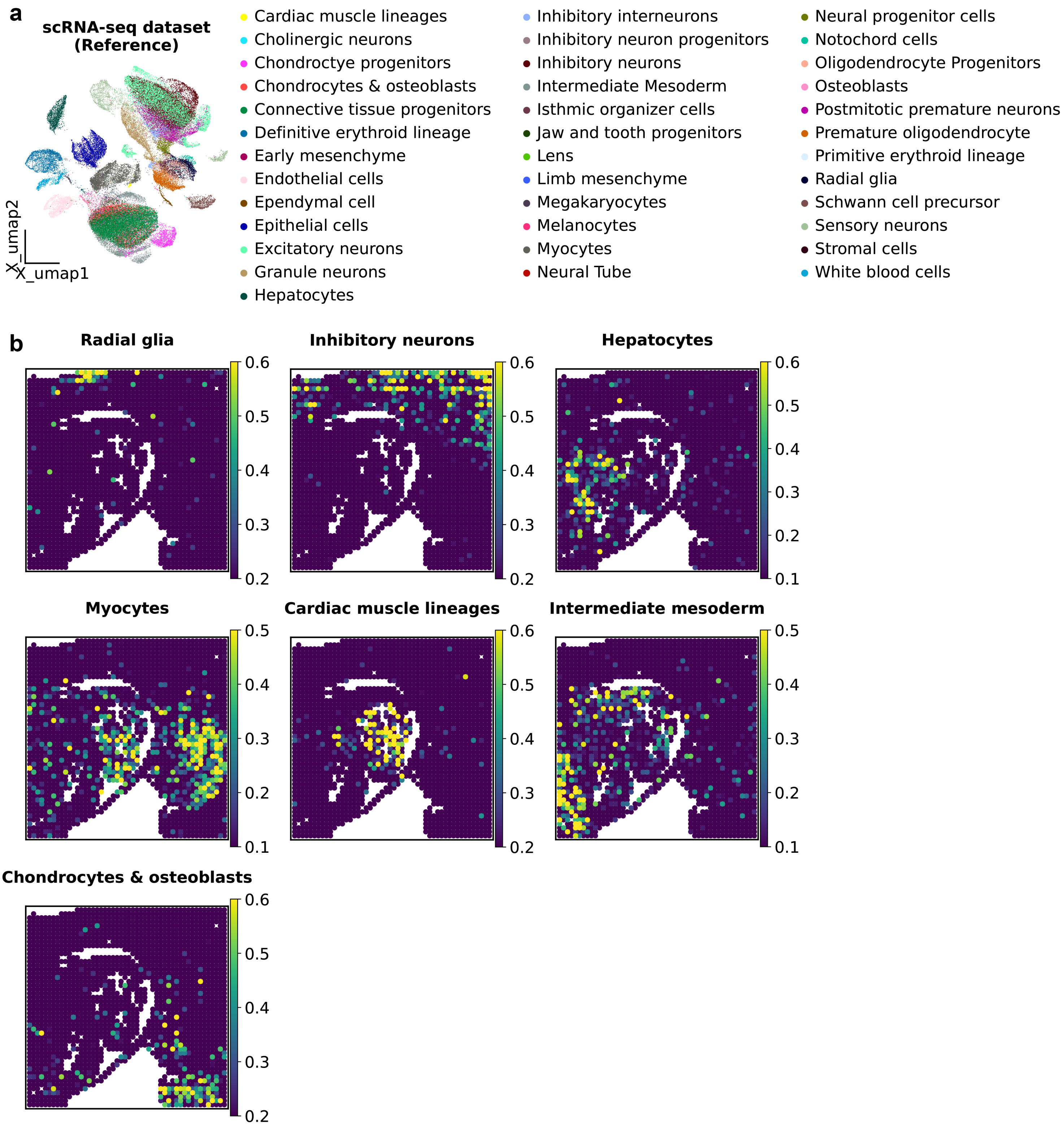
a
scRNA-seq dataset (Reference)
X_umap2
X_umap1
Cardiac muscle lineages
Cholinergic neurons
Chondroctye progenitors
Chondrocytes & osteoblasts
Connective tissue progenitors
Definitive erythroid lineage
Early mesenchyme
Endothelial cells
Ependymal cell
Epithelial cells
Excitatory neurons
Granule neurons
Hepatocytes
Inhibitory interneurons
Inhibitory neuron progenitors
Inhibitory neurons
Intermediate Mesoderm
Isthmic organizer cells
Jaw and tooth progenitors
Lens
Limb mesenchyme
Megakaryocytes
Melanocytes
Myocytes
Neural Tube
Neural progenitor cells
Notochord cells
Oligodendrocyte Progenitors
Osteoblasts
Postmitotic premature neurons
Premature oligodendrocyte
Primitive erythroid lineage
Radial glia
Schwann cell precursor
Sensory neurons
Stromal cells
White blood cells
b
Radial glia
Inhibitory neurons
Hepatocytes
Myocytes
Cardiac muscle lineages
Intermediate mesoderm
Chondrocytes & osteoblasts

**Fig. S16**

**Fig. S16. Projection of scRNA-seq-defined cell types onto spatial multi-omics profiles of the mouse embryo.** **a,** UMAP visualization of the scRNA-seq reference dataset used for deconvolution of the spatial multi-omics data (see Methods), labels indicate cell types as annotated in the source study. **b,** Probabilistic mapping of scRNA-seq cell types onto spatial multi-omics data using Tangram, a deep learning-based method that aligns single-cell transcriptomes with spatial transcriptomic data (see Methods). Color denotes the probability of each cell type being located at each spatial position.
